# Programmable amide bond formation using *de novo* ligases

**DOI:** 10.64898/2026.11.08.765178

**Authors:** Saman Salike, Timo Flügel, Maximilian Gantz, Stanislav S. Moroz, Andrew C Hunt, Henry Jones, Abhishek Singh, Nathaniel Corley, Kieran Didi, Rafael I Brent, Maxwell W Kazman, Zefei Xu, Christina Savvides, Seth M Woodbury, Belen Sotomayor, Hannah Nguyen, Johnny Mendoza, Alex Kang, Asim K Bera, Xinting Li, Sambashiva Banala, Jasper Butcher, Luke D Lavis, Anthony P Green, Donald Hilvert, David Baker

**Affiliations:** Institute for Protein Design, University of Washington, Seattle, WA, USA; Department of Biochemistry, Institute for Protein Design, University of Washington, Seattle, WA, 98195, USA; Department of Chemical Engineering, University of Washington, Seattle, WA, USA; Max Planck Institute for Terrestrial Microbiology, Marburg, Germany; Manchester Institute of Biotechnology, The University of Manchester, Manchester, UK; Department of Chemistry, The University of Manchester, Manchester, UK; Department of Bioengineering, University of Washington, Seattle, WA, USA; NVIDIA Corporation, Santa Clara, CA, USA; Department of Computer Science, University of Oxford, Oxford, UK; Graduate Program in Molecular Engineering, University of Washington, Seattle, WA, USA; Graduate Program in Biological Physics, Structure and Design, University of Washington, Seattle, WA, USA; Janelia Research Campus, Howard Hughes Medical Institute, Ashburn, VA, USA; Paul G. Allen School of Computer Science and Engineering, University of Washington, Seattle, WA, USA; Laboratory of Organic Chemistry, Department of Chemistry and Applied Biosciences, ETH Zurich, Zurich, Switzerland; Howard Hughes Medical Institute, University of Washington, Seattle, WA, USA

## Abstract

Amide bond formation is of central importance in the chemical industry, underpinning the synthesis of small bioactive molecules, macrocycles, peptides and their conjugates. Conventional methods of amide synthesis rely on stoichiometric coupling agents and costly protecting group manipulations. While biocatalysis can alleviate some of these challenges, enzymes typically operate across a narrow range of substrates and their adaptation to new coupling partners requires extensive engineering. Here we show that de novo amide ligases featuring a cysteine catalytic triad can be programmed computationally to operate across diverse substrate classes with high selectivity. We first designed complementary peptide ligases to selectively functionalise either the N-terminus or a single lysine residue of a polypeptide sequence containing 36 competing lysines. Crystal structures of both ligases show excellent agreement with the designs, with structural modelling suggesting that amidation outcompetes hydrolysis because the catalytically competent geometry forms only upon acceptor binding. We then extended our approach to the synthesis of small bioactive molecules by designing stereoselective ligases that mediate a key amide bonding forming step in the synthesis of the preventative HIV therapeutic lenacapavir. These results establish de novo enzyme design as a programmable platform for achieving selective amide bond formations across a wide range of molecular targets.

## Main

Amide bonds are ubiquitous in proteins, natural products, and synthetic materials, and are the most common functional group in medicinal chemistry^1^. However, the synthesis of amides with programmable selectivity remains a central challenge in synthetic chemistry. While conventional couplings provide broad access to amides, they often require protecting groups to restrict reaction to the intended site, adding steps, cost and waste^2^. Such control is especially demanding in protein bioconjugation, where a synthetic molecule must be attached to one amino group among many chemically similar sites while preserving protein function^3^. Enzymes can achieve exceptional selectivity through molecular recognition and can enable shorter, more sustainable synthetic routes^4^. Enzymatic ligation has likewise enabled site-selective protein modification, but adapting these methods to a new target often requires introducing a recognition sequence into the protein or engineering the enzyme’s substrate specificity^5,6^. Developing a biocatalyst for an arbitrary amide-forming reaction can therefore require extensive screening to identify a suitable starting enzyme, followed by directed evolution to achieve the desired selectivity.

We reasoned that deep learning based *de novo* design, which has succeeded in generating enzymes for a variety of chemical reactions^7–10^, could address the challenge of designing highly specific amide bond-forming ligases. Fig 1A schematically depicts a generalised amide-forming reaction between an activated ester donor and a nucleophilic amine acceptor. We reasoned that substrate selectivity could be employed from the outset by designing *de novo* enzymes with donor and acceptor binding pockets flanking a central amide-forming catalytic site (Fig. 1B), and that a cysteine-based catalytic triad mechanism could be particularly effective (Fig 1D). In this mechanism, the catalytic cysteine attacks the donor carbonyl to form an acyl-enzyme intermediate (AEI), which is then attacked by the acceptor amine to afford the desired amide bond. We hypothesised that selectivity for a particular amine within a multifunctional acceptor (regioselectivity, Fig 1B left) or for a defined donor or acceptor enantiomer (stereoselectivity; Fig. 1B middle) could be achieved through appropriately shaped and chemically complementary substrate binding pockets. Chemoselectivity poses an additional challenge: in aqueous solution, the intended acceptor amine must compete effectively with water for attack on the AEI (Fig 1A). We therefore set out to establish design principles that accommodate diverse combinations of acyl donors and amine acceptors, selectively modify the intended amine, and favour aminolysis over hydrolysis, all without recourse to laboratory evolution.

**Figure 1.**
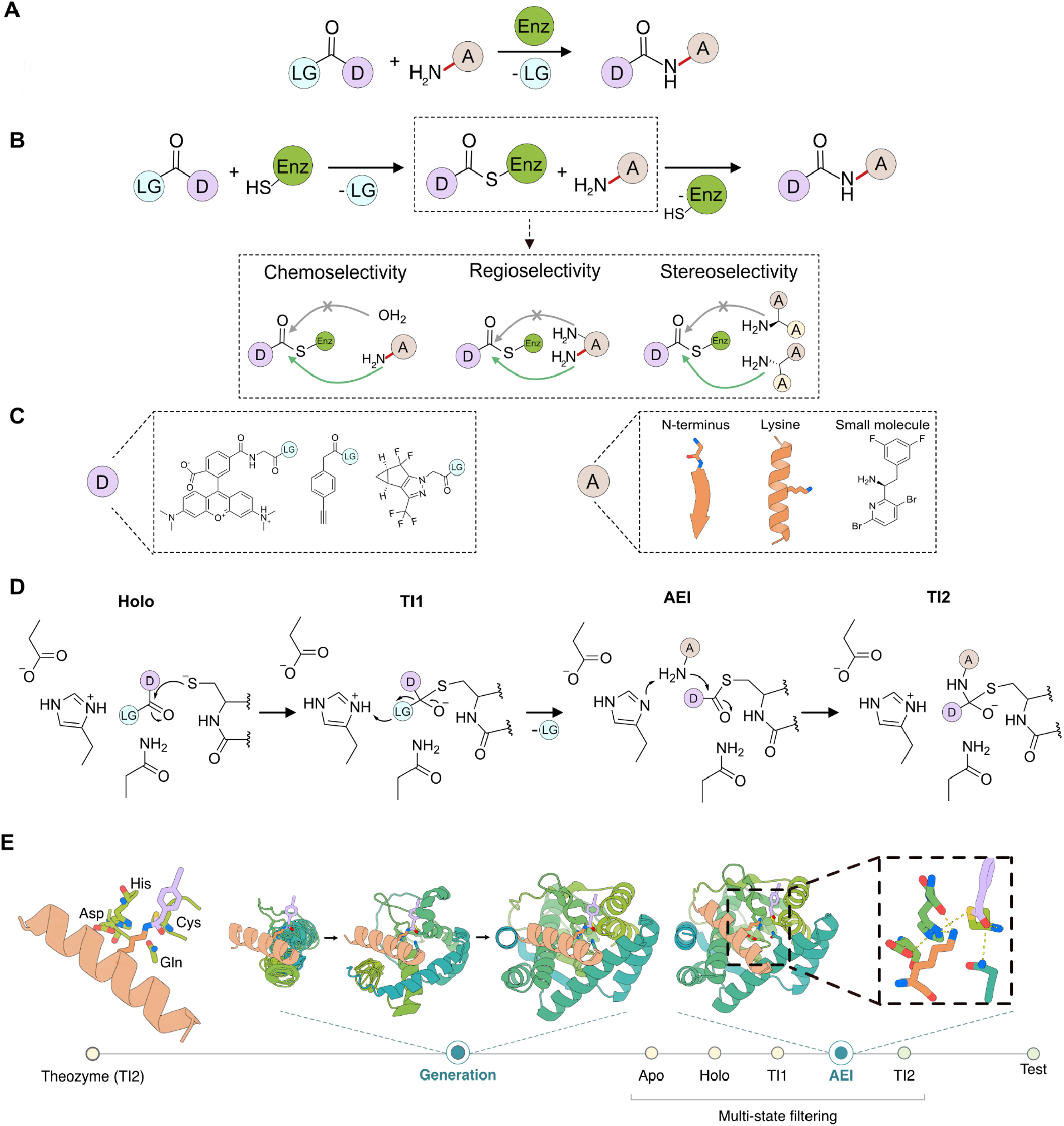
Design strategy for selective amide bond formation. **(A)** Biocatalytic peptide bond formation between an activated ester and an amine. D, donor moiety (purple); A, acceptor (tan); E, enzyme (green); LG, leaving group (cyan). **(B)** Acyl transfer from an activated ester donor to an amine acceptor proceeds through a covalent acyl-enzyme intermediate (AEI). The design objectives include favouring aminolysis of the activated ester over hydrolysis (chemoselectivity), selecting the desired amine acceptor (regioselectivity), and distinguishing enantiomeric acceptors (stereoselectivity) **(C)** The structures at the bottom show the donors and acceptors used for N-terminal peptide acylation, lysine acylation and small-molecule amide formation. **(D)** Proposed catalytic pathway for a Cys–His–Asp active site with a glutamine oxyanion-hole residue, showing the donor-bound state (holo), first tetrahedral intermediate (TI1), AEI and second tetrahedral intermediate (TI2) formed upon amine attack. (**E**) Computational design workflow illustrated for a helical peptide acceptor. Protein scaffolds are generated around a TI2 theozyme and assessed by structure prediction in selected reaction states before experimental testing. The acceptor is orange and the acyl group is purple; dashed lines in the enlarged active-site view indicate modelled hydrogen bonds. Apo refers to the unbound state of the enzyme alone.

### Design strategy for amide-bond formation

For each design case, we began with a cysteine-protease-derived theozyme containing the donor and acceptor in the geometry of the second tetrahedral intermediate (TI2) formed upon amine attack. Cysteine was chosen as the catalytic nucleophile because thioesters are intrinsically more reactive towards amines than the corresponding oxyesters^11^. We explored different catalytic-triad arrangements, as well as backbone and side-chain hydrogen bond donors to stabilise the developing oxyanion. Using RFDiffusion methods^12–14^, we then generated *de novo* protein scaffolds that positioned these idealised active sites and flanking donor and acceptor binding pockets optimally for catalysis. Structure prediction was used to identify designs that maintained the required catalytic and substrate-binding geometries in selected states along the ligation pathway (Fig. 1D). To favour productive aminolysis, we sought to position and orient the acceptor amine for attack on the acyl-enzyme intermediate while limiting water access through active site burial, without compromising substrate binding or product release.

We set out to apply this strategy to first design selective ligases for the acylation of peptide N-terminal α-amines, then the acylation of lysine ε-amines, and finally the stereoselective coupling of two small molecules in a route to the clinically important antiviral agent lenacapavir. Together, these cases test whether *de novo* design can furnish biocatalysts for the synthesis of amides in increasingly demanding chemical transformations.

### *De novo* design of peptide ligases for regioselective peptide modification

To investigate regioselective acylation at an N-terminal α-amine, we selected a target peptide sequence containing an alternating apolar–polar sequence motif (FSLTK), reasoning that its N-terminus could be positioned for aminolysis through strand pairing^10^. The C-terminal lysine provided a competing ε-amine to test regioselectivity. As the acyl donor, we chose an activated ester of TAMRA-conjugated glycine, so that productive peptide-bond formation installs a fluorophore at the peptide N-terminus (Supplementary Fig. 1). The theozyme geometry was based on the active site of tobacco etch virus (TEV) protease (PDB ID: 1lvm)^15^, including the oxyanion loop, in which the backbone amides of the catalytic cysteine and the residue two positions upstream hydrogen bond to the oxyanion in the first and second tetrahedral intermediates (TI1 and TI2).

We used RFDiffusion2 (RFD2)^12^ to generate protein backbones scaffolding this site, and selected designs in which both substrates occupied buried binding pockets (Supplementary Methods). Sequences were designed for the retained backbones with LigandMPNN^16^, and designs were selected whose AlphaFold3 (AF3)^17^ predictions recapitulated the computational design models. Selected designs were then resampled by RFD2 partial diffusion, followed by iterative rounds of ligandMPNN sequence design and AF3 structure prediction to improve agreement with the target catalytic geometry^18^. To favour productive aminolysis, we selected designs that maintained the desired catalytic geometry across the reaction coordinate. Structures were predicted for each state and evaluated against state-specific geometric criteria (Supplementary Methods). We implemented this pipeline using the Chaperone software framework^19^. We selected 51 sequences that passed all six state filters, together with a further 45 sequences that passed only the initial TI2 filter, for experimental characterisation.

The 96 designed sequences, termed N-terminal ligases 1–96 (NL1–NL96) were expressed in *E. coli*, and the FSLTK peptide was prepared by solid-phase synthesis. Ligation product formation and donor hydrolysis were quantified by HPLC (Fig. 2B). Of the 96 designs, ten showed ligase activity and seven showed hydrolase activity. All ten ligase hits and five of the seven hydrolases were obtained from the all-state filtering pipeline, whereas the remaining two hydrolases were selected from the TI2-filtered pool. These results suggest that all-state filtering was crucial for the identification of active ligases. We selected the most active ligase, NL6, for further characterisation (Fig. 2C). To validate NL6’s designed catalytic architecture, we introduced mutations targeting the catalytic Cys–His–Asp triad and oxyanion-stabilizing loop. Mutation of the catalytic cysteine, histidine, and oxyanion-stabilizing loop abolished ligation activity, whereas the aspartate knockout retained residual activity, consistent with a central catalytic role for the Cys–His dyad (Supplementary Fig. 2A).

**Figure 2.**
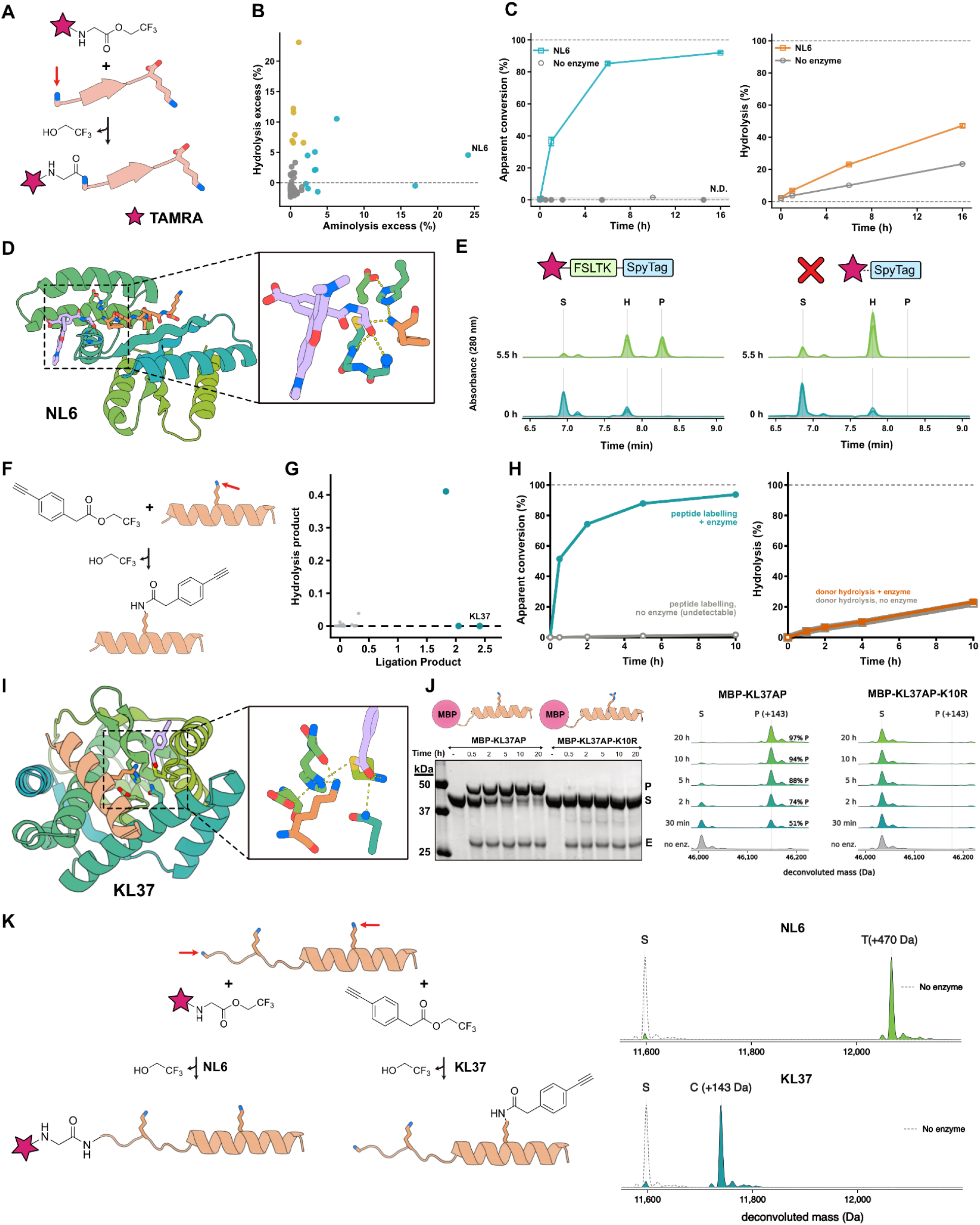
Design of ligases for regioselective peptide acylation. **(A)** N-terminal acylation of FSLTK with the trifluoroethyl ester of TAMRA–glycine. Red arrow, reacting α-amine; the lysine ε-amine is a competing site. **(B)** HPLC screen for N-terminal acylation vs. donor hydrolysis at 27 h; 5 µM design, 100 µM FSLTK, 200 µM TAMRA donor. Points show percentage of ligated product (TAMRA-Gly-FSLTK) and hydrolysed donor (TAMRA-Gly-OH) from 555 nm peak areas, background-subtracted against mean value of the no-enzyme controls (Dashed line) (n = 2). Yellow, 10% most active hydrolases; teal, 10% most active ligases. Data represent single measurements (n = 1). **(C)** Time courses for NL6: apparent FSLTK conversion (teal, 205 nm; left) and donor hydrolysis (orange, 555 nm; right); 10 µM NL6, 50 µM peptide, 100 µM donor. Grey points and line, no-enzyme control; shaded region, range of undetectable conversion; unfilled points, control non-quantifiable conversion. Data represent mean ± SD of three independent reactions (n = 3), with error bars indicating SD. **(D)** AF3 model of NL6 with acyl donor (purple) and peptide acceptor (orange); inset, active site. **(E)** Labelling experiments with SpyTag (blue) versus N-terminally extended FSLTK-Spy (green). Product at 0 and 5.5 h by HPLC (205 and 555 nm); 10 µM NL6, 50 µM peptide, 100 µM donor. **(F)** Acylation of a target lysine ε-amine (red arrow) in a helical peptide with an alkyne-bearing activated ester. **(G)** Initial screen for lysine ligation; KL30, KL33 and KL37 are highlighted in teal. Product formation and donor hydrolysis were measured by mass spectrometry and HPLC, respectively, with both readouts normalized to enzyme concentration. Reaction conditions: 800 µM donor, 70% (v/v) crude-purified MBP–peptide and ligase preparations, 100 mM HEPES pH 8, 50 mM NaCl and 10% DMSO, 20 h at room temperature. **(H)** Time courses of acceptor labelling and donor hydrolysis ± KL37; 4 µM KL37, 16 µM acceptor, 800 µM donor. Data represent single measurements (n = 1) **(I)** AF3 model of the KL37 AEI bound to its helical acceptor; colours as in D. **(J)** Target-lysine dependence of KL37 acylation of an MBP–peptide fusion: SDS–PAGE after PEG48–azide conjugation (E: KL37; S: MBP-AP; P: acylated MBP-AP) (left) and deconvoluted mass spectra (right) for wild type and K10R; 4 µM KL37, 16 µM acceptor, 800 µM donor. **(K)** Acylation of a shared acceptor bearing both recognition elements, in separate reactions with NL6 and TAMRA-donor (+470 Da, top) or KL37 and alkyne donor; 4 µM enzyme, 16 µM acceptor, 500 µM donor. Gel time points in hours; −, no enzyme; S, substrate; P, product (+143 Da); MBP, maltose-binding protein.

To quantify NL6-catalysed substrate-to-product conversion, we incubated 10 µM NL6 with 100 µM TAMRA-Gly-TFE and 50 µM FSLTK in 100 mM HEPES, 50 mM NaCl, and 5% DMSO at pH 7.4, collected time points over 16 h, and measured ligation product formation and donor hydrolysis by HPLC. NL6 converted 90% of the FSLTK substrate to the labelled TAMRA-GSFLTK product within 16 h, corresponding to 4.5 turnovers per enzyme. Over the same period, NL6-mediated hydrolysis accounted for an additional 20% of the donor relative to the no-enzyme control (Fig. 2C). We next confirmed that an extended version of FSLTK fused to SpyTag (FSLTK-SpyTag) was also accepted as a substrate by NL6 (Fig. 2E). To investigate whether NL6 preferentially acylates the N-terminal α-amine over competing lysine ε-amines, we then compared its activity towards FSLTK-SpyTag and SpyTag alone. Removing FSLTK eliminated the intended N-terminal recognition sequence while retaining an alternative N-terminal α-amine and two lysine ε-amines in SpyTag as potential nucleophiles. No ligation product was detected with SpyTag alone, consistent with selective recognition of the FSLTK N-terminus by NL6 (Fig. 2E).

To identify structural features responsible for ligase activity, we retrospectively compared the AF3-predicted catalytic geometries of active and inactive designs from the initial screen. Active ligases and hydrolases showed a tendency to position the acceptor α-amine closer to the catalytic histidine and acyl carbonyl carbon than the inactive designs (Supplementary Fig. 3A). The distributions of these distances were substantially narrower for active ligases than for hydrolases, suggesting that precise positioning of the peptide acceptor was crucial for successful ligase design (Supplementary Fig. 3B).

We next targeted lysine ε-amines, whose higher p*K*_a_ leaves a smaller unprotonated fraction near neutral pH than that of N-terminal α-amines, making aminolysis more challenging. We chose an alkyne-bearing activated ester as the acyl donor to install a handle for subsequent payload conjugation (Fig. 2F, Supplementary Fig. 4) and an α-helical peptide as the acceptor to test substrate recognition beyond strand pairing. The enzyme and acceptor sequences were co-designed to optimise their binding interface and to position the lysine ε-amine for attack on the acyl-enzyme intermediate^9,10^. For this campaign, we constructed the theozyme using the geometry of Ulp1 (PDB ID: 1euv)^20^ to explore alternative cysteine-based catalytic architectures. Here, a glutamine side chain stabilises the oxyanion in addition to the backbone amide nitrogen of the catalytic cysteine. We hypothesised that a backbone carbonyl adjacent to the catalytic histidine could complement the catalytic base by hydrogen bonding to the neutral lysine ε-amine and orienting it for nucleophilic attack, and hence included the histidine backbone in the theozyme. We adapted the N-terminal ligase design pipeline for this motif, replacing residue-based RFD2^12^ with the all atom diffusion model RFdiffusion3 (RFD3)^13^ because of its improved motif-scaffolding performance. Guided by the association between precise amine placement and ligation activity observed in the N-terminal ligase campaign, we applied stringent filters for accurate acceptor-amine placement. This selection yielded 79 designs derived from 23 original backbone scaffolds (see supplementary methods for design pipeline details).

We expressed the enzymes and their cognate peptides separately in *E. coli*, with each peptide fused to the C terminus of maltose-binding protein (MBP). After purification, we screened the enzyme–peptide pairs by mass spectrometry and identified three designs with strong ligation activity: KL30 (lysine ligase 30), KL33 and KL37 (Figure 2G). KL30 also hydrolyzed the donor, as detected by HPLC, but no design showed strong hydrolysis alone. We selected KL37, one of the top hits, for further characterization. KL37 installed the alkyne-bearing acyl group on the MBP–peptide substrate, as detected by mass spectrometry and by a gel shift on SDS–PAGE after conjugation to PEG48–azide (Fig. 2H). With 4 µM KL37, 16 µM MBP–peptide and 800 µM donor, nearly 90% of the acceptor was labelled within 5 h, corresponding to ~3.6 turnovers per enzyme. Over the same period, donor hydrolysis measured by HPLC was indistinguishable from the background rate. To determine whether KL37 modifies the intended lysine rather than one of the other amines in the MBP–peptide, we replaced the target lysine with arginine. No product was detected over 20 h under otherwise identical conditions, despite the 36 remaining lysines and the free N terminus, establishing site selectivity within the MBP–peptide. Individual alanine substitutions of the designed cysteine or aspartate each nearly abolished activity, substitution of histidine substantially also reduced product formation substantially, while the glutamine mutant retained some residual ligase activity (Supplementary Fig. 5). These results support the importance of the designed catalytic motif for function.

Both the N-terminus- and lysine-directed ligases selectively acylated their intended amines. We reasoned that this control over the attachment site should extend to a peptide containing both recognition elements, enabling site selective conjugation to a lysine or N-terminus in the same target. We therefore constructed a new substrate in which the C-terminus of FSLTK was fused to the alpha-helical peptide, followed in turn by the B1 domain from protein G. We then assayed NL6 and KL37 with their cognate donors and fused substrate in separate reactions. Both reactions achieved near-quantitative conversion within 12 h using 4 µM enzyme, 16 µM acceptor, and 500 µM donor, as determined by LC-MS (Fig. 2K). The N-terminal ligase modified only the α-amine, and the lysine ligase only the intended ε-amine; the two designed enzymes selectively address distinct amino groups on the same peptide substrate.

### Structural characterisation and substrate-dependent active-site organisation

To investigate how catalytic geometry depends on reaction state, we compared AF3 predictions of apo NL6 and the NL6 acyl-enzyme intermediate in the presence and absence of the peptide acceptor (Fig. 3A). The apo predictions exhibited substantial variability in oxyanion-loop conformation. In a subset of predictions, part of the oxyanion loop adopted a helical conformation, extending the helix containing the catalytic cysteine and disrupting the geometry required for oxyanion stabilization, thereby compromising the ability of the oxyanion-binding backbone amides to form hydrogen bonds with the oxyanion (Fig. 3A, left). In acceptor-free AEI predictions, the oxyanion loop adopted a more consistent, non-helical conformation, but the acyl group occupied either the designed binding pocket or the peptide-binding site (Fig. 3A, center). With the acceptor present, the acyl group was more consistently positioned in its intended pocket and the peptide adopted the designed conformation (Fig. 3A, right). The median distances between the acyl carbonyl oxygen and the oxyanion-hole backbone amides decreased from 3.8 to 3.0 Å for OXH_1_ and from 5.4 to 3.8 Å for OXH_2_, with substantially narrower distributions across predictions (Fig. 3B). We therefore hypothesised that acceptor binding aligns the acyl group with the oxyanion hole for attack by the N-terminal α-amine. To assess whether NL6 can in fact adopt this local architecture, we determined the peptide-free crystal structure of the inactive C49A mutant at 2.18 Å resolution (Fig. 3C). The crystal structure closely recapitulated the designed architecture, both globally (Cα RMSD of 0.73 Å) and within the catalytic motif comprising the catalytic triad and oxyanion-binding loop (all atom RMSD 0.55 Å following global Cα alignment). A sulfate ion occupied the designed oxyanion-binding pocket, forming hydrogen bonds with both oxyanion-binding backbone amides and a nearby lysine not originally implicated in substrate recognition. By mimicking substrate binding, the sulfate ion appears to stabilise the oxyanion loop and catalytic cysteine helix in their designed conformations, demonstrating that the active site can adopt a geometry compatible with stabilisation of the tetrahedral intermediate.

**Figure 3.**
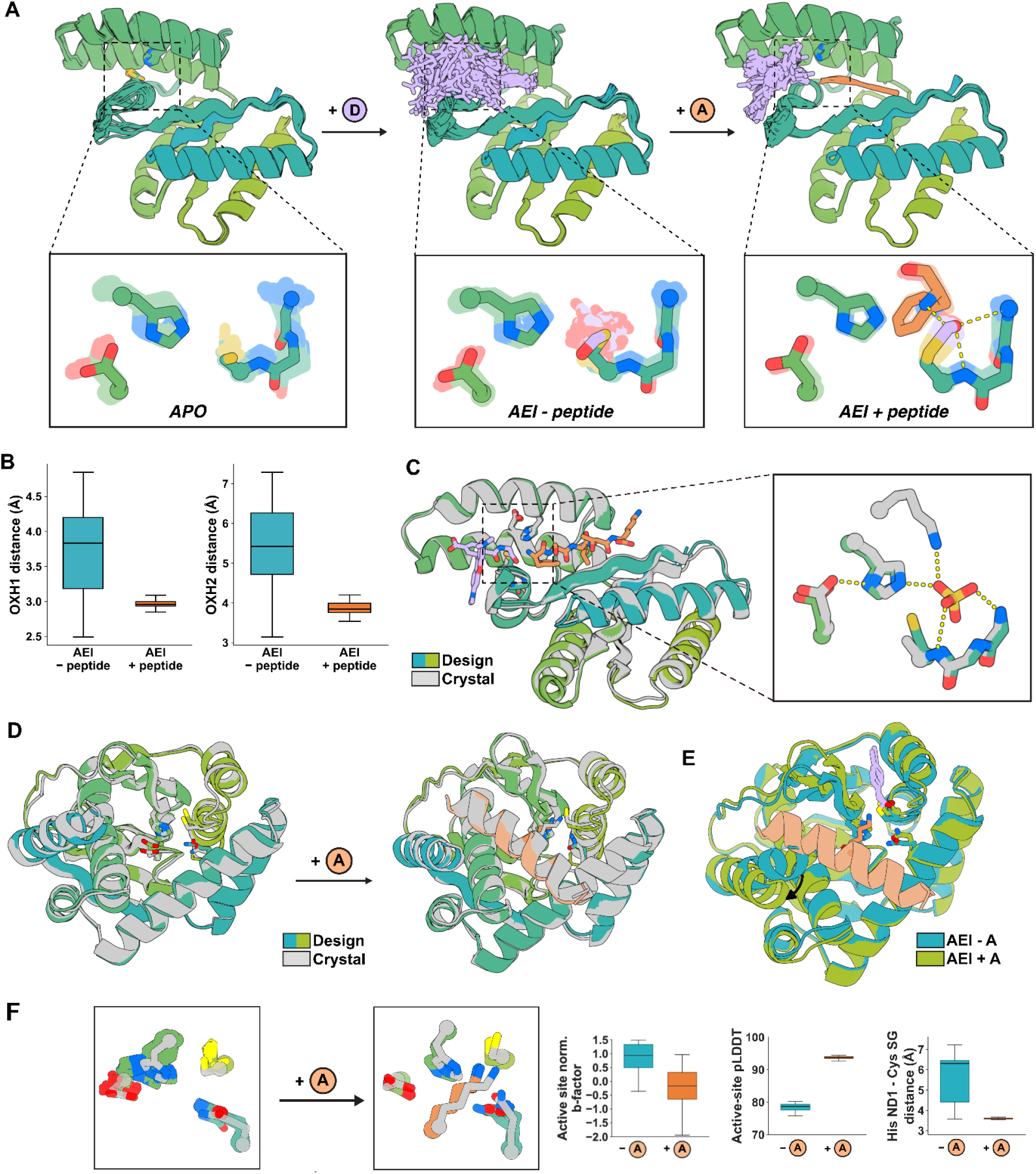
Structural characterisation and substrate-dependent active-site organisation. **(A)** Superposed AF3 predictions of apo NL6 (left), the AEI without acceptor (center), and the AEI with acceptor (right). Acyl group, purple; peptide acceptor, orange. Closeups show top predictions in the ensembles. **(B)** Distributions of distances between the acyl carbonyl oxygen and the two oxyanion-hole backbone amides, OXH1 and OXH2, in NL6 AEI predictions without (cyan) and with (orange) acceptor. **(C)** Superposition of the NL6 AF3 model (coloured) and the peptide-free crystal structure of the inactive C49A mutant (grey). The acyl group and peptide are present only in the model. The inset shows the crystallographic sulfate ion and its hydrogen bonds to the two oxyanion-hole backbone amides, the catalytic histidine and a nearby lysine. **(D)** Superpositions of KL37 AEI predictions (coloured) and crystal structures (grey) of apo KL37 (left) and KL37 with its cognate acceptor fused to the C terminus (right), showing the corresponding backbone arrangements. Neither crystal structure contains a modelled acyl donor. **(E)** Overlay of AF3 AEI models of KL37 with (green) and without (teal) acceptor. Helix movement is indicated with an arrow. **(F)** KL37 active-site overlays without and with acceptor (top), and distributions of normalised crystallographic B factors, AF3 active-site pLDDT and the His119 Nδ1–Cys176 Sγ distance (bottom, left to right). B factors are shown for the 17 side-chain heavy atoms of Gln49, His119, Asp136 and Cys176, standardized separately for each structure using the mean and population standard deviation of B factors for 20 matched enzyme backbone atoms in the local neighbourhood (within 6 Å of any His119 side-chain heavy atom in both structures, excluding all four active-site residues). The pLDDT and distance distributions contain 25 AF3 models per state; pLDDT is averaged over the same 17 active-site atoms within each model. Cyan denotes the absence of the acceptor and orange denotes its presence. For (E), boxes span the interquartile range, centre lines are medians and whiskers extend to the most extreme points within 1.5x the interquartile range; outliers are omitted.

We next asked whether binding of the acceptor peptide similarly organises the KL37 active site, comparing AF3 predictions of the AEI with and without peptide (25 models per state). Without the helical acceptor, the catalytic histidine varied in position and active-site prediction confidence was lower (79 median pLDDT versus 94 with acceptor; Fig. 3E). The His119 Nδ1–Cys176 Sγ distance ranged from 3.6 to 7.2 Å without peptide but from 3.5 to 3.7 Å with peptide, indicating more reproducible recovery of the designed catalytic arrangement. To investigate the predicted rearrangement experimentally, we determined crystal structures of apo KL37 and a construct with the cognate acceptor fused to its C terminus, 3.8 and 3.1 Å resolution. Both the apo and acceptor-bound structures recapitulated the design globally (Cα RMSD of 0.82 Å for apo and 0.60 Å for acceptor-bound) and within the catalytic tetrad (all-atom RMSD after global Cα superposition of 0.86 Å for apo and 0.69 Å for acceptor-bound). In the bound structure, the acceptor peptide closely matched its designed pose in the binding pocket (Cα RMSD 0.526 Å; acceptor lysine heavy-atom RMSD 0.763 Å), demonstrating that the design procedure accurately positioned both the peptide backbone and the reacting lysine. However, the two structures differ in the placement of the N-terminal helix (Fig. 3E) in a manner that’s consistent with the predictions. In the apo structure, this helix adopts an orientation that partially occludes the acceptor site; in the acceptor-bound structure it is displaced by 6.8 Å (maximum Cα displacement after global Cα superposition), opening space for the peptide in its designed conformation. (Fig. 3E). Active-site B factors were lower in the peptide-bound structure, consistent with greater local order in that crystal form (Fig. 3F).

Together, these observations support a model in which the amine acceptor both supplies the nucleophile and organises the catalytic geometry for aminolysis. Such coupling offers an explanation for the ability of NL6 and KL37 to promote amine acylation preferentially over competing donor hydrolysis. In the acceptor-free state, misalignment of the acyl group and oxyanion hole could disfavour hydrolysis in NL6; in KL37, incomplete positioning of the catalytic histidine could limit water activation, and the hydrophobic active-site environment may further disfavour productive water attack. A related geometric explanation was proposed for earlier serine-hydrolase designs that formed covalent acyl-enzyme intermediates but did not hydrolyse them efficiently, where modelling suggested that acyl-group reorientation disrupted productive water attack^7^. Our design procedure prioritised the position and orientation of the acceptor amine for attack on the acyl carbonyl. Optimising this geometry with the peptide present could favour active sites whose productive organisation depends on peptide contacts. In AF3 predictions of the 79 KL designs, peptide inclusion shortened Cys–His in most designs and increased active-site prediction confidence in all (Supplementary Fig. 6). Curiously, the ligase KL30 that also hydrolysed the donor, showed much smaller changes in this distance than KL37 (Supplementary Fig. 6). Consistent with this model, retrospective analysis of the 96 NLs showed substantially reduced variability in predicted oxyanion-loop geometry upon acceptor inclusion for active ligases, with a less pronounced effect on the hydrolases (Supplementary Fig. 3C). Future campaigns could exploit this dependence through explicit negative design, selecting against preorganisation for water attack in the acceptor-free AEI while retaining donor acylation and productive amine-attack geometry with the acceptor.

### *De novo* design of ligases for stereoselective amide bond formation in lenacapavir

We next sought to extend control over amine reactivity from site selection in peptides to enantiomer selection in small-molecule synthesis. Compared with peptidic substrates, small-molecule substrates pose a greater design challenge as they occupy a far more diverse chemical and conformational space. Moreover, protein–small-molecule complexes are sparsely represented in the PDB relative to protein–protein and protein–peptide interfaces. As a stringent test for our design pipeline, we chose an amide-forming step in the synthesis of lenacapavir, a long-acting antiretroviral agent whose manufacturing cost limits global access^21–23^. The target transformation couples an activated trifluoroethyl ester to the (*S*)-enantiomer of a sterically demanding chiral amine (Fig. 4A, Supplementary Fig. 7), requiring an active site that accommodates the correct amine enantiomer while favouring aminolysis over competing hydrolysis of the activated ester.

**Figure 4.**
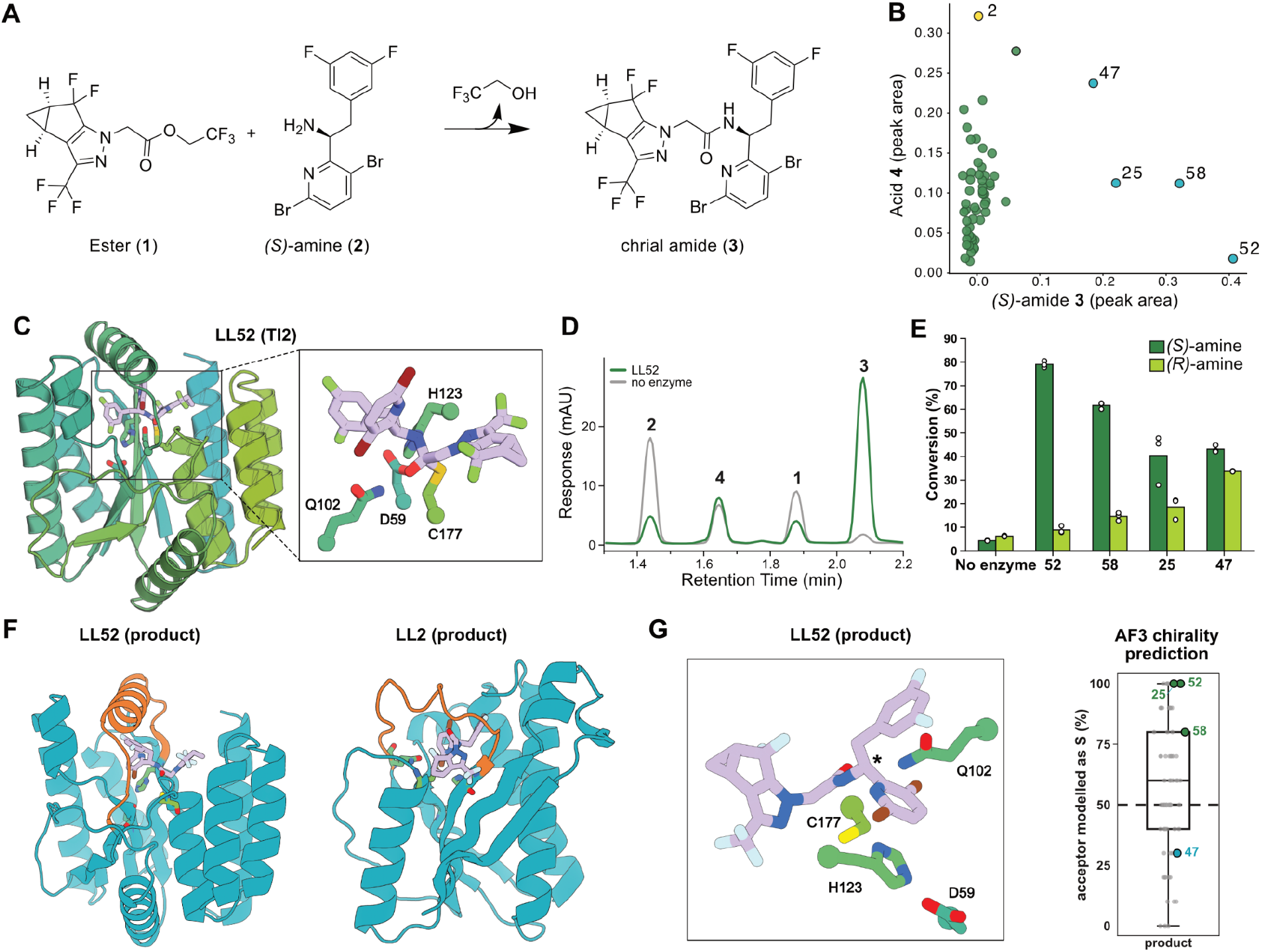
Stereoselective assembly of a lenacapavir precursor by designed ligases. **(A)** Target reaction coupling activated ester **1** with (*S*)-amine **2** to form the chiral amide **3. (B)**, Background-normalised HPLC peak areas of amide **3** and the donor hydrolysis product, acid **4**, across the 55 characterised designs. Designs LL25, LL47, LL52 and LL58 are highlighted in cyan. **(C)** AF3 model of LL52 in the TI2 state. **(D)** HPLC traces for reactions with LL52 (green), no enzyme (grey). Reactions contained 0.5 mM acceptor, 1 mM donor and, where indicated, 50 µM enzyme, and were incubated for 16 h at 30 °C in 0.1 M potassium phosphate, pH 6, with 20% DMSO. **(E)** Amide formation in separate reactions with the (*S*)- and (*R*)-amines for LL52 and LL58. Reactions contained 0.5 mM acceptor, 1 mM donor and, where indicated, 50 µM enzyme, and were incubated for 16 h at 30 °C in 0.1 M potassium phosphate, pH 6, with 20% DMSO **(F)** AF3 predictions of LL52 and LL2 with product state showing a well-structured helical lid (orange) covering the active site of 52 but not in LL2 **(G)** Active site of product bound state of LL52 with stereocentre next to amine labelled with (*). Predicted chirality of stereocentre next to amine across AF3 predictions with 10 seeds of all designs are shown as box plots. LL52, LL58, LL25 are shown in green and LL47 in teal.

We constructed a Ulp1-derived theozyme incorporating a model of TI2 for this reaction. An initial RFD3 campaign targeting this transformation yielded no active designs prompting us to use RFdiffusion4 (RFD4)-Proteina, a latent diffusion model that jointly designs sequence and structure, to generate proteins scaffolding the theozyme^14^. AF3 predictions were filtered for pLDDT and active-site geometry, and passing designs were modelled in the acyl-enzyme intermediate state to assess retention of the catalytic geometry and chirality (Supplementary Methods). Nineteen designs passed these filters; partial diffusion and LigandMPNN sequence redesign added 30 and 18 variants, respectively, yielding 67 candidates for experimental testing.

55 of 67 designs expressed well in *E. coli*, and four of the 55 designs catalysed formation of the desired amide (Supplementary Fig. 8). LL52 converted 79% of the (*S*)-amine to the target amide using 50 µM enzyme, 0.5 mM acceptor amine and 1 mM acyl donor after 16 h at 30 °C, without increasing donor hydrolysis above background levels (0.29 versus 0.26 mM acid in the no-enzyme control; Fig. 4B-E). Mutation of the designed cysteine decreased conversion 12-fold in LL52 and resulted in no measurable activity over background for LL58, supporting its contribution to catalysis (Supplementary Fig. 9). Comparing separate reactions with the (*S*)- and (*R*)-amines, LL52, LL58 and LL47 preferentially coupled the desired (*S*) enantiomer, LL52 with a 28-fold preference, LL58 with a 7-fold preference and LL47 with a 3-fold preference over the (*R*) enantiomer after background correction (Fig. 4E). This shows that our amide ligase design platform can deliver stereoselective ligases for a key step in lenacapavir synthesis.

We next asked what distinguishes the amide-forming designs from those that only hydrolyse the donor. Of the 19 original RFD4-Proteina designed backbones, 9 have a well-structured lid covering the active site, and the four amide-forming ligases (LL25, LL47, LL52 and LL58) are in this subset (Supplementary Fig. 10). By contrast, LL2, which catalysed only donor hydrolysis with no detectable amide formation, has a disordered loop rather than a lid over the pocket (Fig. 4F). A structured lid could shield the acyl-enzyme intermediate from solvent, whereas a disordered loop would leave it accessible and permit hydrolysis. We did not filter on the enzyme–product state in our design campaign, but found in retrospect that when this state was modelled with AF3, the amide-forming designs (LL25, LL47, LL52 and LL58) consistently recovered the target product-state geometry while the hydrolysing design (LL2) did not (Supplementary Fig. 11). We next analysed the potential of the product state geometry to serve as a stereoselectivity filter by predicting across multiple seeds whether AF3 recapitulates the desired product enantiomer. Across 10 AF3 seeds per design, LL52, LL25 and LL58 ranked among the top designs, and LL52, the most stereoselective of the three, was predicted to contain the (*S*)-configured amine centre in all 10 models. In contrast, LL47 that showed nearly no stereoselectivity in experimental characterisation ranked at the lower end for this metric. These results suggest that such product state analyses could serve as useful filters in future ligase design campaigns.

## Discussion

Here we present a computational platform for the *de novo* design of ligases for selective amide bond formation. We scaffolded cysteine-protease-derived theozymes containing a cysteine–histidine–aspartate triad, selected backbones for substrate burial, and filtered designs for catalytic geometry across key reaction intermediates. Screening 96 or fewer designed sequences per campaign yielded ligases that acylate peptide N-termini or lysine side chains with programmed regioselectivity, or catalyse a stereoselective amide-forming step in lenacapavir synthesis. In all three cases, the ligases showed strong chemoselectivity for amide formation over hydrolysis. This highlights the utility of our platform to deliver bespoke ligases with modest experimental effort.

Progressing from RFD2 to RFD3, and RFD4-Proteina, we were able to meet increasingly demanding substrate recognition and selectivity requirements. While RFD2 and RFD3 enabled the design of ligases for site-selectively conjugating small molecules to peptides, an initial campaign targeting the lenacapavir transformation using RFD3 yielded no active designs. However, using RFD4-Proteina produced several active ligases, likely due to its improved representation of ligand functional groups, greater adherence to specified stereochemistry, and more effective scaffolding of ligand-containing catalytic motifs^14^. These results illustrate how advances in generative modelling can broaden the application of our strategy to complex pharmaceutical substrates.

We anticipated that competing hydrolysis of the acyl-enzyme intermediate would be a major obstacle to designing chemoselective ligases. Yet we obtained several ligases that catalysed amide formation without detectably accelerating donor hydrolysis. Their characterisation suggests two complementary strategies for favouring aminolysis over hydrolysis. In the peptide ligases, structural predictions and crystal structures support a model in which acceptor binding promotes productive active-site organisation; in the lenacapavir ligase, a hydrophobic helix is predicted to cover the active site and may thereby exclude water. Both strategies have counterparts in natural enzymes that must commit a reactive acyl or amide intermediate to an intended nucleophile rather than to water. Sortase A, a cysteine transpeptidase that ligates a sorting signal to an incoming amine in competition with hydrolysis of the same thioacyl intermediate, holds its catalytic cysteine and histidine too far apart in the resting state to interact, and substrate binding closes an active-site loop and rotates the two residues into catalytically competent positions^24,25^. Nonribosomal peptide synthetase thioesterase domains face the same partitioning problem, releasing their product by macrocyclisation in competition with hydrolysis of a common acyl-enzyme intermediate, and carry a lipase-like lid of three α-helices that interconverts between open and closed states over the active site^26^.

These results broaden the scope of de novo enzyme design beyond hydrolytic chemistry and model reactions to synthetically useful bond-forming transformations. In all three design cases, we achieved near-quantitative conversion of the amine to the desired product. Notably, the N-terminal and lenacapavir ligases required only a twofold excess of acyl donor, highlighting the potential for substrate-efficient biocatalytic processes relevant to pharmaceutical synthesis. The stereoselectivity of the lenacapavir ligase opens the door to (dynamic) kinetic resolution of racemic substrates, offering alternative routes to a key precursor that could further reduce manufacturing costs. In an accompanying manuscript, we show that the same design approach yields ligases that ubiquitylate Proliferating Cell Nuclear Antigen (PCNA) at a single site among 45 competing lysines^27^, demonstrating how the platform generalises to the selective modification of large globular proteins. Such site-specific attachment of functional molecules could support the generation of defined protein conjugates, ranging from probes and imaging agents to antibody–drug conjugates. By coupling designed substrate recognition to amide bond formation, our approach provides a programmable route to selective small-molecule synthesis and protein conjugation.

## Acknowledgments

This work was funded by the Gates Foundation (INV-092396), an EPSRC Centre to Centre Partnership between MIB and the IPD (EP/Z531157/1), Gates INV-092950 (B.S., A.K., A.K.B., X.L., D.E., R.I.B. and M.W.K), the Grantham foundation (S.S., N.C. and J.B.), DARPA D26AP50001 (T.F., S.S.M and A.S.) an EMBO postdoctoral fellowship to M.G (ALTF 226-2026), a Schmidt fellowship to A.C.H., a scholarship from the Tianjin Institute of Industrial Biotechnology, Chinese Academy of Sciences (TIB-CAS, Z.X.), Schmidt Sciences (D.B.) and the Howard Hughes Medical Institute (D.B., S.B. and L.D.L.). This research used resources ID-17-1(AMX) of the National Synchrotron Light Source II, a U.S. Department of Energy (DOE) Office of Science User Facility operated for the DOE Office of Science by Brookhaven National Laboratory under Contract No. DE-SC0012704. The center for BioMolecular Structure

(CBMS) is primarily supported by the National Institutes of Health, National Institute of General Medical Sciences (NIGMS) through a center Core P30 Grant (P30GM133893), and by the DOE Office of Biological and Environmental Research (KP1607011). We are grateful to Dr. John Walker, Dr. John Dillon, Dr. Trevor Laird and Dr. Mark Krook for helpful discussions on enzymatic synthesis of pharmaceutical intermediates.

