## Supplementary Information for "Programmable amide bond formation using *de novo* ligases"

### Table of contents

#### 1 Supplementary figures

#### 2 Supplementary methods

##### 2.1 Computational design pipeline

2.1.1 Overall design pipeline

2.1.2 N-term ligase design campaign

2.1.3 Lysine ligase design campaign

2.1.4 Lenacapavir ligase design campaign

##### 2.2 Experimental materials and methods

2.2.1 Genetic constructs

2.2.2 Molecular cloning

2.2.3 Protein expression and purification

2.2.4 Hydrolysis and aminolysis activity screen

2.2.4.1 General procedure for N-term ligase screening

2.2.4.2 General procedure for lysine ligase screening

2.2.4.3 General procedure for analytical scale biotransformations for lenacapavir-API amide ligation

2.2.5 LC/MS assays and analysis

2.2.5.1 N-term

2.2.5.2 Lysine ligase

2.2.6 HPLC assays

2.2.7 Gel assays

2.2.8 X-ray crystallography

2.2.9 Analysis of X-ray crystal structure and AF3 predictions

#### 3 Sequences

##### 3.1 General proteins

##### 3.2 N-terminal ligases

##### 3.3 Lysine ligases

##### 3.4 Lenacapavir ligases

### 1 Supplementary figures

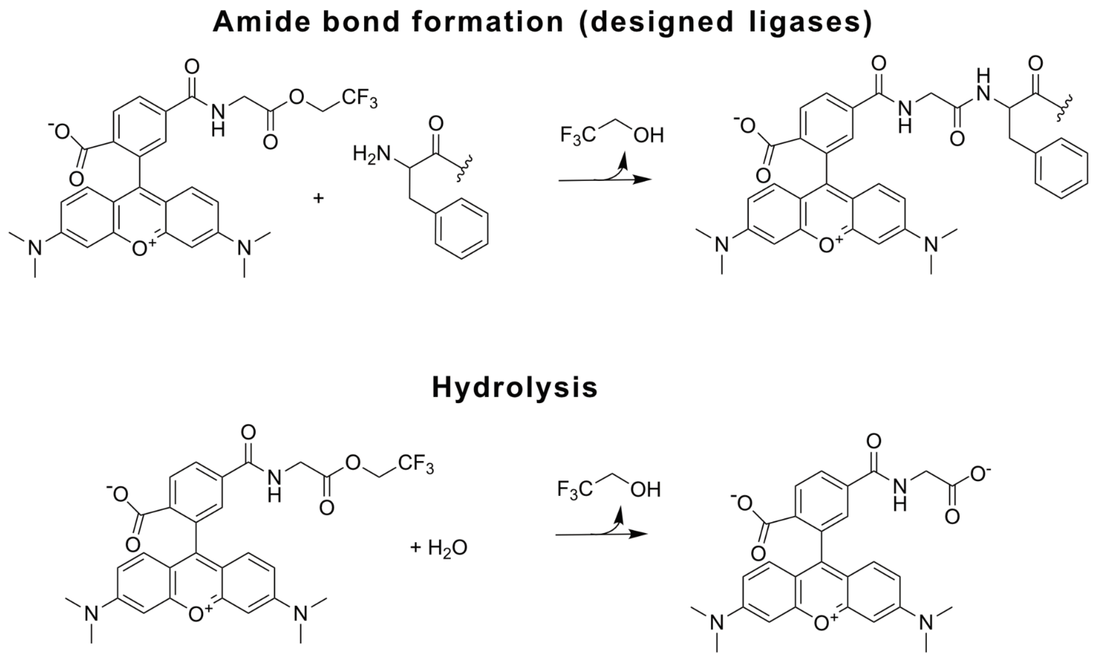

**Supplementary Figure 1 | Reaction scheme for N-terminal labelling reaction.** Both the N-terminal labelling of the FSLTK peptide and the hydrolytic side reaction are shown.

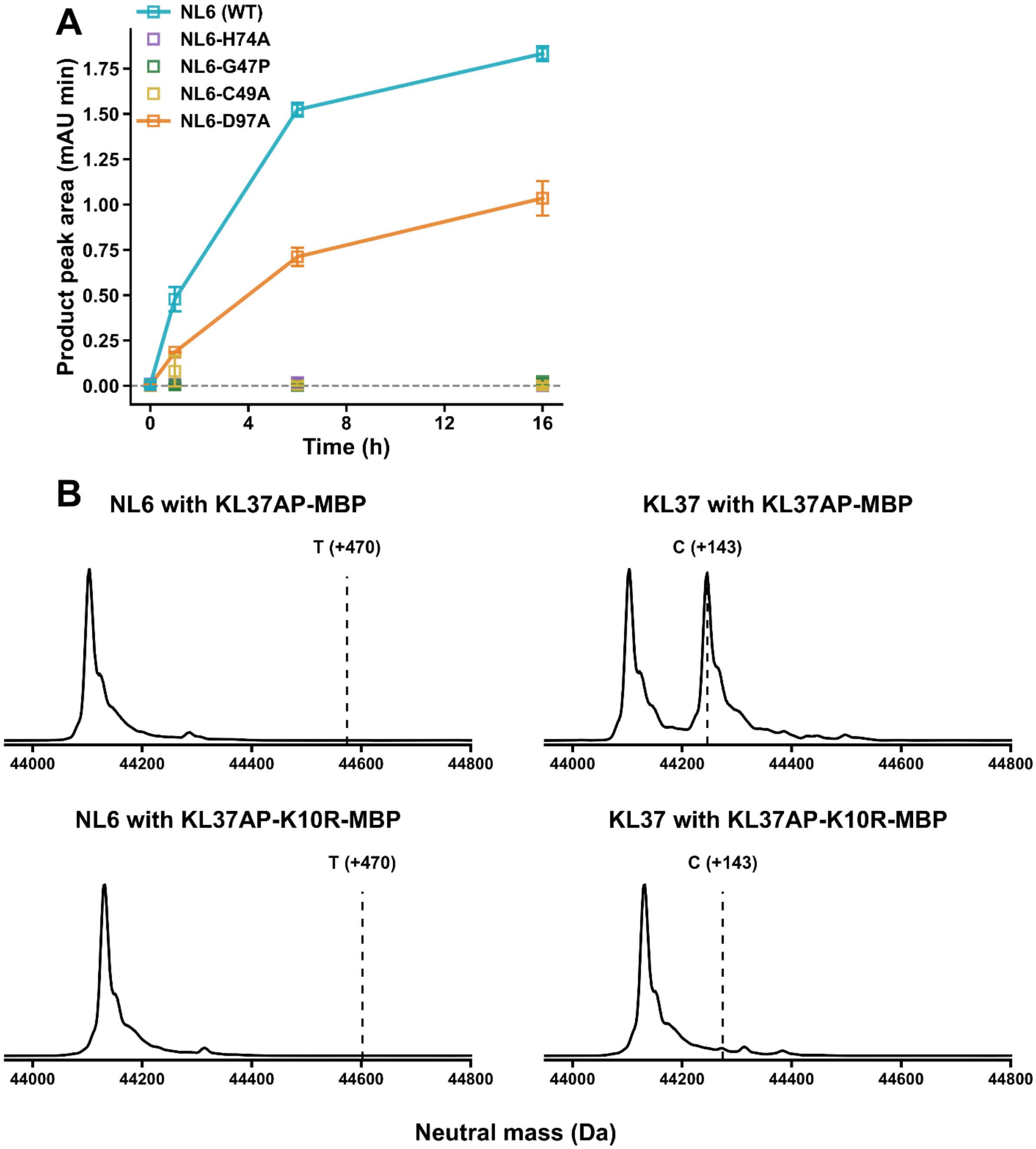

**Supplementary Figure 2 | Catalytic residue dependence and substrate selectivity of NL6. (A)** Time-course analysis of aminolysis activity of N-terminal ligase 6 (NL6) and catalytic knockout variants targeting the catalytic cysteine (NL6-C49A), histidine (NL6-H74A), aspartate (NL6-D97A), and oxyanion-stabilizing loop (NL6-G47P). Reactions contained 10 µM enzyme, 50 µM FSLTK acceptor, and 100 µM TAMRA-Gly-TFE donor and were monitored by HPLC over 16 h. Product formation is reported as the integrated chromatographic peak area (mAU·min). Data represent mean ± SD of three independent reactions (n = 3); Non-detectable peak areas are shown with solid fill with zero area. **(B)** Deconvoluted LC–MS spectra of the lysine ligase KL37 substrate KL37AP-MBP (top) and its K-to-R mutant KL37AP-K10R-MBP (bottom) following incubation with NL6 and TAMRA-Gly-TFE or KL37 and alkyne donor. Dashed lines (left) indicate the expected mass shift (+470 Da) upon TAMRA-Gly acylation. Expected mass shift indicated with dashed line (right) from KL37 acylating cognate substrate is observed with mass shift (+143 Da), but not in target lysine knockout (K10R). No corresponding product peaks were detected for either substrate with NL6 and TAMRA-Gly-TFE, indicating that NL6 does not appreciably acylate these noncognate peptide substrates under the conditions tested. Reactions contained 4 μM enzyme, 16 μM peptide acceptor and 500 μM donor and measured by LC-MS after 6 hours.

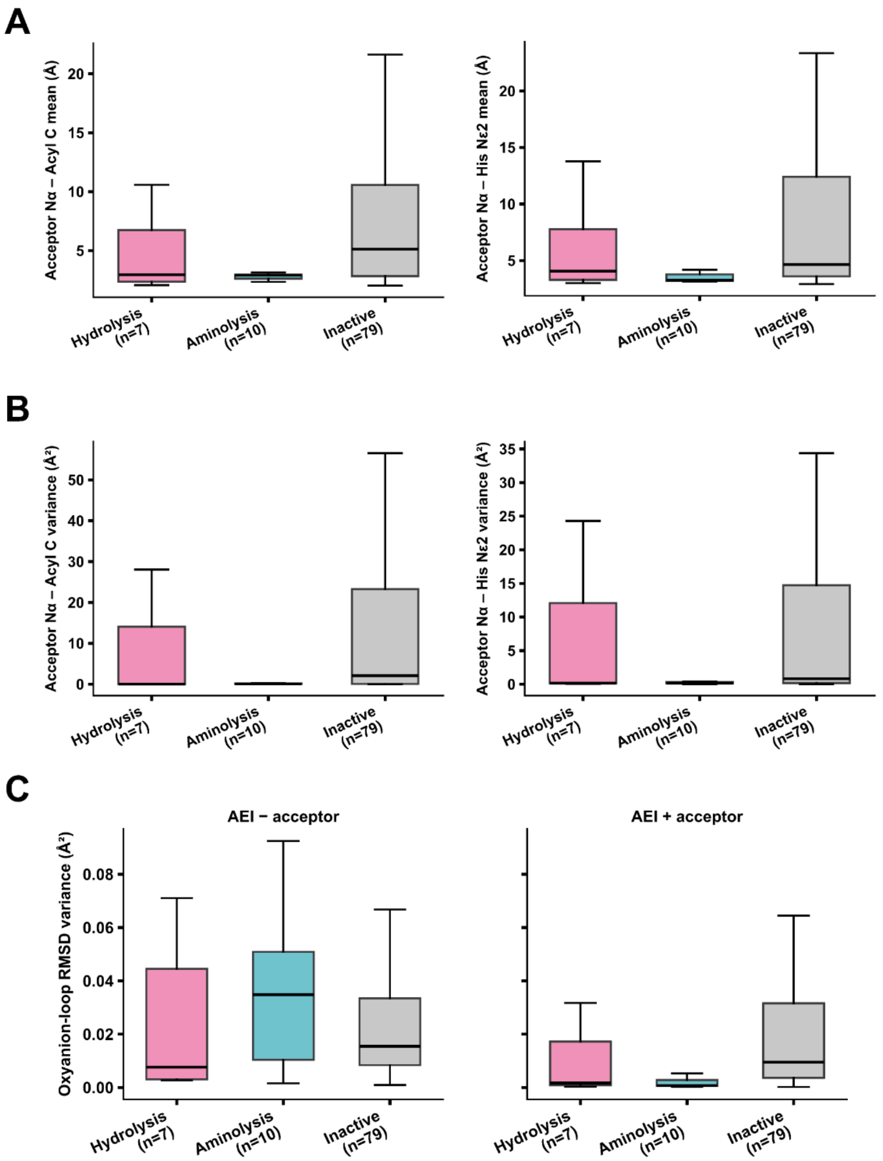

**Supplementary Figure 3 | Retrospective analysis of catalytic geometry and active-site preorganisation in N-terminal ligases.** The 96 designs were classified as active, based on hydrolysis (pink) and/or aminolysis activity (cyan), or inactive (grey). Fifty AlphaFold3 predictions per design and state were analysed. **(A)** Mean distances between the acceptor N-terminal α-amine and the reactive acyl carbonyl carbon or catalytic histidine Nε2 in predicted acyl-enzyme intermediate (AEI) structures in the presence of peptide acceptor. **(B)** Variance of the corresponding distances across predictions for each design, reflecting the consistency of predicted acceptor positioning relative to the catalytic centre. **(C)** Variance in root-mean-square deviation (RMSD) from the original designed catalytic motif, calculated for the full motif, catalytic triad, and oxyanion loop across predictions of the apo state, the AEI without acceptor, and the AEI with acceptor. Aminolysis-active designs exhibit reduced variability in predicted oxyanion-loop geometry upon acceptor inclusion, whereas this reduction is less pronounced for hydrolysis-active designs. Boxplots summarise per-design values within each activity class; centre lines indicate medians, boxes indicate interquartile ranges, and whiskers extend to the most extreme values within 1.5x the interquartile range. RMSDs were calculated relative to the original designed motif using the selected catalytic side-chain and backbone atoms. Variance reflects differences among independently predicted structures rather than experimentally measured conformational dynamics.

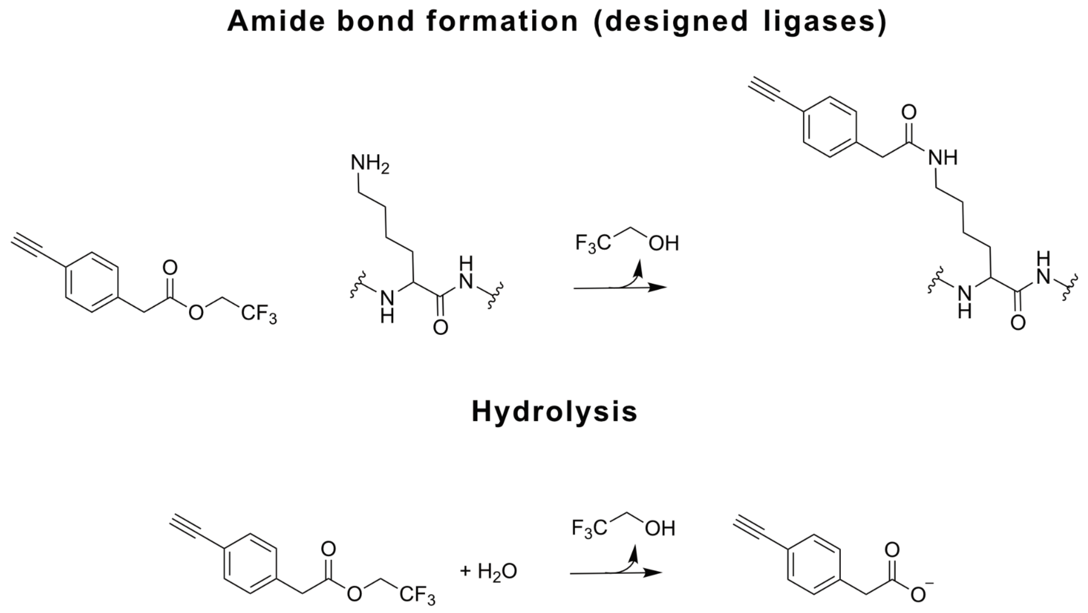

**Supplementary Figure 4 | Reaction scheme for the lysine labelling reaction.** Both the amide bond-forming reaction between the acyl donor and a lysine and the hydrolysis side reaction of the acyl donor are shown.

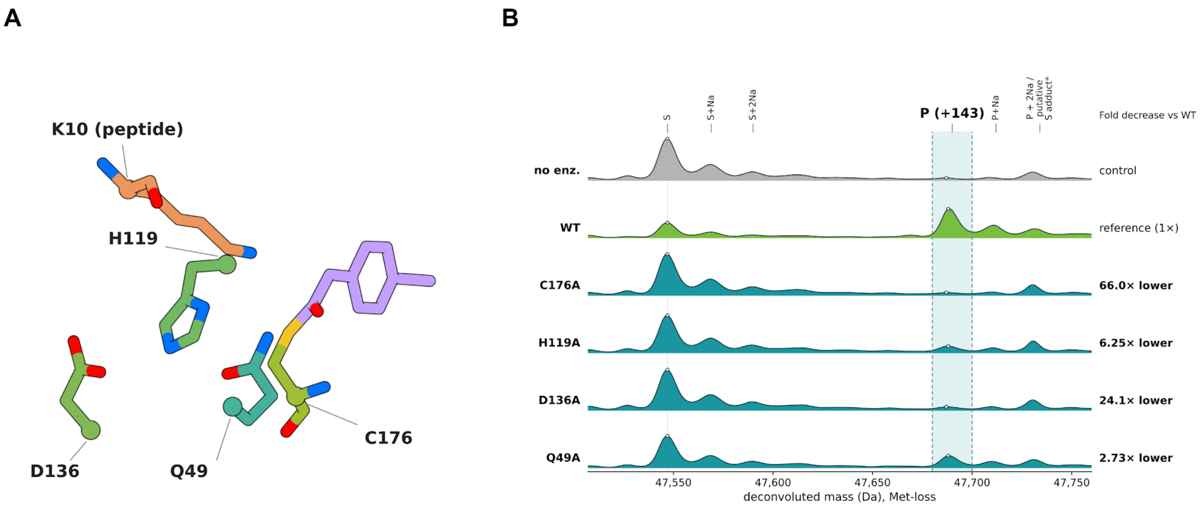

**Supplementary Figure 5 | KL37 active-site model and apparent ligation activity of alanine-substitution variants.** Active-site view of the top AlphaFold 3 prediction of the KL37 acyl-enzyme intermediate bound to its cognate peptide. Gln49, His119, Asp136, Cys176 from the enzyme and Lys10 from the peptide substrate are labelled; the acyl group is modelled covalently attached to Cys176. Enzyme carbons are green/teal, Lys10 orange and the acyl group lavender; nitrogen, oxygen and sulfur are blue, red and yellow. This is a predicted intermediate. **(B)** Deconvoluted intact-protein MS after 5-h reactions at 37 °C in 100 mM HEPES, 50 mM NaCl, pH 8, and 10% DMSO, with 800 µM donor, cognate peptide–MBP acceptor (at 10 µM) and WT or variant enzyme (at 2 µM), alongside a no-enzyme control. S is the substrate (mass calculated - Met); P is the product expected at +143 Da. All spectra show 47,507–47,760 Da; "+Na" denotes sodium-adduct assignments. The shaded region (47,680–47,700 Da) marks the first product peak. Fold decreases at right are $(h_{WT} - h_0)/(h_{variant} - h_0)$, where $h = P/(S + P)$ uses the first substrate and product peak heights and $h_0$ is the no-enzyme value. * P + 2Na overlaps a putative AEBSF–substrate adduct (expected Δm, +183.2 Da; observed Δm, +183 Da, from 47,547 to 47,730 Da in the no-enzyme control). An apparent ligated counterpart is also observed (not shown); the chemical identity remains unconfirmed.

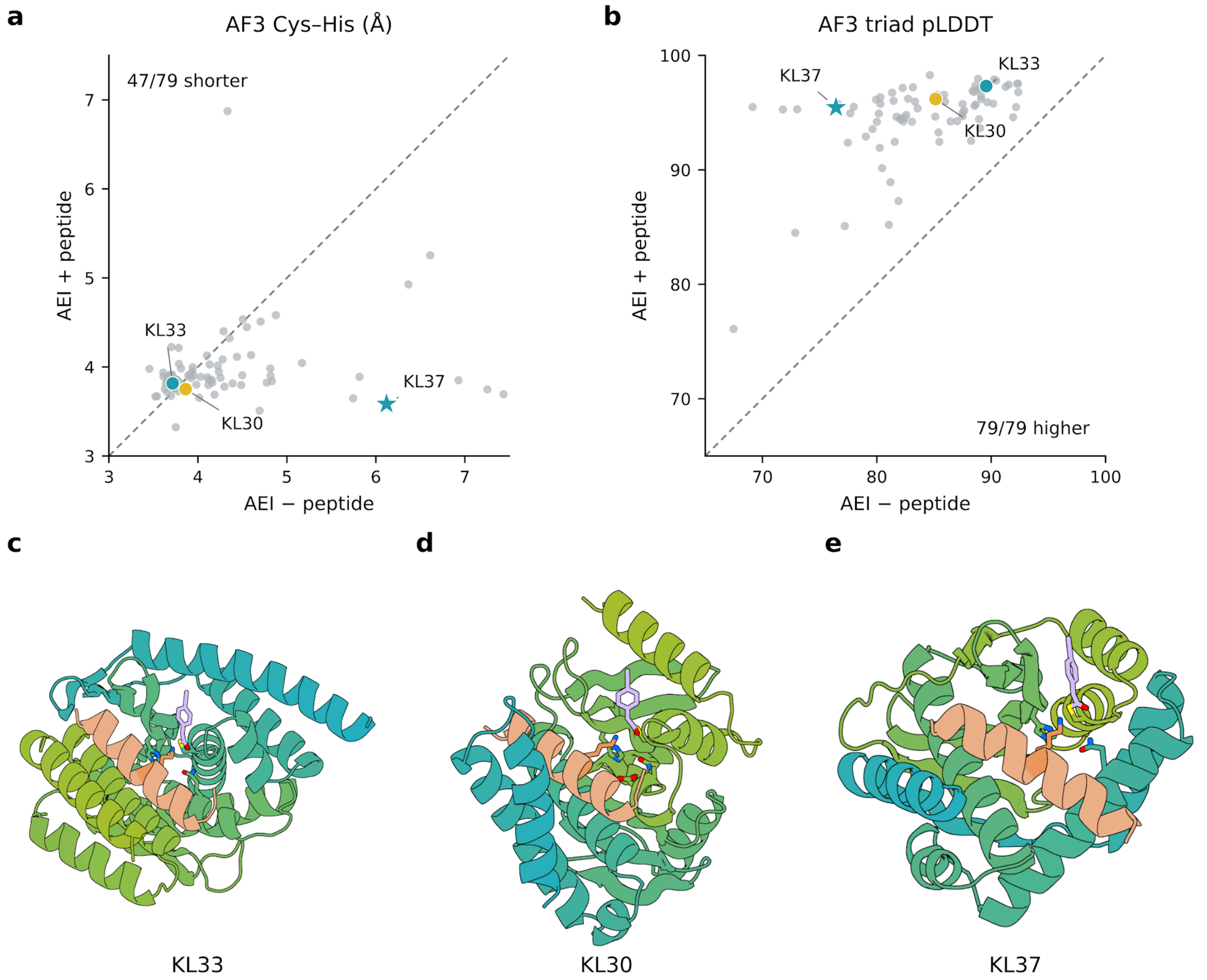

**Supplementary Figure 6 | Peptide-dependent organisation of predicted lysine-ligase active sites. (A and B)** Ensemble-mean Cys SG–His ND1 distance (Å; a) and Cys–His–Asp triad prediction confidence (pLDDT; b) in AlphaFold3 (AF3) predictions of the acyl-enzyme intermediate without (x axis) and with (y axis) acceptor peptide. Each point represents one of 79 designs, with 50 predictions per state (10 seeds × 5 samples). Triad pLDDT was averaged equally over 12 side-chain heavy atoms (Cys CB/SG; His CB/CG/ND1/CD2/CE1/NE2; Asp CB/CG/OD1/OD2) within each model and then across models. Dashed lines indicate equality; KL33 is a teal circle, KL30 a yellow circle and KL37 a teal star, with other designs in grey. One grey design lies outside both displayed ranges but is included in all counts. Peptide inclusion shortened the dyad distance in 47 of 79 designs and increased triad confidence in all 79, consistent with widespread peptide-dependent active-site organisation. KL37, the best-performing design in this set, showed pronounced reorganisation in the presence of the acceptor peptide, with the dyad shortening from 6.12 to 3.58 Å and a 19.02-point increase in triad pLDDT. KL30, which catalysed both ligation and hydrolysis, retained a short dyad in both states (3.86 and 3.75 Å) and gained 11.05 pLDDT points, approximately the 56th percentile of triad-pLDDT increases across the 79 designs (ascending rank 44/79). KL33, which also showed strong hydrolysis, had an already short dyad and little peptide-dependent change (3.72 to 3.81 Å), despite increased triad confidence **(C-E),** Top-ranked AF3 predictions of peptide-bound acyl-enzyme intermediates for KL33 (c), KL30 (d) and KL37 (e). Each structure is a single model; the enzyme is shown in a teal-to-green gradient, the acceptor peptide in peach and the covalently attached acyl group in purple.

**A**

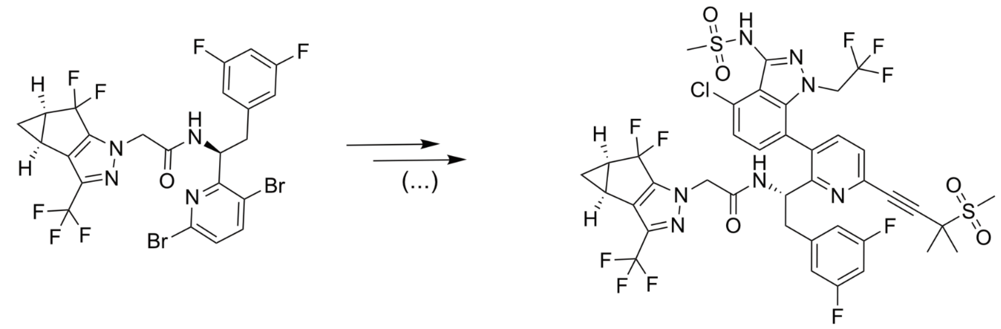

chiral amide (**3**) Lenacapavir

**B**

#### Amide bond formation (designed ligases)

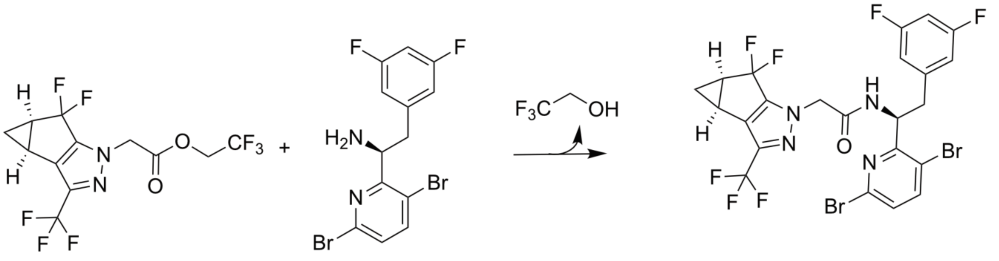

Ester (**1**) *(S)*-amine (**2**) chiral amide (**3**)

#### Hydrolysis

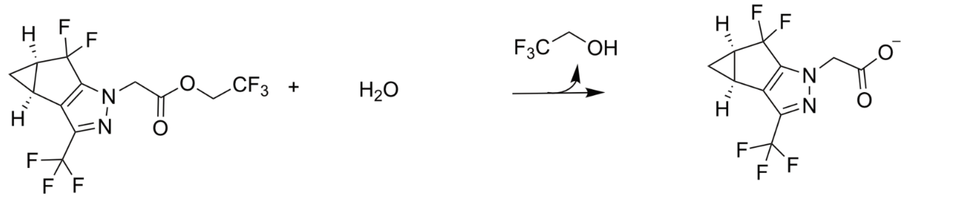

Ester (**1**) Acid (**4**)

**Supplementary Figure 7 | Extended reaction scheme for lenacapavir ligase. (A)** chiral amide (3) is an intermediate in a potential synthesis route to the HIV drug lenacapavir **(B)** Synthesis of lenacapavir intermediate **3** from ester **1** and amine **2** showing the amide bond formation as well as the hydrolytic side reaction to acid **4**.

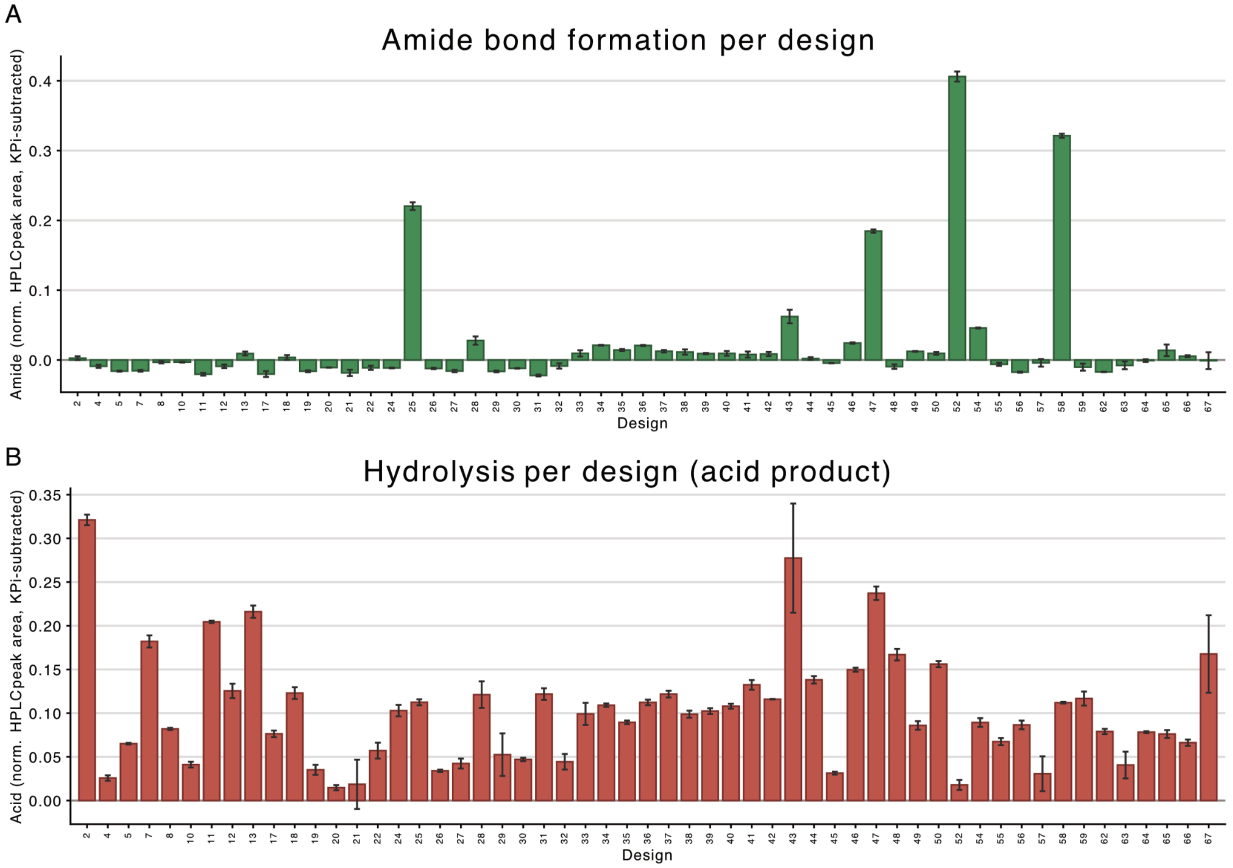

**Supplementary Figure 8 | Screening data for lenacapavir ligases.** Background-normalised HPLC peak areas of amide **3 (A)** and the donor hydrolysis product, acid **4 (B)**, across the 55 characterised designs. Reaction conditions: 50 µM enzyme, with 1 mM donor 1, and 0.5 mM amine 2 in 0.1 M PKi pH 6.0 and 20% DMSO, incubated for 16 h at 30°C.

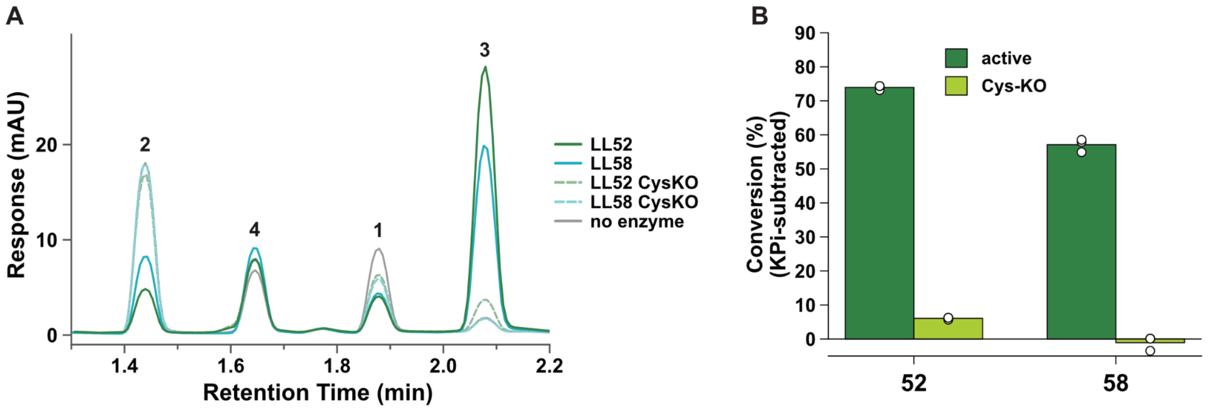

**Supplementary Figure 9 | Cysteine knock-outs of lenacapavir ligases. (A)** Raw HPLC traces of LL52 (teal) LL58 (cyan), LL52 cysteine knock out (teal dashed), LL58 cysteine knock out (cyan dashed) and no enzyme control (grey). **(B)** Conversion to amide product for cysteine knock-out variants after background subtraction showing a 12-fold reduction in activity for LL52 upon cysteine knockout and no measurable activity for LL58 over background after cysteine knock-out.

**A**

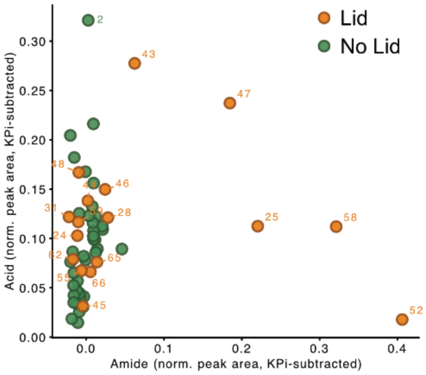

**B**

| Sequence(s) | Parent | Origin |
|---|---|---|
| 1, 2 | LL49 | Partial diffusion |
| 3 | LL51 | Partial diffusion |
| 4–22 | LL54 | Partial diffusion |
| 23, 24 | LL55 | Partial diffusion |
| 25 | LL58 | Partial diffusion |
| 26, 27 | LL61 | Partial diffusion |
| 28 | LL65 | Partial diffusion |
| 29, 30 | LL67 | Partial diffusion |
| 31 | LL52 | MPNN / AF3 cycling |
| 32–42 | LL63 | MPNN / AF3 cycling |
| 43–48 | LL66 | MPNN / AF3 cycling |

**C**

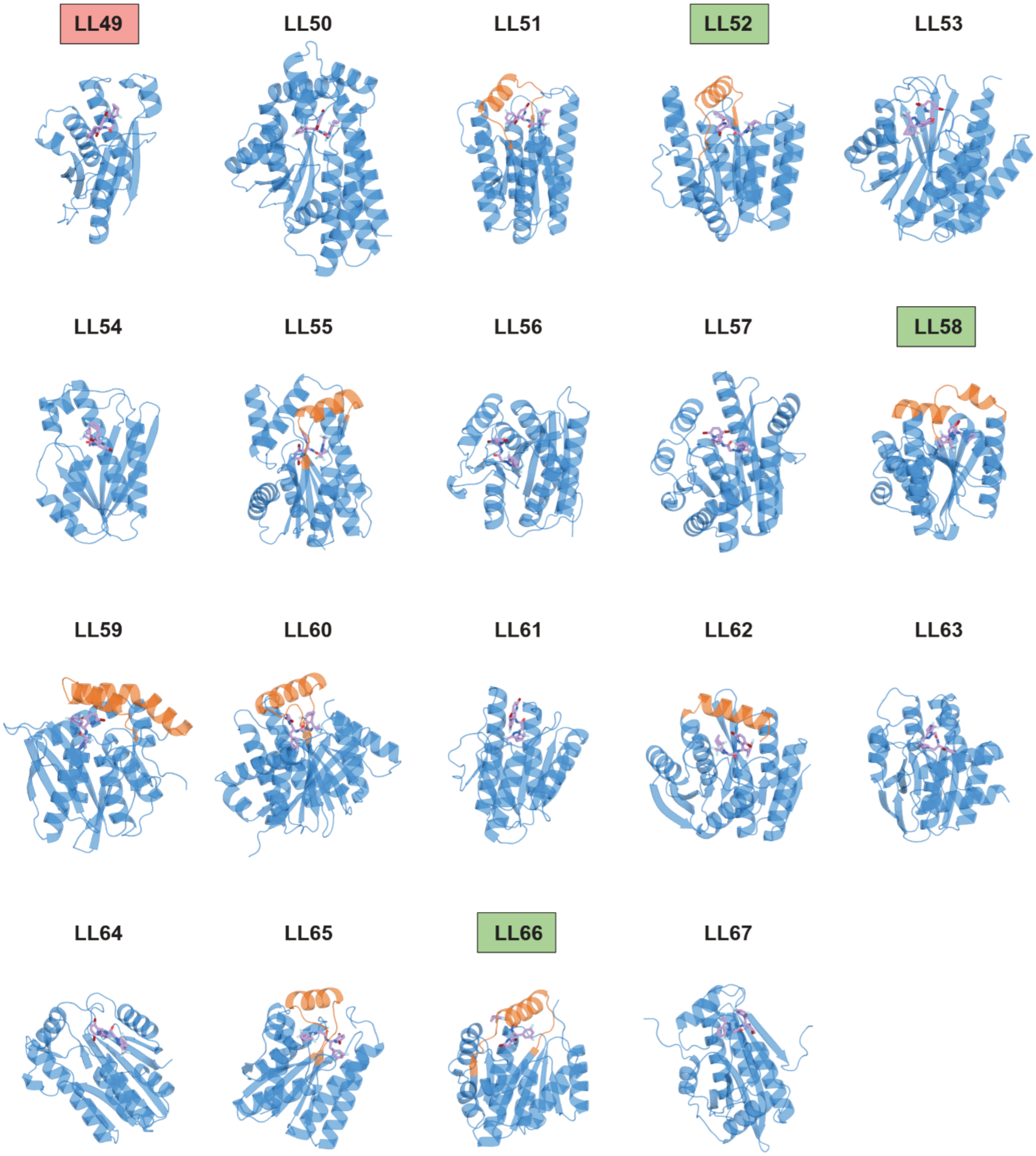

**Supplementary Figure 10 | RFD4-generated backbones with substantial ligase activity have moveable helical lids in common. (A)** Background corrected hydrolysis and amide bond formation activity for all tested designs **(B)** Table describing the starting backbone for each design generated by partial diffusion or MPNN/AF3 cycling **(C)** AF3 predictions of all RFD4-generated backbones (LL49-LL67) with bound product are shown. Lids attached to loops that could conceivably move over the ligand binding pocket or already cover it are shown in orange, and the rest of the structure in blue. Nine of 19 designed backbones have such lids. Among them are LL52 and LL58 (marked green), the top amide ligase hits of the campaign. Redesign of LL58 by partial diffusion gave LL25, which had slightly lower aminolysis activity but similar hydrolysis activity. Although the original LL66 design (marked green) was inactive, sequence redesign yielded LL47, which catalyses both aminolysis and (substantial) hydrolysis. Similarly, inactive LL49 (red) was sequence-redesigned by partial diffusion to give LL2, the top hit for hydrolysis. The other lid-containing designs are either catalytically inactive or show limited hydrolytic activity with the exemption of LL43 which shows low amide bond formation along with hydrolytic activity. It is derived from the same backbone as LL47. This shows that lids are a feature of designs with substantial ligase activity but that there are other sequence-specific features that allow hydrolytic activity in lid-containing designs.

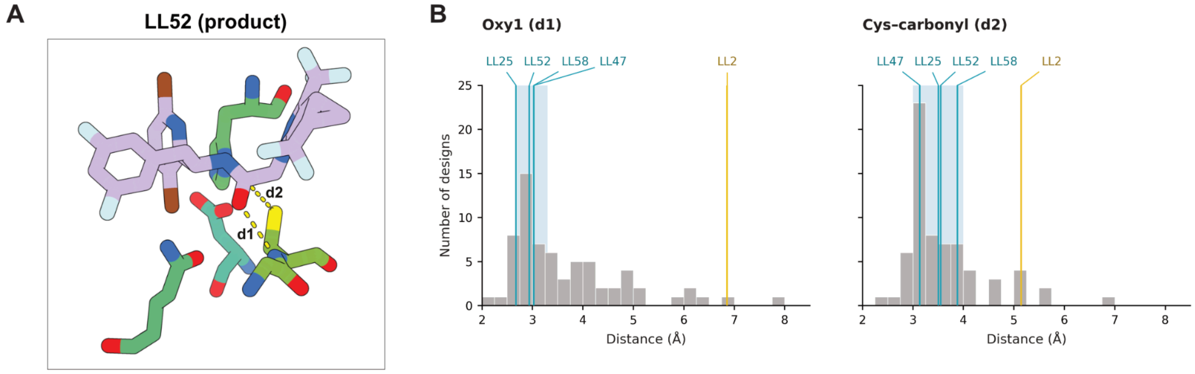

**Supplementary Figure 11 | Predicted product-state geometry of lenacapavir ligase designs. (A)** Close-up of the LL52 binding pocket with bound amide **3.** d1 (Oxy1) is the distance between the amide backbone nitrogen of the catalytic cysteine and the carbonyl oxygen of amide 3 (O29); d2 (Cys–carbonyl) is the distance between the gamma sulfur of the catalytic cysteine and the carbonyl carbon of amide 3 (C28). is the Cys Sγ–product carbonyl C28 distance. **(B)** Predicted product-state geometry of lenacapavir ligase designs. Histograms show the number of designs with specific d1 and d2 values.. Grey bars comprise all 67 designs, including the five highlighted designs, binned at 0.25 Å. Each distance was minimised independently across ten AlphaFold3 product-state predictions per design, so the two minima may derive from different predictions. Labeled teal lines indicate the four ligase hits LL25, LL47, LL52 and LL58; the yellow line shows LL2, which catalysed donor hydrolysis without detectable amide formation. Shaded regions mark the design target ranges (2.7–3.3 Å for d1 and 3.0–4.0 Å for d2). Retrospective filtering for d1 < 3.5 Å and d2 < 4.0 Å retained all four ligase hits among 32 of the 55 experimentally tested designs, increasing the hit fraction from 4/55 (7.3%) to 4/32 (12.5%), a 1.72-fold enrichment with 100% recall. The 12 designs without experimental readouts were excluded from enrichment calculations.

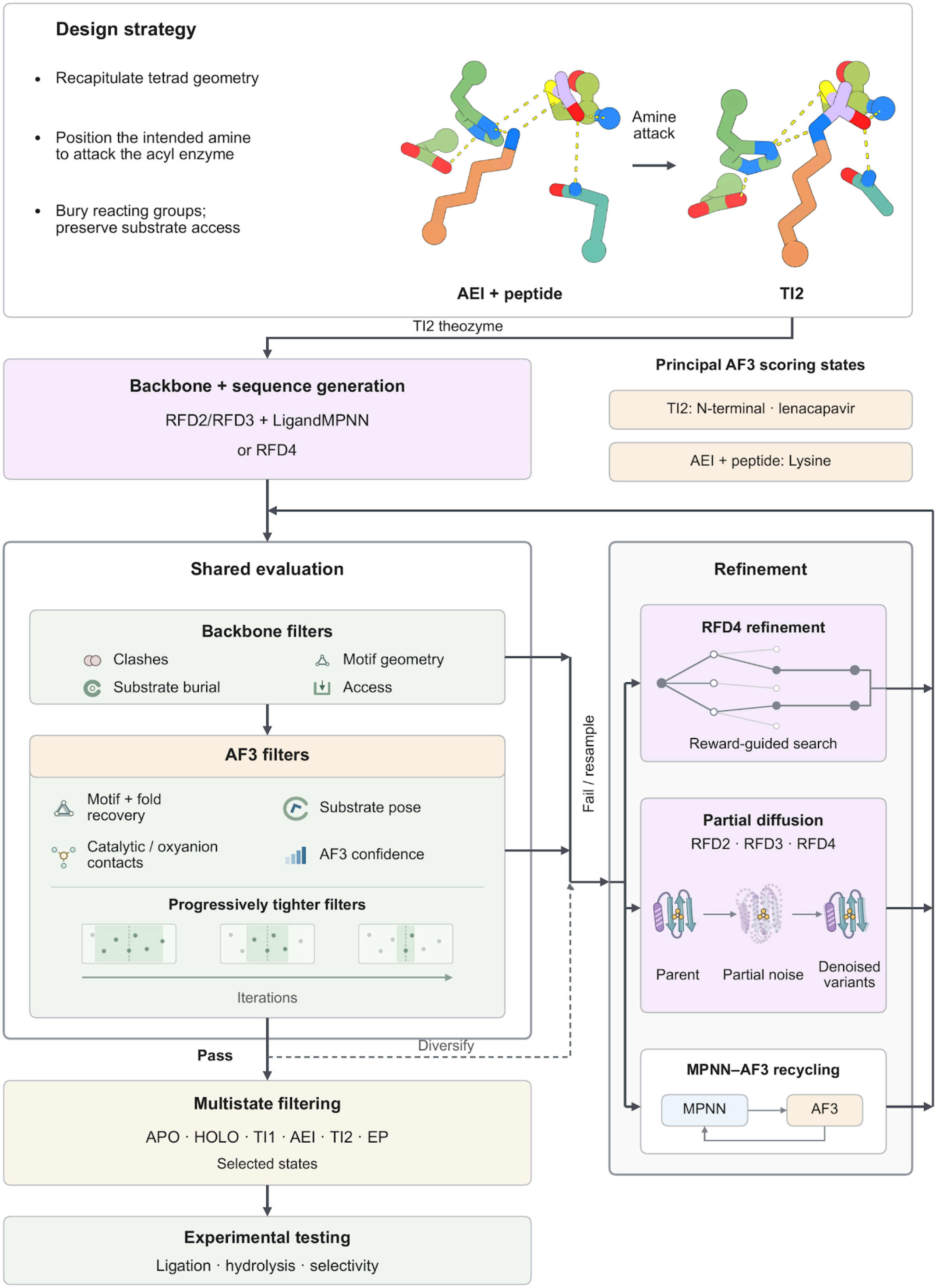

Design strategy
Recapitulate tetrad geometry
Position the intended amine to attack the acyl enzyme
Bury reacting groups; preserve substrate access
Amine attack
AEI + peptide
TI2
TI2 theozyme
Backbone + sequence generation
RFD2/RFD3 + LigandMPNN
or RFD4
Principal AF3 scoring states
TI2: N-terminal · lenacapavir
AEI + peptide: Lysine
Shared evaluation
Backbone filters
Clashes
Motif geometry
Substrate burial
Access
AF3 filters
Motif + fold recovery
Substrate pose
Catalytic / oxyanion contacts
AF3 confidence
Progressively tighter filters
Iterations
Fail / resample
Refinement
RFD4 refinement
Reward-guided search
Partial diffusion
RFD2 · RFD3 · RFD4
Parent
Partial noise
Denoised variants
MPNN–AF3 recycling
MPNN
AF3
Pass
Diversify
Multistate filtering
APO · HOLO · TI1 · AEI · TI2 · EP
Selected states
Experimental testing
Ligation · hydrolysis · selectivity

**Supplementary Figure 12 | General computational workflow for ligase design.** We sought to retain catalytic geometry, position the intended amine for attack on the acyl-enzyme intermediate (AEI), and bury reacting groups while preserving substrate access. Structural models illustrate AEI plus peptide and the second tetrahedral intermediate (TI2). Starting from a TI2 theozyme, we generated backbones with RFD2/RFD3 followed by LigandMPNN sequence design, or jointly generated sequence and structure with RFD4. We screened clashes, motif geometry, burial and access, then used AlphaFold3 (AF3) to assess fold, motif and substrate-pose recovery, catalytic and oxyanion contacts, and prediction confidence with progressively tighter filters. We used TI2 for principal AF3 scoring in the N-term ligase design campaign and Lenacapavir Ligase design campaign, and AEI plus peptide in the Lysine Ligase design campaign. Refinement through reward-guided RFD4 search, RFD2/RFD3/RFD4 partial diffusion or MPNN–AF3 recycling returned candidates to shared evaluation; we used these routes separately or together, and passing parents could seed diversification. We evaluated designs in selected subsets of the apo (APO), donor-bound (HOLO), first tetrahedral intermediate (TI1), AEI, TI2 and enzyme–product (EP) states before testing ligation, hydrolysis and selectivity experimentally. State subsets and thresholds differed by campaign.

### 2 Supplementary methods

#### 2.1 Computational design pipeline

##### 2.1.1 Overall design pipeline

We sought to construct active sites that recognise an acyl donor and an amine acceptor, position the intended amine for attack on a cysteine-bound acyl-enzyme intermediate (AEI), and favour aminolysis over competing hydrolysis. We retained a Cys–His–Asp catalytic triad to support acyl transfer and placed backbone or side-chain hydrogen-bond donors around the reacting carbonyl to stabilise the tetrahedral intermediates. To organise the second bond-forming step, we positioned the acceptor amine near both the acyl carbonyl carbon and the catalytic histidine. We hypothesised that burying the reacting groups within complementary donor- and acceptor-binding pockets could limit water access and favour productive aminolysis while retaining openings for the substrates. This structural hypothesis determined the theozyme coordinates, the burial and opening criteria applied to generated scaffolds, and the catalytic contacts and substrate geometries evaluated by structure prediction.

We applied this strategy to three targets. In the N-term ligase design campaign, we positioned the N-terminal α-amine of FSLTK through strand-pairing interactions, with its C-terminal lysine providing a competing amine. In the Lysine Ligase design campaign, we co-designed the enzyme and a helical peptide to position a specified lysine ε-amine for acylation. In the Lenacapavir Ligase design campaign, we targeted stereoselective coupling of an activated ester and a chiral small-molecule amine to form an amide intermediate in lenacapavir synthesis.

We conducted the N-term ligase design campaign as an exploratory test of structural criteria for ligation, testing candidates selected in the second tetrahedral intermediate (TI2) alone alongside candidates that passed filters in additional reaction states. All ten ligase hits came from the multistate-filtered cohort, and retrospective AlphaFold3 (AF3) analysis showed narrower distributions of acceptor-amine placement for ligases than for hydrolases. These observations motivated stricter selection of amine placement relative to the AEI carbonyl and catalytic histidine, together with evaluation in additional reaction states, in the subsequent campaigns. We implemented the workflow as successive stages of theozyme construction, backbone and sequence generation, backbone filtering, AF3 evaluation, refinement and selected-state filtering before experimental testing (Supplementary Fig. 12). At each refinement stage, we returned new candidates through structural evaluation and retained those meeting the current criteria for further refinement or final selection. The following sections describe these shared operations, followed by the models, settings and state-specific criteria for each campaign in sections 2.1.2–2.1.4.

#### Theozyme construction

We constructed each theozyme around TI2, which forms upon amine attack on the AEI. This representation places the cysteine sulfur and acceptor nitrogen on the same tetrahedral acyl centre and specifies the oxyanion contacts required for the intended bond-forming step. We used RDKit[3] to generate the small-molecule components and sample alternative conformers, then relaxed the conformers by energy minimisation with molecular force fields such as OpenFF Sage 2.2.1. We assembled the substrates in the idealised TI2 geometry within a cysteine-protease-derived catalytic site, retaining the Cys–His–Asp arrangement and positioning backbone or side-chain hydrogen-bond donors around the oxyanion. We included selected neighbouring backbone atoms to specify the local architecture supporting the catalytic residues and the intended acceptor contacts. For peptide acceptors, we extended the reacting residue into the chosen backbone conformation to define the substrate-binding surface. We passed the resulting atomic coordinates, covalent connectivity and fixed-atom selections to scaffold generation as constraints on the catalytic site and substrate placement.

#### Backbone and sequence generation

We used RFdiffusion2 (RFD2) or RFdiffusion3 (RFD3) to generate backbones around the theozyme, retaining the specified catalytic coordinates and substrate context while sampling the surrounding protein. We then used LigandMPNN to generate sequences for retained backbones, fixing catalytic identities and including substrates as structural context. Where we allowed the acceptor sequence to vary, we sampled enzyme and peptide sequences together while preserving the reacting residue. Alternatively, we used RFD4-Proteina to generate structure and sequence jointly around the same type of theozyme input. These routes produced candidate enzyme–substrate complexes for evaluation; we could apply backbone-only criteria before sequence design to avoid further sampling of unsuitable scaffolds.

#### Backbone evaluation

We first checked generated structures for clashes, chain integrity, retention of the catalytic motif and the local backbone conformations required to support it. We evaluated substrate burial and access to retain pockets that enclosed the reacting groups while leaving an opening to the substrates. We advanced candidates passing the relevant backbone criteria to sequence design or structure prediction and used retained parents for further scaffold sampling or refinement when candidates failed. We chose the criteria and numerical cutoffs for each campaign by visually inspecting the resulting designs.

#### Substrate burial and access

We calculated relative solvent-accessible surface area (RASA) as $SASA_{complex}/SASA_{isolated}$, clipped to [0, 1]. For each component, we summed its accessible surface area in the full complex to obtain $SASA_{complex}$ and repeated the calculation at the same coordinates after removing all other components to obtain $SASA_{isolated}$. We defined burial as $1 - \text{RASA}$, ranging from 0 for unchanged exposure to 1 for complete loss of accessible surface. This calculation accounted for occlusion by all other components in the complex.

We used peptide and ligand opening angles in both peptide-ligase campaigns to measure the largest unoccupied angular sector around each component. For the peptide, we defined the longitudinal axis as the first principal component of its mean-centred backbone N, Cα and C coordinates, with their centroid as the origin. We projected surrounding enzyme backbone N, Cα and C atoms onto the plane perpendicular to this axis. For the ligand, we defined the axis using a designated pair of ligand atoms, with their midpoint as the origin, and projected enzyme heavy atoms onto the perpendicular plane, excluding peptide and ligand atoms. For each projection, we sorted atom azimuths and took the largest gap between consecutive angles, including the last-to-first interval across 360°, as the opening angle. We used no radial or axial cutoffs in either calculation. Burial and opening angles provided geometric proxies for enclosure and access; they did not measure water exclusion, substrate binding or product release.

To combine burial with angular clearance, we defined the ligand pocket score as ligand burial × min(ligand opening angle, 180°)/180°. The opening contribution therefore saturated at 180°. We chose burial and access thresholds by visually inspecting the resulting designs. We specify atom selections, cutoffs and the stages at which we applied these criteria within each campaign below.

#### AF3 prediction and structural evaluation

We evaluated candidate sequences with AF3 in a principal scoring state: TI2 in the N-term ligase design campaign and the Lenacapavir Ligase design campaign, and AEI plus peptide in the Lysine Ligase design campaign. For each state, we specified the enzyme, the appropriate substrates and any covalent modifications. We used custom chemical component dictionary (CCD) definitions to encode noncanonical residues, ligand chemistry and internal bond orders, and supplied explicit intercomponent bonds where required. We represented covalent intermediates as modifications of the catalytic cysteine and supplied noncovalently bound substrates as separate components. For apo predictions, we used the unmodified enzyme; for donor-bound predictions, we included a separate activated ester with the unmodified enzyme. Thus, the TI2 representation specified the formed C–N bond, whereas the AEI-plus-acceptor representation allowed us to assess whether a separate amine recovered the intended approach to the acyl carbonyl.

We compared predicted structures with the design models for recovery of the enzyme fold, catalytic motif and substrate pose. We also evaluated catalytic-residue contacts, oxyanion-hole geometry, acceptor placement and AF3 confidence. In the AEI, we assessed amine-attack geometry through acceptor contacts with the acyl carbonyl carbon and catalytic histidine, and through approach-angle criteria where used, retaining the campaign-specific distinction between hard filters and ranking metrics. Together, these measurements tested whether the designed sequence recovered both the scaffold and the local geometry required for the intended reaction state. This stage produced predicted structures with associated geometric and confidence scores. We advanced passing designs to selected-state evaluation or used them as parents for refinement, then re-evaluated their descendants through the same backbone and AF3 filtering stages.

#### Refinement and re-evaluation

We used three refinement routes separately or in combination, according to the campaign. In reward-guided RFD4 search, we evaluated generated candidates against structural objectives and used favourable candidates to guide further sampling. Where we used beam search, rewards incorporating designed and AF3-predicted catalytic geometry selected parents for further diversification.

For partial diffusion, we started from a retained structure, introduced a limited amount of noise and denoised it while preserving the specified catalytic and substrate geometry. We returned the resulting backbone variants to backbone filtering, sequence design where required, and AF3 evaluation. This route diversified scaffolds around a parent that had already satisfied the preceding selection criteria.

For MPNN–AF3 cycling, we used passing predicted structures as the coordinates for sequence redesign, followed by another round of prediction and filtering. Before MPNN redesign, we converted modified cysteines to standard cysteine plus separate ligand components without changing atomic coordinates. Where we included constrained relaxation, we maintained the target catalytic interactions during sequence refinement. We used passing descendants as inputs for subsequent rounds and progressively tightened filters to select improved agreement with the intended fold, catalytic geometry and substrate placement. We returned each new candidate through structural evaluation before further refinement or selected-state filtering. We specify the refinement routes and number of rounds for each campaign below.

##### Selected-state filtering and experimental testing

We predicted sequences retained after refinement in additional selected states to assess compatibility with other steps of the reaction pathway. The available states comprised apo enzyme (APO), donor-bound enzyme (HOLO), the first tetrahedral intermediate (TI1), AEI with acceptor, TI2 and enzyme–product complex (EP). These assessments tested whether candidates selected around one intermediate could also recover the fold, substrate placement and catalytic contacts appropriate to other states. We applied state-specific geometric and confidence criteria, distinguishing hard acceptance thresholds from metrics used only for ranking. For each campaign, we evaluated the selected set of states in the order and with the advancement criteria specified below. We then ranked and selected candidates for experimental measurements of ligation, donor hydrolysis and selectivity.

##### Retrospective AF3 ensemble analysis

We retrospectively analysed independent AF3 predictions to examine how consistently the designs recovered catalytic and substrate geometry and whether this consistency differed with measured activity. We kept the molecular composition, chemical definitions and covalent connectivity fixed within each state, while varying the prediction samples. For comparisons with and without an acceptor, we retained the same enzyme sequence and acyl modification and added or removed the acceptor component. We measured catalytic contacts, substrate placement, motif recovery and prediction confidence across the resulting ensembles and compared the relevant per-design summaries with the experimental activity classes. Depending on the analysis, we used distributions, means, variances or minimum distances, with the states, sample counts, atom selections and statistics specified in the campaign sections below.

We used these retrospective comparisons to examine acceptor-dependent active-site organisation and product accommodation, and to identify geometric features associated with ligation or donor hydrolysis. We interpreted ensemble variability as differences among AF3 predictions, not as thermodynamic sampling or measured conformational dynamics. These analyses were separate from prospective selection: the observations of acceptor-dependent organisation and product-state geometry informed mechanistic hypotheses and possible filters for future campaigns.

#### 2.1.2 N-term ligase design campaign

##### Theozyme and substrate geometry

We derived the catalytic motif from tobacco etch virus protease (PDB: 1LVM)[1], retaining the Cys–His–Asp triad and the three residues preceding the catalytic cysteine, including the oxyanion-binding backbone amides. To improve oxyanion-hole geometry, we partially diffused the oxyanion loop using RFdiffusion2[2] while holding the remaining catalytic motif fixed, and selected a conformation by visual inspection. We added an alanine with α-helical backbone geometry immediately after the cysteine to promote extension of the cysteine-containing helix.

We generated TI2 conformers comprising catalytic cysteine, Janelia Fluor 549 (JF549) and the N-terminal phenylalanine of the FSLTK acceptor with RDKit. We selected two conformers and positioned them within the catalytic site using PyMOL (Schrödinger, LLC). We built the remaining SLTK residues as an idealised β-strand and positioned polyalanine guide strands on either side to form backbone hydrogen bonds with the acceptor. Inputs contained no guide strand, one strand or two strands in parallel, antiparallel or mixed orientations. We partially diffused these theozymes at a time of 10 steps while fixing the catalytic motif and acceptor phenylalanine before full scaffold generation.

#### Backbone generation, sequence design and refinement

RFD2 sampled scaffold lengths of 155–170 residues with randomised contig placement, retaining selected backbone and side-chain atoms of the catalytic motif, TI2 and acceptor peptide. The JF549-containing component had partially fixed coordinates and atomwise accessibility constraints to favour burial of the reactive centre and fluorophore; peptide-interaction hotspots guided formation of the acceptor-binding site. Backbone filters assessed steric clashes, loop content, maximum helix length, substrate burial and peptide and ligand opening angles as defined in section 2.1.1. We partially diffused passing backbones at partial_T = 40 while preserving the catalytic motif, ligand and acceptor geometry.

LigandMPNN[4] sampled only the enzyme sequence at temperature 0.1, with ligand-aware side-chain context and the JF549-containing ligand and FSLTK peptide present. We fixed the catalytic triad and FSLTK sequence and excluded additional cysteines at designable positions. Initial AlphaFold3[5] predictions used one randomised seed and five diffusion samples, with empty multiple-sequence alignments and no protein templates. TI2 inputs connected the modified catalytic cysteine, a separate N-terminal phenylalanine component and the remaining SLTK peptide through the required covalent bonds. After initial TI2 filtering, we partially diffused selected parent backbones at partial_T = 30 and returned them to sequence design and prediction. Subsequent MPNN–AF3 rounds used the same temperature and residue restrictions with progressively more stringent confidence, motif-recovery and active-site geometry criteria.

#### Multistate filtering and selection

We evaluated sequences generated from structures passing the final TI2 filters sequentially in APO, HOLO, TI1, AEI plus peptide, TI2 and EP, advancing them only after they passed the current state's criteria (Supplementary Table 1). Each state used one randomised seed, with five diffusion samples for TI1 and TI2 and one for each remaining state. The table gives the final sequential multistate criteria; it does not specify the progressively tightened gates used during the preceding TI2 refinement rounds. We selected a separate experimental cohort from the highest-scoring candidates in the final TI2-filtered pool. Ranking combined catalytic distances, oxyanion-hole geometry, motif RMSD, backbone dihedrals and confidence across the evaluated states. We scored individual metrics relative to thresholds, assigning positive scores to favourable values, exponentially increasing penalties to unfavourable values and $-10^6$ to missing metrics. The composite score was the arithmetic mean of the metric scores.

Supplementary Table 1 | Sequential multistate filters for N-term ligase design. We required all criteria within a state, using inclusive inequalities; dashes indicate metrics without a threshold. Distances and RMSDs are in Å, and backbone dihedrals and oxyanion angles are in degrees. APO, unmodified enzyme; HOLO, donor-bound enzyme; AEI + peptide, acyl-enzyme intermediate with acceptor; EP, enzyme–product complex. TI1 and TI2 denote the first and second tetrahedral intermediates. pLDDT, predicted local distance difference test; pTM, predicted template modelling score; ipTM, interface predicted template modelling score. Input motif labels A1, A2 and A6 correspond to His, Asp and catalytic Cys, and A4 and A5 to the residues two and one positions before Cys, respectively. Modified-Cys confidence refers to the represented chemical component, which can include covalently attached ligand atoms. We retain operational metric aliases in parentheses to identify the original filters.

| **Criterion** | **APO** | **HOLO** | **TI1** | **AEI + peptide** | **TI2** | **EP** |
|---|---|---|---|---|---|---|
| Enzyme pLDDT (plddt_chain_A_mean) | ≥75 | ≥75 | ≥75 | ≥75 | ≥90 | ≥75 |
| Catalytic-motif RMSD (rmsd_motif_mean) | ≤1 | ≤1 | ≤1 | ≤1 | ≤1 | — |
| ipTM | — | ≥0.7 | — | — | — | — |
| pTM | — | ≥0.7 | ≥0.7 | ≥0.7 | ≥0.9 | ≥0.7 |
| Modified-Cys pLDDT (plddt_A6_mean) | — | — | ≥70 | — | ≥90 | — |
| Peptide-chain pLDDT (plddt_chain_B_mean) | — | — | — | ≥75 | ≥82 | — |
| Histidine contact (LG_HIS_dist_mean) | — | — | — | ≤4 | ≤4 | — |
| Peptide end-to-end metric (pep_N_to_C_dist_mean) | — | — | — | — | ≥15 | — |
| Histidine contact (Nu_HIS_dist_mean) | — | — | — | — | ≤4 | — |
| Backbone oxyanion contact (Oxy_bb_dist_mean) | — | — | — | — | ≤3.5 | — |
| Second backbone oxyanion contact (oxh2_bb_dist_mean) | — | — | — | — | ≤3.7 | — |
| Acid–histidine contact (Acid_His_dist_mean) | — | — | — | — | ≤3.5 | — |
| Cys−2 backbone φ (oxh2_phi_A4_mean) | — | — | — | — | ≥−20 | — |
| Cys−1 backbone φ (oxh2_phi_A5_mean) | — | — | — | — | ≤−50 | — |
| Catalytic-Cys backbone φ (oxh2_phi_A6_mean) | — | — | — | — | ≤−40 | — |
| Second oxyanion donor angle (oxh2_donor_angle_mean) | — | — | — | — | ≥90 | — |
| Second oxyanion acceptor angle (oxh2_acceptor_angle_mean) | — | — | — | — | ≥82 | — |
| Catalytic-His pLDDT (plddt_A1_mean) | — | — | — | — | ≥90 | — |
| Catalytic-Asp pLDDT (plddt_A2_mean) | — | — | — | — | ≥90 | — |
| Cys−2 pLDDT (plddt_A4_mean) | — | — | — | — | ≥90 | — |
| Cys−1 pLDDT (plddt_A5_mean) | — | — | — | — | ≥90 | — |

#### Retrospective AF3 ensemble analysis

We analysed 50 AF3 predictions per design and state for the 96 screened designs, covering apo enzyme and AEI with and without acceptor. For NL6, we compared the predicted structures across these states (Fig. 3A) and measured distances from the acyl carbonyl oxygen to the two backbone amide nitrogens forming OXH1 and OXH2 in the acceptor-free and acceptor-bound AEI ensembles (Fig. 3B). Across the

96 designs, we compared experimentally assigned aminolysis-active, hydrolysis-only and inactive groups using per-design means and variances of the acceptor α-amine distances to the acyl carbonyl carbon and catalytic His Nε2 in acceptor-bound AEI predictions (Supplementary Fig. 3A,B). We calculated motif RMSDs after fitting selected catalytic side-chain and backbone atoms to the original motif, comprising Cys, His, Asp and the two oxyanion-loop residues preceding cysteine. For each state, we compared variances across predictions for the complete motif, catalytic triad and oxyanion loop (Supplementary Fig. 3C).

#### 2.1.3 Lysine ligase design campaign

We co-designed enzymes and 19-residue helical peptides to position a specified lysine ε-amine for attack on a cysteine-bound acyl intermediate. The campaign used RFdiffusion3 (RFD3)[6] scaffolding, four rounds of sequence design and AF3 evaluation, and one intervening round of partial diffusion (Supplementary Fig. 12).

##### Theozyme and molecular representations

The Ulp1-derived motif (PDB: 1EUV)[7] contained the Cys–His–Asp triad and an oxyanion hole formed by glutamine and the cysteine backbone amide. Backbone atoms surrounding cysteine specified its local conformation, and the histidine backbone retained the orientation of a neighbouring carbonyl proposed to help position the lysine amine. Reference motif labels A3, A8, A12 and A14 denoted Cys, His, Asp and Gln, respectively (Supplementary Table 2). Chain A denoted the enzyme and chain B the peptide. We extended the 15-residue α-helical acceptor segment by two residues at each terminus, placing the reacting reference lysine B8 at position 10 of the designed 19-residue peptide. The initial scaffolding retained two reference peptide glutamines, whereas subsequent sequence design fixed only the reacting lysine within the peptide.

AF3 AEI inputs represented the 2-(4-ethynylphenyl)acetyl group as a cysteine thioester and the acceptor as a separate peptide chain. In the geometric definitions, C7 and O11 denote the acyl carbonyl carbon and oxygen, and O1 denotes the ester oxygen attached to the trifluoroethyl leaving group in the donor-bound state. C6 and C9 denote the benzylic and terminal ethynyl carbons used to define the ligand opening axis.

Supplementary Table 2 | Motif atoms fixed during RFD3 scaffolding and used for catalytic-motif RMSD. Backbone atoms comprise N, Cα, C and O; residue labels refer to the input motif.

| Reference residue | Fixed coordinates during scaffolding | Atoms used for motif RMSD |
|---|---|---|
| A2 | Backbone | Backbone |
| A3, Cys | Backbone, Cβ, Sγ | Backbone, Cβ, Sγ |
| A4 | Backbone | Backbone |
| A5, A9, A10, A11, A13 | Backbone | None |
| A8, His | Backbone, Cβ, Cγ, Nδ1, Cδ2, Cε1, Nε2 | Cβ, Cγ, Nδ1, Cδ2, Cε1, Nε2 |
| A12, Asp | Backbone, Cβ, Cγ, Oδ1, Oδ2 | Backbone, Cβ, Cγ, Oδ1, Oδ2 |
| A14, Gln | Cγ, Cδ, Oε1, Nε2 | Cγ, Cδ, Oε1, Nε2 |

#### Backbone generation and selection

RFD3 scaffolding used a total-length parameter of 175–275 residues, with two generated peptide residues on either side of reference B1–B15. We allowed amino-acid identities to vary at enzyme positions for which Supplementary Table 2 specifies only backbone coordinates. We perturbed alternative generation centres independently by uniform displacements of −4 to +4 Å along each Cartesian axis. We sampled scaffolds with and without ligand accessibility conditioning; for conditioned inputs, we assigned a value of 1 to all 11 acyl-group heavy atoms, comprising ten carbons and O11. This value was a model-conditioning parameter rather than a measured accessible-surface fraction. We generated scaffolds with randomised seeds, 200 diffusion steps, origin-jitter parameter 3.0, step scale 2 and $\gamma_0$ = 0.8, and removed virtual atoms and guideposts before evaluation.

We selected backbones using the criteria in Supplementary Table 3. We classified DSSP assignments H, E/B and all others as helix (a), strand (b) and loop (c), respectively, and calculated the enzyme non-loop fraction as the combined helix and strand fraction. We required three-residue windows centred on reference A2, A3, A8 and A12 to match {cca, ccc}, {cca, caa}, {cbb, ccb, bbb, bbc, bcc} and {cbb, ccb, bbb, bbc, bcc}, respectively, and retained only backbones passing all four windows. Burial calculations used all peptide-chain atoms and all HETATM-labelled acyl-group atoms. We calculated SASA with the Shrake–Rupley implementation in Biotite[8], element-based Single radii, a 1.4 Å probe and 1,000 Fibonacci-distributed surface points per atom. Peptide and ligand opening angles followed section 2.1.1, with the ligand axis joining C6 and C9. Ligand opening contributed to selection through the pocket score.

Supplementary Table 3 | Backbone-selection criteria. We required all criteria within a stage, using inclusive inequalities. “Not applied” indicates that the criterion was not part of that stage’s filter.

| Criterion | Initial scaffolds | Partial-diffusion scaffolds |
|---|---|---|
| RFD3-reported ligand clashes | 0 | Not applied |
| RFD3-reported chain breaks | 0 | Not applied |
| Four motif secondary-structure windows | All pass | All pass |
| Enzyme helix/strand fraction | ≥0.65 | ≥0.65 |
| Ligand burial | ≥0.75 | Not applied |
| Peptide burial | ≥0.40 | Not applied |
| Peptide opening angle | ≥40° | ≥40° |
| Ligand pocket score | ≥0.06 | ≥0.1 |

#### Initial sequence design and AF3 evaluation

LigandMPNN co-designed enzyme and peptide sequences at temperature 0.1 with the catalytic triad, oxyanion-hole glutamine and reacting lysine fixed, excluding additional cysteines at designable positions. We supplied empty paired and unpaired multiple-sequence alignments and no protein templates to AF3, and calculated structural metrics from the highest-ranked sample. Round 1 first required AEI-plus-peptide enzyme Cα RMSD ≤2 Å, catalytic-motif RMSD ≤1.5 Å and peptide RMSD ≤2 Å. We predicted passing candidates in the apo state and applied the tighter combined AEI/apo criteria in Supplementary Table 5. Enzyme Cα RMSD followed alignment to the corresponding sequence-design model, and motif RMSD followed fitting of the 32 atoms in Supplementary Table 2 to the input motif. Peptide RMSD used N, Cα,

C and O of reference B6–B10, corresponding to designed residues 8–12, after enzyme Cα alignment to measure peptide placement relative to the enzyme.

#### Partial diffusion and constrained refinement

After round 1, backbone family selection retained the trajectory with the lowest AEI-state peptide RMSD from each original RFD3 backbone family. The original generated coordinates associated with that trajectory, rather than its AF3 coordinates, supplied the input for partial diffusion at times 1, 5, 10 and 15. We retained the catalytic motif, reference peptide segment B1–B15 and ligand, and set ligand accessibility conditioning to 1. We screened partial-diffusion backbones with the same motif secondary-structure, enzyme helix/strand and peptide-opening criteria, increased the ligand pocket-score threshold to ≥0.1 and omitted the separate burial requirements (Supplementary Table 3).

Round 2 combined LigandMPNN sampling at temperature 0.1 with side-chain packing and constrained Rosetta relaxation; every relaxed candidate proceeded to AF3 evaluation. Relaxation used one repacking-only FastDesign repeat, beta weights, MonomerDesign2019 and non-Cartesian dfpmin_armijo_nonmonotone minimisation. We enabled backbone, side-chain and rigid-body movements except for the catalytic cysteine backbone. Distance, angle and torsion constraints maintained the seven interactions in Supplementary Table 4 with constraint score weights of 1.0. These were relaxation targets, separate from AF3 acceptance criteria.

Supplementary Table 4 | Geometric targets during constrained relaxation. For atom frames A = (a1, a2, a3) and B = (b1, b2, b3), d = a1–b1, α = a2–a1–b1, β = a1–b1–b2, $\tau_A$ = a3–a2–a1–b1, $\tau_{AB}$ = a2–a1–b1–b2 and $\tau_B$ = a1–b1–b2–b3. Distances are in Å and angles and torsions in degrees. Distance terms used a tolerance of 0.1 Å and force-constant parameter of 100; angle and torsion terms used a tolerance of 5° and force-constant parameter of 50, with periodicity 360°. We designated the C7–Cys Sγ interaction as covalent.

| **Atom frame A** | **Atom frame B** | **d** | **α** | **β** | **$\tau_A$** | **$\tau_{AB}$** | **$\tau_B$** |
|---|---|---|---|---|---|---|---|
| Acyl C7, O11, C6 | Cys Sγ, Cβ, Cα | 1.7 | 121.6 | 111.5 | −179.7 | −1.9 | −90.9 |
| Acyl O11, C7, C6 | Cys N, Cα, C | 3.3 | 87.0 | 99.0 | −136.3 | −68.4 | 130.4 |
| Acyl O11, C7, C6 | Gln Nε2, Cδ, Oε1 | 2.5 | 157.0 | 129.8 | −164.9 | −21.5 | −15.4 |
| Cys Sγ, Cβ, Cα | His Nδ1, Cε1, Nε2 | 3.8 | 63.3 | 108.7 | −169.5 | −2.4 | −143.2 |
| Lys Nζ, Cε, Cδ | His Nδ1, Cε1, Nε2 | 2.9 | 97.5 | 136.9 | −83.8 | −76.2 | 167.9 |
| His Nε2, Cε1, Nδ1 | Asp Oδ1, Cγ, Oδ2 | 2.6 | 128.8 | 120.4 | −170.9 | 177.8 | −42.6 |
| Lys Nζ, Cε, Cδ | Cys Sγ, Cβ, Cα | 2.9 | 148.1 | 99.3 | −173.4 | 53.3 | −137.0 |

Round-2 AEI filtering tightened the RMSD limits and added ≤3.5 Å limits for Lys Nζ–C7 and His Nδ1–Lys Nζ before apo evaluation (Supplementary Table 5). Passing AEI structures supplied the coordinates for round 3 after conversion of the modified cysteine to standard cysteine plus a separate acyl component, preserving coordinates and the peptide chain. Round 3 repeated LigandMPNN sampling and constrained relaxation with the same residue restrictions and catalytic constraints as round 2. AEI filtering added Cys–His and acyl-oxygen–cysteine-backbone contact requirements, followed by apo filtering and donor-bound prediction. Round 4 used passing round-3 AEI structures for EnhancedMPNN sampling and constrained relaxation, retaining the temperature, fixed identities and constraints. We retained the round-3 AEI and apo thresholds and used donor-bound predictions for ranking without a separate hard filter.

Supplementary Table 5 | AF3 structural and geometric selection criteria. Round-1 values describe the combined acyl/apo screen after the initial acyl prescreen. In rounds 2–4, we applied acyl criteria before apo prediction, then required candidates to meet the apo criteria. All distances and RMSDs are in Å, and inequalities are inclusive.

| Criterion | Round 1 | Round 2 | Rounds 3 and 4 |
|---|---|---|---|
| Acyl enzyme Cα RMSD | ≤2.0 | ≤1.5 | ≤1.5 |
| Acyl catalytic-motif RMSD | ≤1.25 | ≤1.0 | ≤1.0 |
| Acyl peptide RMSD | ≤1.0 | ≤1.0 | ≤1.0 |
| Acyl Lys Nζ–C7 | Not applied | ≤3.5 | ≤3.5 |
| Acyl His Nδ1–Lys Nζ | Not applied | ≤3.5 | ≤3.5 |
| Acyl Cys Sγ–His Nδ1 | Not applied | Not applied | ≤4.0 |
| Acyl O11–Cys N | Not applied | Not applied | ≤3.5 |
| Apo enzyme Cα RMSD | ≤2.0 | ≤1.5 | ≤1.5 |
| Apo catalytic-motif RMSD | ≤1.25 | ≤1.0 | ≤1.0 |

#### Ranking and final selection

We ranked round-3 and round-4 designs using the weighted targets in Supplementary Table 6. For each lower-is-better metric x with target T, the contribution was zero at $x \leq T$ and $-\exp((x - T)/T)$ otherwise. The Lys Nζ–C7–O11 attack-angle contribution was zero at ≥90° and $-\exp((90 - \text{angle})/10)$ below 90°. We assigned $-10^6$ to missing values and divided the weighted sum by 15, preferring higher values. The chain-pair PAE summary was the minimum of the two directional off-diagonal entries in AF3's chain-pair-minimum-PAE matrix, summarising enzyme–peptide uncertainty in AEI and enzyme–donor uncertainty in the donor-bound state. We combined composite ranking with shortlisting within original-backbone families to retain variants from multiple scaffolds.

Supplementary Table 6 | Soft ranking targets and weights.

| Metric | Preferred value | Weight |
|---|---|---|
| Holo catalytic-motif RMSD | ≤1.0 Å | 1 |
| Holo His Nδ1–donor O1 | ≤3.0 Å | 3 |
| Holo donor C7–Cys Sγ | ≤3.0 Å | 3 |
| Holo Cys Sγ–His Nδ1 | ≤4.0 Å | 3 |
| Holo chain-pair PAE summary | ≤1.5 Å | 1 |
| Acyl chain-pair PAE summary | ≤1.5 Å | 2 |
| Acyl O11–Gln Nε2 | ≤3.5 Å | 1 |
| Acyl Lys Nζ–C7–O11 angle | ≥90° | 1 |

##### Retrospective AF3 ensemble analysis

To assess acceptor-dependent organisation across the 79 designs, we generated 50 AF3 predictions of the AEI with and without the cognate peptide, using ten seeds and five samples per seed (Supplementary Fig. 6). We retained all 7,900 predictions without filtering by confidence, activity, rank or geometry. For each design and state, we calculated the arithmetic mean Cys Sγ–His Nδ1 distance across models. We calculated triad confidence by averaging pLDDT equally over the 12 side-chain heavy atoms of Cys, His and Asp within each model, then averaging those values across models. The atom set comprised Cys Cβ/Sγ; His Cβ/Cγ/Nδ1/Cδ2/Cε1/Nε2; and Asp Cβ/Cγ/Oδ1/Oδ2. It excluded backbone atoms, oxyanion-hole glutamine, the acyl group and peptide. We compared the two state-specific means for each design and related the changes to measured ligation and donor hydrolysis. For the separate KL37 comparison in Fig. 3, we used 25 predictions per state and averaged active-site confidence over the 17 side-chain heavy atoms of Cys, His, Asp and Gln within each model.

#### 2.1.4 Lenacapavir ligase design campaign

##### Theozyme and RFD4 generation

We used a single Ulp1-derived theozyme throughout this campaign, comprising His514, Asp531, Gln574 and the Asp579–Cys580–Gly581 nucleophile elbow. The TI2 representation contained the intended acceptor stereochemistry and explicit Cys Sγ–C28 and C28–acceptor N43 bonds. AEI inputs instead contained the cysteine thioester and a separate, noncovalently bound amine.

The original RFD4-Proteina version required indexed noncanonical residues. We therefore indexed the remaining catalytic residues and evaluated all six sequence-order permutations of Asp, Gln and His (DQH, DHQ, QDH, QHD, HDQ and HQD), retaining the nucleophile elbow in fixed order at the C-terminal end of the motif. The permutations shared catalytic geometry but specified different sequence routings through the active site. We prioritised orderings that more frequently yielded low AF3 motif RMSD for RFD4-Proteina structure and sequence generation, using motif RMSD as the primary search reward.

##### Initial filtering and parent selection

We screened generated designs for motif RMSD and non-bonded protein–ligand clashes before AF3 prediction, then applied the fold, motif and confidence criteria in Supplementary Table 7. We evaluated passing candidates against the TI2 catalytic-distance criteria in Supplementary Table 8, then predicted them as AEI plus amine and applied the criteria in Supplementary Table 9.

Supplementary Table 7 | Initial design filters.

| Filter | Requirement | Stage applied |
|---|---|---|
| Design protein-motif RMSD (motif_rmsd) | ≤ 1.0 Å | Pre-fold |
| Refold protein-motif RMSD (motif_allatom_rmsd) | ≤ 1.0 Å | Post-fold |
| Refolded backbone RMSD versus design | ≤ 1.5 Å | Post-fold |
| AF3 overall pLDDT | ≥ 80 | Post-fold |

| Filter | Requirement | Stage applied |
|---|---|---|
| Protein–ligand clashes | Zero, with clashes defined as any non-bonded atoms within 1.5A | Pre- and post-fold |

Supplementary Table 8 | TI2 distance filters

| Filter | Requirement |
|---|---|
| Distance Cys SG to His ND1 or NE2 | ≤ 4.0 Å |
| Distance His ND1 or NE2 to Asp OD1 or OD2 | ≤ 3.2 Å |
| Distance Ligand O29 to Cys N | ≤ 3.0 Å |
| Distance Ligand O29 to Gln NE2 | ≤ 3.5 Å |

Supplementary Table 9 | AEI + amine distance filters

| Filter | Requirement |
|---|---|
| Distance ligand N43 to His | < 6.0 Å |
| Distance His to Asp | < 4.0 Å |
| Distance Cys SG to His | < 6.0 Å |
| Distance ligand O29 to Cys N | < 4.5 Å |
| Distance ligand O29 to Gln NE2 | < 6.0 Å |
| Distance acyl C28 to acceptor N43 | < 4.5 Å |
| ligand pLDDT | > 60 |
| protein pLDDT | > 80 |

We evaluated ten AF3 seeds per design in each of TI2 and AEI. We assigned the CIP configuration of acceptor C01 with RDKit from each predicted structure and the corresponding state's CCD bond graph, and calculated ligand confidence as the mean pLDDT over ligand atoms. We required parents to recover S configuration in at least one prediction independently in each state and to meet the confidence floors in Supplementary Table 9. We then visually inspected the structures and rejected pockets that were too shallow or too deeply buried. We selected retained parents directly for experiment or expanded them by either of the following refinement routes.

#### Partial-diffusion expansion and filtering

RFD4 partial diffusion diversified retained parents within a beam-search expansion using half of the 400-step sampling trajectory. The value 0.5 specified the fraction of trajectory steps used for partial diffusion. Rewards incorporated TI2 catalytic distances in the generated structure, the corresponding distances in AF3 TI2 predictions and AEI-plus-amine geometry. Retention required the TI2 design criteria together with the TI2 and AEI prediction criteria and pLDDT ≥80 for the designed enzyme chain in each predicted state. An early implementation reused TI2 confidence for the AEI gate; we subsequently filtered those designs using AEI confidence so that all retained designs met the same standard.

We evaluated final geometry on the top-ranked model from a fresh ten-seed AF3 prediction of each sequence in each state. We retained all four TI2 distance filters in Supplementary Table 8. We applied the six AEI distance criteria in Supplementary Table 9 together, tightening O29–Gln Nε2 to <4.5 Å and acyl C28–acceptor N43 to <4.0 Å while leaving the other four limits unchanged. We did not reapply generation-stage confidence, RMSD, clash or motif-integrity gates during this final geometric screen. A separate stereochemical filter required C01 to adopt S configuration in at least two of ten predictions independently in TI2 and AEI, using each state's CCD bond graph for assignment. Supplementary Tables 10 and 12 give the filtering funnel and backbone provenance.

Supplementary Table 10 | Design funnel for partial diffusion of lenacapavir ligases

| **Stage** | **Distinct sequences** | **Backbones** |
|---|---|---|
| Delivered from partial diffusion | 949 | 17 |
| AEI geometry passers | 235 | 16 |
| TI2 geometry passers | 130 | 13 |
| Dual passers (AEI and TI2) | 46 | 9 |
| ≥2 S seeds in both states | 30 | 8 |

#### Iterative LigandMPNN–AF3 expansion

A separate sequence-diversification branch used three rounds of LigandMPNN design on AF3-predicted TI2 structures from the retained parents. After the first two rounds, structures passing motif, confidence and contact filters supplied coordinates for the next redesign round (Supplementary Table 11). LigandMPNN used temperature 0.1, ligand and side-chain context, and side-chain packing. We fixed His514, Asp531, Gln574 and Cys580 identities while allowing the flanking Asp579 and Gly581 positions to vary. We excluded additional cysteines at designable positions while allowing histidine, aspartate and glutamine elsewhere. In rounds 2 and 3, we distributed sampling among passing descendants within each parent family to limit overrepresentation of families with many passing variants.

Cycling predictions used five randomised seeds and five diffusion samples per seed, with empty multiple-sequence alignments and no protein templates. We averaged motif RMSD, chain-A heavy-atom pLDDT and contact distances over predictions passing the state-specific stereochemical checks, and took pTM from the original highest-ranked output. We used the highest-ranked model to represent the sequence unless it failed the stereochemical checks and a passing alternative was available, in which case we used the passing model with highest chain-A pLDDT. Here chain A was the designed enzyme.

Supplementary Table 11 | TI2 filters during iterative LigandMPNN/AF3 expansion. We required all criteria within a column simultaneously.

| **Criterion** | **First round** | **Second and final rounds** |
|---|---|---|
| Mean motif RMSD | ≤ 1.25 Å | ≤ 1.0 Å |
| Mean chain-A pLDDT | ≥ 75 | ≥ 80 |
| pTM | > 0.75 | > 0.85 |
| Mean histidine contact distance, Nu_HIS | ≤ 4.0 Å | ≤ 4.0 Å |
| Mean histidine contact distance, LG_HIS | ≤ 4.0 Å | ≤ 4.0 Å |

| Criterion | First round | Second and final rounds |
|---|---|---|
| Mean backbone oxyanion-hole contact distance, Oxy_bb | ≤ 3.5 Å | ≤ 3.5 Å |
| Mean side-chain oxyanion-hole contact distance, Oxy_sc | Not applied | ≤ 4.0 Å |

For experimental selection, we screened variants passing the final TI2 criteria against the same six tightened AEI distance cutoffs used for partial-diffusion descendants. Contacts involving histidine and aspartate used the minimum over His Nδ1/Nε2 and Asp Oδ1/Oδ2, respectively. Fresh ten-seed predictions with one diffusion sample per seed then required S configuration at C01 in at least four predictions independently in TI2 and AEI. We selected the resulting variants as LL31–LL48 (Supplementary Table 12). A separate four-state assessment evaluated donor-bound enzyme, TI1, AEI and TI2, combining the final TI2 criteria with motif RMSD ≤1.5 Å and pTM >0.80 in each of the other states; it was distinct from selection of the tested MPNN variants.

#### Retrospective AF3 ensemble analysis

To assess product accommodation, we generated ten AF3 product-state predictions for each of the 67 designs, using ten seeds and one sample per seed (Supplementary Fig. 11). We represented the enzyme with an unmodified catalytic cysteine and supplied the amide product as a separate component. For each prediction, we measured d1 from the catalytic cysteine backbone nitrogen to product carbonyl O29 and d2 from Cys Sγ to product carbonyl C28. We took the minimum of each distance independently across the ten predictions, so a design's d1 and d2 minima could come from different models. We compared these per-design minima with the experimental activity classes and retrospectively applied d1 <3.5 Å and d2 <4.0 Å cutoffs. For the enrichment calculation, we included only the 55 designs with experimental readouts: the criteria retained all four ligase hits among 32 designs, compared with four hits among all 55 tested designs. We did not use these product-state criteria for prospective selection.

Supplementary Table 12 | Origins of all tested lenacapavir ligase designs

| Design ID | Origin | RFD4-Proteina parent design |
|---|---|---|
| LL1 | Partial diffusion | LL49 |
| LL2 | Partial diffusion | LL49 |
| LL3 | Partial diffusion | LL51 |
| LL4 | Partial diffusion | LL54 |
| LL5 | Partial diffusion | LL54 |
| LL6 | Partial diffusion | LL54 |
| LL7 | Partial diffusion | LL54 |
| LL8 | Partial diffusion | LL54 |
| LL9 | Partial diffusion | LL54 |
| LL10 | Partial diffusion | LL54 |
| LL11 | Partial diffusion | LL54 |

| Design ID | Origin | RFD4-Proteina parent design |
|---|---|---|
| LL12 | Partial diffusion | LL54 |
| LL13 | Partial diffusion | LL54 |
| LL14 | Partial diffusion | LL54 |
| LL15 | Partial diffusion | LL54 |
| LL16 | Partial diffusion | LL54 |
| LL17 | Partial diffusion | LL54 |
| LL18 | Partial diffusion | LL54 |
| LL19 | Partial diffusion | LL54 |
| LL20 | Partial diffusion | LL54 |
| LL21 | Partial diffusion | LL54 |
| LL22 | Partial diffusion | LL54 |
| LL23 | Partial diffusion | LL55 |
| LL24 | Partial diffusion | LL55 |
| LL25 | Partial diffusion | LL58 |
| LL26 | Partial diffusion | LL61 |
| LL27 | Partial diffusion | LL61 |
| LL28 | Partial diffusion | LL65 |
| LL29 | Partial diffusion | LL67 |
| LL30 | Partial diffusion | LL67 |
| LL31 | MPNN / AF3 cycling | LL52 |
| LL32 | MPNN / AF3 cycling | LL63 |
| LL33 | MPNN / AF3 cycling | LL63 |
| LL34 | MPNN / AF3 cycling | LL63 |
| LL35 | MPNN / AF3 cycling | LL63 |
| LL36 | MPNN / AF3 cycling | LL63 |
| LL37 | MPNN / AF3 cycling | LL63 |
| LL38 | MPNN / AF3 cycling | LL63 |
| LL39 | MPNN / AF3 cycling | LL63 |
| LL40 | MPNN / AF3 cycling | LL63 |
| LL41 | MPNN / AF3 cycling | LL63 |
| LL42 | MPNN / AF3 cycling | LL63 |
| LL43 | MPNN / AF3 cycling | LL66 |
| LL44 | MPNN / AF3 cycling | LL66 |
| LL45 | MPNN / AF3 cycling | LL66 |

| Design ID | Origin | RFD4-Proteina parent design |
| --- | --- | --- |
| LL46 | MPNN / AF3 cycling | LL66 |
| LL47 | MPNN / AF3 cycling | LL66 |
| LL48 | MPNN / AF3 cycling | LL66 |
| LL49 | Direct RFD4-Proteina | — (de novo RFD4-Proteina output) |
| LL50 | Direct RFD4-Proteina | — (de novo RFD4-Proteina output) |
| LL51 | Direct RFD4-Proteina | — (de novo RFD4-Proteina output) |
| LL52 | Direct RFD4-Proteina | — (de novo RFD4-Proteina output) |
| LL53 | Direct RFD4-Proteina | — (de novo RFD4-Proteina output) |
| LL54 | Direct RFD4-Proteina | — (de novo RFD4-Proteina output) |
| LL55 | Direct RFD4-Proteina | — (de novo RFD4-Proteina output) |
| LL56 | Direct RFD4-Proteina | — (de novo RFD4-Proteina output) |
| LL57 | Direct RFD4-Proteina | — (de novo RFD4-Proteina output) |
| LL58 | Direct RFD4-Proteina | — (de novo RFD4-Proteina output) |
| LL59 | Direct RFD4-Proteina | — (de novo RFD4-Proteina output) |
| LL60 | Direct RFD4-Proteina | — (de novo RFD4-Proteina output) |
| LL61 | Direct RFD4-Proteina | — (de novo RFD4-Proteina output) |
| LL62 | Direct RFD4-Proteina | — (de novo RFD4-Proteina output) |
| LL63 | Direct RFD4-Proteina | — (de novo RFD4-Proteina output) |
| LL64 | Direct RFD4-Proteina | — (de novo RFD4-Proteina output) |
| LL65 | Direct RFD4-Proteina | — (de novo RFD4-Proteina output) |
| LL66 | Direct RFD4-Proteina | — (de novo RFD4-Proteina output) |
| LL67 | Direct RFD4-Proteina | — (de novo RFD4-Proteina output) |

### 2.2 Experimental materials and methods

#### 2.2.1 Genetic constructs

Construction of pET26_LL52_C177A and pET26_LL58_C144A

Genes encoding LL52_C177A and LL58_C144A were constructed using overlap extension PCR with end primers designed for Golden Gate assembly. Genes were cloned into a pET26-ccdB vector using BsaI-HFv2, employing the golden gate assembly method leading to fusion to a C-terminal hexa-histidine tag coding sequence.

Supplementary Table 13 | Primers for lenacapavir ligases

| | |
|---|---|
| D52_ C177A _F | GCATTAAAGGCATTGGCAACGCGATTGTGACCATT |
| D52_ C177A _R | GTTGCCAATGCCTTTAATGC |
| D52_GGFlank_F | atactacggtctcaaggaGAAGAAGAAATCGAGAAAGCG |
| D52_GGFlank_R | gcattacggtctcgggaaccCAGTTCTTCCAGAATTTTCAGC |
| D58_ C144A _F | GTCATCCGGCGAACGCGGAAGTGACTACC |
| D58_ C144A _R | GTTCGCCGGATGAC |
| D58_GGFlank_F | atactacggtctcaaggaCCGCCGGCTGAA |
| D58_GGFlank_R | gcattacggtctcgggaaccCGCCGGTTCAAACAC |

#### 2.2.2 Molecular cloning

DNA sequences encoding the designed ligases were ordered as synthetic gene fragments from Integrated DNA Technologies (IDT). Gene fragments were cloned into the pDT1 (Addgene plasmid #262810) expression vector using Golden Gate assembly with the NEBridge Golden Gate Assembly Kit (BsaI-HFv2; New England Biolabs, Cat. No. E1601). Assembly reactions were subjected to 30 cycles of 37 °C for 1 min and 16 °C for 1 min, followed by incubation at 60 °C for 5 min. The pDT1 vector contains a T7 promoter, lac operator, kanamycin resistance cassette, and a *ccdB* negative-selection cassette.

#### 2.2.3 Protein expression and purification

##### High-throughput protein expression and purification for screening

Golden Gate assembly products were transformed into chemically competent *E. coli* BL21(DE3) cells (New England Biolabs, Cat. No. C2527) according to the manufacturer's protocol. Following transformation, cells were grown overnight in LB medium containing kanamycin (50 µg/mL) without isolating individual colonies, maintaining a polyclonal population for each design. For each design, eight 1 mL cultures for N-terminal ligases (NL) or two 1 mL cultures for lysine ligases (KL) of Terrific Broth II (TBII; MP Biomedicals, Cat. No. 113046022) autoinduction medium were inoculated with 10 µL of the corresponding overnight LB culture per 1 mL of medium. The autoinduction medium was supplemented with 1× M salts (25 mM $Na_2HPO_4$, 25 mM $KH_2PO_4$, 50 mM $NH_4Cl$, and 5 mM $Na_2SO_4$), 1× 5052 (0.5% [w/v] glycerol, 0.05% [w/v] glucose, and 0.2% [w/v] α-lactose), and kanamycin (50 µg/mL). Cultures were incubated for 6 h at 37 °C, followed by 22 h at 22 °C. Following expression, cells were harvested by centrifugation at 4,000 × g for 30 min. Cell pellets were resuspended in lysis buffer (see individual ligase sections for details) containing 100 mM HEPES (pH 7.4 for NL and pH 8 for KCL purifications), 50 mM NaCl, DNase (10 µg/mL), lysozyme (1 mg/mL), 1 mM tris(2-carboxyethyl)phosphine (TCEP;

Sigma-Aldrich, Cat. No. 646547), and 1× BugBuster Protein Extraction Reagent (MilliporeSigma, Cat. No. 70921), and incubated at 37 °C for 30 min with shaking. Lysed cells were clarified by centrifugation at 4,000 × g for 40 min. Clarified lysate was then applied to 100 µL of Strep-Tactin XT 4Flow high-capacity resin (IBA Lifesciences, Cat. No. 2-5030-025), pre-equilibrated with 100 mM HEPES (pH 7.4 for NL and pH 8 for KCL purifications), 50 mM NaCl, and 1 mM TCEP. After washing with five column volumes (CVs) of equilibration buffer, bound proteins were eluted with 50 mM biotin in 100 mM HEPES (pH 7.4 for NL and pH 8 for KCL purifications), 50 mM NaCl, and 1 mM TCEP. Eluted proteins were further purified by size-exclusion chromatography (SEC) using a Superdex™ 75 Increase column 5/150 GL (Cytiva; #29148722) on an Agilent 1290 Infinity III Bio LC system . Fractions containing purified enzymes were pooled and flash-frozen in liquid nitrogen for storage at −80 °C until use.

#### Expression and purification of N-term ligases

Refer to the general cloning procedure for cloning details. Monoclonal sequence verified glycerol stocks of NL6 and NL6 KOs BL21 (DE3) cells were cultured in 50 mL volumes of Terrific Broth II (TBII; MP Biomedicals, Cat. No. 113046022) containing kanamycin (50 µg/mL) for 6 hours or until reaching $OD_{600}$ of 0.6 to 1.0 at 37 ºC. Expression was then induced using IPTG (1mM). Expression temperature was reduced to 20 ºC and incubated for an additional 12-16 hours before harvesting. Cells were harvested after expression by centrifugation at 4000 g for 30 minutes. Cell pellets were then resuspended in 100 mM Hepes pH 7.4, 150 mM NaCl, 1 mM EDTA, 1 mM PMSF, 1 mg/mL lysozyme buffer. Lysis was performed by sonication (Qsonica 500 with 4 tip ¼" horn; #Q500A-110, #4674) at 70 % amplitude, 5 seconds on and 5 seconds off, for a total active time of 10 minutes. Lysed samples were then clarified by centrifugation at 14000 g for 35 minutes. Clarified lysate was dispensed over pre-equilibrated Cytiva™ Protein Select™ resin (Cytiva; 17542101) with 100 mM HEPES pH 7.4, 50 mM NaCl and 2 mM TCEP. After washing with multiple column volumes, samples were left to incubate on resin overnight at room temperature for intein based N-terminal cleavage. Eluate was then collected with resin wash with additional 100 mM Hepes pH 7.4 and 50 mM NaCl buffer. Eluate was concentrated using Amicon® Ultra Centrifugal Filter, 3 kDa MWCO (Millipore; UFC8003). Concentrated samples then underwent size exclusion chromatography on a Cytiva AKTA PURE FPLC system with a Superdex 200 Increase SEC column, 10/300 GL (Cytiva; 28990944) equilibrated with a 100 mM Hepes pH 7.4, 50 mM NaCl and 2 mM TCEP buffer. Finally, monomeric fractions were collected, pooled and spin concentrated using aforementioned centrifugal filters. Aliquots were then flash frozen with liquid nitrogen and stored at -80 ºC before use.

#### Expression and purification of fused target substrate

Cloning, expression, harvesting of cells was performed as previously described. Cells were resuspended in 100 mM Hepes pH 7.4, 150 mM NaCl, 1 mM EDTA, 1 mM PMSF, 1× cOmplete™, EDTA-free Protease Inhibitor Cocktail (Roche; 04574834001) and 1 mg/mL lysozyme. Lysis was performed by sonication (Qsonica 500 with 4 tip ¼" horn; #Q500A-110, #4674) at 70 % amplitude, 5 seconds on and 5 seconds off, for a total active time of 10 minutes. Lysed samples were then clarified by centrifugation at 14000 g for 35 minutes. Clarified lysate was then dispensed over pre-equilibrated Streptactin XT 4Flow high-capacity resin (IBA Lifesciences, Cat. No. 2-5030-025) with 100 mM Hepes pH 7.4, 50 mM NaCl buffer. Multiple CV washes were performed with alternating salt concentrations (500 mM and 50 mM NaCl) in the aforementioned buffer. Samples were eluted with the equilibration buffer supplemented with 50 mM Biotin. Elutate was then dispensed over a pre-equilibrated column containing Cytiva™ Protein Select™ resin (Cytiva; 17542101) with 100 mM Hepes pH 7.4 and 50 mM NaCl buffer. After washing with multiple column volumes, samples were left to incubate on resin overnight at room temperature for intein based N-terminal cleavage. Eluate was then collected with additional 2 CV resin washes with equilibration buffer. Eluate was concentrated using Amicon® Ultra Centrifugal Filter, 3 kDa MWCO (Millipore; UFC8003). Aliquots were then flash frozen with liquid nitrogen, and stored at -80 ºC before use.

#### Expression and purification of lysine ligases

Designs were obtained as full-length synthetic constructs from Twist Biosciencies in pET29b vectors carrying a C-terminal strep tag and fused to GB1 (Sequences section). Plasmids were transformed into chemically competent *E. coli* BL21 (DE3). Single colonies were used to inoculate 1 mL LB starter cultures containing kanamycin (50 µg/mL), which were grown for 4–6 h. Then 200 µL of starter culture was used to inoculate 50 mL autoinduction medium (50 g/L Terrific Broth II (MP Biomedicals), 2 mM $MgSO_4$, 1× 5052 and 50 µg/mL kanamycin). Cultures were grown for 6–8 h at 37 °C followed by overnight expression at 18 °C.

Cells were harvested by centrifugation (4,000 g, 10 min) and pellets were used directly or stored at −80 °C. Pellets were resuspended in 35 mL lysis buffer (100 mM HEPES, 50 mM NaCl, pH 8.0, 1 mM TCEP, supplemented with PMSF, lysozyme and DNase I from 100×, 1,000× and 1,000× stocks, respectively) by shaking for 10 min, and lysed by sonication at 80% amplitude with 10 s on/10 s off cycles for 10 min total on-time. Lysates were clarified by centrifugation (14,000 g, 30 min). All subsequent steps were performed at 4 °C. A Strep-Tactin column was washed with 20 mL water and equilibrated with 20 mL lysis buffer, the clarified lysate was loaded and left on the resin for 20 min, and the resin was washed three times with 15 mL of 100 mM HEPES, 50 mM NaCl, pH 8.0, 1 mM TCEP. Protein was pre-eluted with 800 µL of the same buffer supplemented with 50 mM biotin, and then eluted with a further 1.5 mL of this elution buffer left on the resin for 10 min before flowing through. Eluates were cleared by centrifugation (14,000 g, 5 min) and purified by size exclusion chromatography in 100 mM HEPES, 50 mM NaCl, pH 8.0, 1 mM TCEP. Fractions containing purified enzymes were then pooled and flash frozen in liquid nitrogen for storage at -80 ºC until used.

#### Expression and purification of lenacapavir ligases

Designs were expressed in chemically competent E. coli BL21 (DE3) cells. 5 mL LB medium containing kanamycin (50 µg/mL) was inoculated and cultured for 18 h. Starter cultures (500 µl) were then used to inoculate baffled flasks containing auto-inducible TB II media (50 mL) and kanamycin (50 µg/mL). Cultures were grown (37 °C, 180 r.p.m.) for 6 h, then the incubation temperature was reduced to 25 °C for a further 16 h. Cells were collected by centrifugation (3,220 x g for 10 min). Cells were resuspended in lysis buffer (50 mM HEPES, 300 mM NaCl, pH 7.4, 10 mM imidazole, 10 µg mL−1 DNase I, 1 mM phenylmethylsulfonyl fluoride) and lysed by sonication. Cell lysates were clarified by centrifugation (27,216 x g for 15 min), and the soluble fraction was subjected to affinity chromatography using Ni-NTA agarose (Qiagen). Unbound proteins were removed with a lysis buffer supplemented with an additional 10 mM imidazole. Purified protein was eluted using 50 mM HEPES, 300 mM NaCl, pH 7.4, containing 300 mM imidazole. Proteins were desalted using 10DG desalting columns (Bio-Rad) with 100 mM KPi buffer, pH 6.

#### X-ray crystallography protein expression and purification

Cloning procedure followed as previously described. Expressions were conducted from sequence verified (Plasmidsaurus) monoclonal glycerol stock culture of NL6 containing active site cysteine to alanine mutation. 5 mL Luria Broth (LB) overnight starter cultures containing kanamycin (50 µg/mL) were used to inoculate 2 L of TBII (TBII; MP Biomedicals, Cat. No. 113046022) in baffled 1L Erlenmeyer flasks.

Expression was induced using IPTG (1 mM) after 4-6 hours of incubation at 37 ºC or until reaching $OD_{600}$ of 0.6-1.0. Incubation temperature was then reduced to 20 ºC and cultures continued to incubate for an additional 12-16 hours before harvest. Cells were harvested by centrifugation at 4000xg for 30 minutes. The cell pellet was then resuspended in a lysis buffer containing 100 mM Hepes pH 7.4, 50 mM NaCl, 1 mM EDTA, 2 mM TCEP, 1 mg/mL lysozyme, and 1 mM PMSF. Homogenised pellet was then lysed by sonication using previously described method. Insoluble material was clarified by centrifugation at 14000 g for 35 minutes. Clarified lysate was dispensed over pre-equilibrated Cytiva™ Protein Select™ resin (Cytiva; 17542101) with 100 mM HEPES pH 7.4, 50 mM NaCl and 2 mM TCEP. Flow through was reloaded over the column once. The wash was then performed with multiple CVs using an equilibration buffer with alternating salt concentrations (50 mM and 500 mM NaCl). Flow was stopped and resin was left to incubate at room temperature for 16 hours for N-terminal cleavage. Eluate was then collected with multiple CVs of equilibration buffer. Collected samples were then concentrated using Amicon® Ultra Centrifugal Filter, 3 kDa MWCO (Millipore; UFC8003). Concentrated samples underwent size exclusion chromatography using AKTA PURE FPLC system equipped with a Superdex 200 Increase SEC column, 10/300 GL (Cytiva; 28990944) equilibrated with a 20 mM Hepes pH 7.4, 20 mM NaCl and 200 µM TCEP buffer. Monomeric fractions were then pooled and concentrated using Amicon® Ultra Centrifugal Filter, 3 kDa MWCO (Millipore; UFC5003) to a final concentration of 49.05 mg/mL for crystallisation condition screening.

#### 2.2.4 Hydrolysis and aminolysis activity screen

##### 2.2.4.1 General procedure for N-term ligase screening

All reactions containing NL designs except B12, C9, H3, and H4, which contained 3.3, 0.9, 4.4, and 2.4 µM of enzyme, were normalised to 5 µM of enzyme per reaction. Designs C8 and H11 failed to express and purify and were excluded from the screening experiment. 94 reactions were prepared containing 100 µM of SPPS prepared FSLTK substrate (Genescript) and 200 µM of TAMRA-Gly-TFE (TAMRA-donor) in 100 mM Hepes pH 7.4, 50 mM NaCl, 1 mM TCEP, and 5% DMSO. Reactions were then allowed to proceed for 27 hours at 22 ºC before quenching with acetonitrile to a final concentration of 75%. Quenched samples were then centrifuged at 2000 g for 5 minutes. Sample species were then quantified using reverse phase liquid chromatography on an Agilent 1260 Infinity II HPLC system equipped with a Higgins PROTO 300 $C_{18}$, RS-1546-W185, 150 × 4.6 mm, 5 µm, column using a stepwise gradient method (Supplementary Table 14). Mobile phase A consisted of 99.9% water and 0.1% trifluoroacetic acid (TFA), and mobile phase B consisted of 99.9% acetonitrile and 0.1% TFA (all percentages v/v). Chromatograms were collected across a series of absorbances including 205 nm and 555 nm absorbance.

Observed peak areas for 555 nm chromatograms were used in the quantification of substrate to product conversion. Substrate to product conversion was estimated using the following formulation: $p/(h + p + d) \times 100 = \%\, conversion$ (Eq. 1); where $p$ is the observed product peak area; $d$, is the observed donor peak area; $h$, is the observed hydrolysis peak area. The hydrolysis conversion was similarly calculated using hydrolysis peak area over the same summed species peak areas. Two controls were included to measure the background rate of donor hydrolysis and product formation without enzyme.

Supplementary Table 14 | HPLC Gradient for Aminolysis / Hydrolysis screening

| Time (minutes) | A (%) | B (%) | Flow (mL/min) |
|---|---|---|---|

| | | | |
|---|---|---|---|
| 0.00 | 95 | 5 | 1.0 |
| 2.50 | 95 | 5 | 1.0 |
| 3.86 | 30 | 70 | 1.0 |
| 7.19 | 15 | 75 | 1.0 |
| 8.91 | 10 | 90 | 1.0 |
| 9.51 | 5 | 95 | 1.0 |
| 11.51 | 5 | 95 | 1.0 |
| 12.71 | 95 | 5 | 1.0 |
| 15.0 | 95 | 5 | 1.0 |

##### General procedure for N-term ligase conversion, knockout and regioselectivity experiments

Monoclonal and sequence verified expressed and purified enzymes were used in the following experiments. Reactions were performed in triplicate with 10 μM enzyme, 50 μM FSLTK/FSLTK-Spy/spytag003 (Bio-Rad; BLP086), 100 μM TAMRA-donor, in 5% DMSO, 100 mM Hepes pH 7.4 and 50 mM NaCl buffer at 22 ºC for 16 hours. Aliquots for each reaction were taken across the time series (0 to 16 hours) and quenched with acetonitrile to 75% followed by centrifugation at 2000 g for 5 minutes. Samples were then measured using reverse phase HPLC method as previously described.

Observed peak areas of interest at 205 nm were used to estimate the conversion of peptide substrate to ligated peptide product using the following equation: $p\,/\,(p + s) \times 100 = \%\,conversion$ (Eq. 2); where, $p$, is the observed product peak area and, $s$, is the observed substrate peak area. Controls were included to measure the rate of background donor hydrolysis and product formation without enzyme.

Observed peak areas of interest at 555 nm were used to estimate the hydrolysis of TAMRA-donor over the course of the reaction using the equation following Eq. 1. Peak areas were quantified as previously described.

##### 2.2.4.2 General procedure for lysine ligase screening

Lysine ligases as well as the corresponding peptide substrates were expressed and purified using the protocol outlined in 2.2.3 and screened in 50 µl reactions with 10 µl crude enzyme, 35 µl crude substrate peptide fused to the C-terminus of MBP, and 800 uM acyl donor in 10% (v/v) DMSO and 100 mM HEPES, 50 mM NaCl, 1 mM TCEP, pH 8.0. Reactions were run overnight at room temperature and quenched with 0.1% formic acid for LC/MS analysis and with 90% (v/v) acetonitrile for HPLC analysis. Product was quantified through apparent conversion (Eq 2) from mass spectrometry and the measured concentration of the substrate from nanodrop, and normalized to the measured enzyme concentration, which was also measured via nanodrop. Lysine ligases were screened using the same method as previously described for N-terminal ligases.

##### 2.2.4.3 General procedure for analytical scale biotransformations for lenacapavir-API amide ligation

Analytical scale biotransformations were performed using ester 1 (1 mM), enantiopure amine 2 (0.5 mM), and the relevant purified enzyme (50 µM) in KPi buffer (100 mM, pH 6) with 20% DMSO, incubated at 30°C (16 h, 850 r.p.m.). The reaction was quenched with 1 volume acetonitrile containing 0.1% HCl and shaken at 30°C, 850 r.p.m. for 20 min. Precipitated protein was removed by centrifugation (3,220 x g for 15 minutes) and the supernatant used for UPLC analysis.

#### 2.2.5 LC/MS assays and analysis

##### 2.2.5.1 N-term

###### Fusions

Reactions were performed with 4 µM enzyme, 16 µM acceptor, and 500 µM Tamra-donor in 10% DMSO, 100 mM Hepes pH 8 and 50 mM NaCl. Fusion experiments were performed using Fusion-GB1 and FusionKO-GB1 (knockout) substrates. Knockout contains the mutation of Lysine (K) to Arginine (R) on the KL37 target substrate. Reactions were incubated at 22 ºC for 12 hours and then quenched with cold formic acid to a final concentration of 0.1% and stored at 4 ºC before sample acquisition.

Specimens were analysed using LC/MS on an Agilent 1290 Infinity II coupled to an Agilent 6230C TOF mass spectrometer.

##### 2.2.5.2 Lysine ligase

Reactions were set up with 4 µM enyzme, 16 µM acceptor and 800 µM donor and quenched in 0.1% formic acid. Specimens were analysed using the aforementioned LC/MS system.

#### 2.2.6 HPLC assays

Hydrolysis was measured in a separate set of reactions. Each was 160 µL and contained 4 µM KL37-GB1 and 16 µM KL37AP-MBP, or donor alone, with 800 µM alkyne donor and 10% (v/v) DMSO in 100 mM HEPES, 50 mM NaCl, pH 7.4 at 37°C. 10 µL of reaction was quenched into 90 µL of acetonitrile, vortexed and centrifuged, and the supernatant was analysed by  Following the same methodology outlined in N-terminal ligase hydrolysis screening.

#### 2.2.7 Gel assays

###### Lysine ligase in gel fluorescence and gel shift assay

10 µL was quenched into 3.3 µL of 4× SDS loading buffer and boiled at 95 °C for 5 min for SDS-PAGE analysis. Boiled SDS aliquots (13.3 µL) were labelled with m-$PEG_{48}$-azide (TAMRA-azide for in gel fluorescence analysis) by copper-catalysed azide–alkyne cycloaddition to give a mass shift resolvable by SDS-PAGE. To each sample was added 4 µL of click master mix, giving final concentrations of 3.61 mM TAMRA-azide, 2.5 mM THPTA, 1 mM $CuSO_4$ and 5 mM sodium ascorbate in 17.3 µL. The master mix was prepared by combining THPTA with $CuSO_4$, then adding m-$PEG_{48}$-azide and buffer, with sodium ascorbate added last immediately before use. Reactions proceeded for 30–60 min at room temperature in the dark. Gels were either stained using coomassie or imaged in a gel scanner.

#### 2.2.8 X-ray crystallography

All crystallisation experiments were conducted using the sitting drop vapour diffusion method. Crystallisation trials were set up in 200 nL drops using the 96-well plate format at 20 ˚C. Crystallisation plates were set up using a Mosquito LCP from SPT Labtech, then imaged using UVEX microscopes and UVEX PS-256 from JAN Scientific. Diffraction quality crystals for GLS_2049 (NL6, cystine to alanine mutation) formed in 0.2M ammonium sulfate, 0.01M cadmium chloride hemi(pentahydrate), 0.1M PIPES pH 7, 15% v/v PEG 1,000, and 10% v/v ethylene glycol. The apo form of KL37 crystallised with 0.05 M Magnesium chloride, 0.1 M HEPES, pH 7.5, 30 %(v/v) PEG 550 MME and the holo form with 0.2 M Sodium thiocyanate, 20% w/v PEG 3350.

Diffraction data was collected at the 17-ID-1 (AMX) beamline of the National Synchrotron Light Source II. X-ray intensities and data reduction were evaluated and integrated using XDS[9] and merged/scaled using Pointless/Aimless in the CCP4 program suite[10]. Structure determination and refinement starting phases were obtained by molecular replacement using Phaser[11] using the designed model for the structures. Following molecular replacement, the models were improved using phenix.autobuild[12]; efforts were made to reduce model bias by setting rebuild-in-place to false, and using simulated annealing and prime-and-switch phasing. Structures were refined in Phenix[12]. Model building was performed using COOT[13]. The final model was evaluated using MolProbity (6). Data collection and refinement statistics are recorded in Supplementary Table 15.

Supplementary Table 15 | GLS_2049 crystal structure

| | GLS_2049 |
|---|---|
| Data collection | |
| Space group | *P 61* |
| Cell dimensions | |
| *a*, *b*, *c* (Å) | 94.548, 94.548, 82.754 |
| α, β, γ (°) | 90, 90, 120 |
| Resolution (Å) | 41.05 - 2.18 (2.25 - 2.18) |
| $R_{sym}$ or $R_{merge}$ | 0.152 (1.751) |
| $I / \sigma I$ | 12.5 (2.2) |
| Completeness (%) | 100.0 (100.0) |
| Redundancy | 21.4 (21.7) |
| Refinement | |
| Resolution (Å) | 41.05 - 2.18 (2.28 - 2.18) |
| No. reflections | 22,008 (2,732) |
| $R_{work}$ / $R_{free}$ | 0.2184 (0.2969) / 0.2553 (0.3481) |

| | GLS_2049 |
|---|---|
| Data collection | |
| Space group | *P 61* |
| Cell dimensions | |
| *a*, *b*, *c* (Å) | 94.548, 94.548, 82.754 |
| α, β, γ (°) | 90, 90, 120 |
| No. atoms | |
| Protein | 2,684 |
| Ligand/ion | 59 |
| Water | 57 |
| *B*-factors | |
| Protein | 69 |
| Ligand/ion | 86 |
| Water | 63 |
| R.m.s. deviations | |
| Bond lengths (Å) | 0.003 |
| Bond angles (°) | 0.51 |

*Single Crystal used for each data collection. *Values in parentheses are for highest-resolution shell.

##### 2.2.9 Analysis of X-ray crystal structure and AF3 predictions

Experimentally determined crystal structures were compared with the corresponding highest-confidence AlphaFold3 (AF3) predictions using PyMOL (Schrödinger, LLC). Predicted structures were superimposed onto the crystal structures by alignment of corresponding protein Cα atoms. Global Cα root-mean-square deviations (RMSDs) were calculated without iterative outlier rejection. To assess agreement of the catalytic architecture, all atom catalytic motif RMSDs were calculated following global Cα alignment without additional superposition. Active-site geometries and interactions were analysed by visual inspection and measurement of interatomic distances in PyMOL.

#### 3 Sequences

##### 3.1 General proteins

FSLTK-Helix-GB1
MSGSGHHHHHHGGSGGSMVKIVSRKSLGVQNVYDIGVEKDHNFLLANGLIASNFSLTKGGSGGSGGSGGS
SIAEAFREAKEYLLAAAQQGGSGGSGGSGGSTYKLILNGKTLKGETTTEAVDAATAEKVFKQYANDNGVD
GEWTYDDATKTFTVTE

FSLTK-HelixKO-GB1

MSGSGHHHHHHGGSGGSMVKIVSRKSLGVQNVYDIGVEKDHNFLLANGLIASNFSLTKGGSGGSGGSGGS
SIAEAFREAREYLLAAAQQGGSGGSGGSGGSTYKLILNGKTLKGETTTEAVDAATAEKVFKQYANDNGVD
GEWTYDDATKTFTVTEGGSGGSWSHPQFEK

FSLTK-Spy
FSLTKGGSGGSGGSGGRGVPHIVMVDAYKRYK

SpyTag
RGVPHIVMVDAYKRYK

KL37AP-MBP
MSGHHHHHHGSGSGSMVKIVSRKSLGVQNVYDIGVEKDHNFLLANGLIASNSIAEAFREAKYLLAAAQQG
SGSGSKIEEGKLVIWINGDKGYNGLAEVGKKFEKDTGIKVTVEHPDKLEEKFPQVAATGDGPDIIFWAHDRF
GGYAQSGLLAEITPDKAFQDKLYPFTWDAVRYNGKLIAYPIAVEALSLIYNKDLLPNPPKTWEEIPALDKELK
AKGKSALMFNLQEPYFTWPLIAADGGYAFKYENGKYDIKDVGVDNAGAKAGLTFLVDLIKNKHMNADTD
YSIAEAAFNKGETAMTINGPWAWSNIDTSKVNYGVTVLPTFKGQPSKPFVGVLSAGINAASPNKELAKEFLE
NYLLTDEGLEAVNKDKPLGAVALKSYEEELAKDPRIAATMENAQKGEIMPNIPQMSAFWYAVRTAVINAAS
GRQTVDEALKDAQTN

KL37AP-K10R-MBP (KO)
HHHHHHGSGSGSMVKIVSRKSLGVQNVYDIGVEKDHNFLLANGLIASNSIAEAFREAREYLLAAAQQGSGS
GSKIEEGKLVIWINGDKGYNGLAEVGKKFEKDTGIKVTVEHPDKLEEKFPQVAATGDGPDIIFWAHDRFGG
YAQSGLLAEITPDKAFQDKLYPFTWDAVRYNGKLIAYPIAVEALSLIYNKDLLPNPPKTWEEIPALDKELKAK
GKSALMFNLQEPYFTWPLIAADGGYAFKYENGKYDIKDVGVDNAGAKAGLTFLVDLIKNKHMNADTDYSI
AEAAFNKGETAMTINGPWAWSNIDTSKVNYGVTVLPTFKGQPSKPFVGVLSAGINAASPNKELAKEFLENY
LLTDEGLEAVNKDKPLGAVALKSYEEELAKDPRIAATMENAQKGEIMPNIPQMSAFWYAVRTAVINAASGRQ
TVDEALKDAQTN

3.2 N-terminal ligases

NL1
MSGMLRLEIVDKEEALENFKRIIFEYLPKALKIFDEMLEKVYDQMNEEEKKKFEELKERHKEALE
LIDKIEEILDIEYVEHPDYDEISIKVKIDGKEERLVLFRLYHDPALADKSKETGGCLLIGLSQLDVTI
KVDNEKMLEFLKRMAEILGIPVEIIESAWSHPQFEK

NL2
MSGAAELLQKALAWLILLELKKRGVTLEELIAALKKADEIIAKEDPELAEKLRKKHEKTIAYLKA
HPELETMPLEEMEEKIEKLLDEAIEALKKIRTEEELREALDKAFRQGVCLAIAGYLDEVLAAWDR
LVERLVEALPEWRELIEEIWREKKEELKKLASAWSHPQFEK

NL3

MSGAAEKLREALSWLIFRELRKRGVTLEELRAALEAFDAEVAATDPELAARLRERHDATLELLER
NPELLEMPLEELDRLIEEKLDEAIEELKKIESDEELEKALREALKEGWCLARRGYLDEVLAAYEEL
RERLIEALPEKRELIEKIFDEAIARLRELASAWSHPQFEK

NL4
MSGMPPLTLLLVGRGITEEELELAREAAREVGLDDLPVYRIGWDEEKGEGYCFKLAAEKKAPRIP
TDKTLEELAEEHKKNAKEALKIEDEEKREEVLIEDVGRFIGALDRVLLLARDEEILETFKELLERA
GVEFLAFLHPDPSRPWTLEELKAWLREAQERFASAWSHPQFEK

NL5
MSGMELLTLILVGEGITEEEIELAREAARAVGMEGLPVYRFDWKPEEGKGYCFELAAEKDAPVIP
TDKTLEELAEEHKKNSEEAKKIKDKEEREKILIEDVGRFIGALKRVLLLTRTERERETFEKLLRRAG
VEYLIFTHPDPSRPWTKEELERWLREAQERFASAWSHPQFEK

NL6
MSGKIEIILIGFREEDLEKVKEELEKILKEKGVEDVTIYRSLIDPEKGEGYCLLLALERYDEVPETD
KTIEEAVEEHKENVKEIRKIPDPAERDKALAEDAIKVLATWKGKYVIIVYRPEEEEKARWLKEKLE
RLGMKVFLLNIEEPNWKEKLEEIADEIIKEHSAWSHPQFEK

NL7
MSGMLEIILIGFDEEIINEVKEELEKILKEMGVEDVTIYRMLYRPEEGEGYCLKLALERYDEVPPSD
KTIEEAVAEHKENVKEILKIPDPEKRDAALAEDAIKVLATFKGKPVIVVYTPENKEEAEELAEKLR
KLGMKVFLLDVTEENWKEKLEEIAKEIIEMYSAWSHPQFEK

NL8
MSGMLEIFLIGFDEEDLEEVRKELEEILKKRGVKDVTIYRSLYRPEEGEGYCLKLALEMYDEVPET
DKTIEEAVEEHKENVKEIEKIEDPEERDKALAEDAIKVLATLKGKPVIVVYRPEDRERAEELAEKL
RKLGMDVYLLNVEEEGWRERLEEIVDEILAKQSAWSHPQFEK

NL9
MSGMIEIILIGRREEDLEEVEKELKEILKEKGIENVTIYKALIQPEKGEGWCLLLALERYDSVPETD
MTIEEAIKKHEENVKEIRKIEDPEERERALAEDAIKVLATLKGKFVIIVYRPEDKEWAEELAEKLR
KLGMKVFLLNIEDPNWKEKLEEIVDKIIEEASAWSHPQFEK

NL10
MSGMLEVVLIGKDREALEKVKEELEKLLKERGVDNVTIFIAHLDPEKGEGWCLFLALERYDEVP
DTDKTLEEVNKEHKERVKEIRKIPDPAKREEALAEDAVKLLSALKGKIVIVVYRPEDEEEARELAE
KLRKIGADVHLLNVEAPDYEERLEEIVDEIIARLSAWSHPQFEK

NL11

MSGMKEVVLIGRDEKALREVKERLERILKERGVDNVTIFIALLDPEKGEGWCLFLALERYEEVPE
TDKTLEEVNAEHRKNVEEIRKIPDPEERERALAEDALKLLSALKGKIVIVVYKPEEREEAEELAEK
LRRIGAEVYLLDVEDPDYEERLEEIADEIIARLSAWSHPQFEK

NL12
MSGMKEVVLIGWDREALEEVKKRLEEILKERGVDNVTIYIAHLDPEKGEGWCLFLALEKYEEVP
DTDKTLEEVNKEHRENVKEIRKIPDPKKREEALAEDALKLLSALKGKIVIVVYRPERREEAEELAE
KLRRIGAEVYLLDVEAPDYEKRLEEIADEIISRLSAWSHPQFEK

NL13
MSGMMEIVLIGRDREKLEEVKEYLEELLKERGIENIKIYIGLIDPEKGEGWCLLLALERYDEVPPT
DKTLEEVMKEHRERVKEIEKIEDPERREEALAEDALPIIAALKGKPVIIVYTPEEEEWARELAEKLR
RFGAKVFLLNVEDPDWKKKLEEIVDKIIEMRSAWSHPQFEK

NL14
MSGMMEVVLIGRDREKLEEVKKELEKILKEKGIDNVKIYIGLIDLEEGKGWCLLLALEWYDEVE
PTDKTLEEVMKEHEERVKEIEKIEDPEERDKALAEDALPIIAALKGKFVIIVYTPEDKEFAEELAEK
LRRFGAKVYLLNVKDPNWKEKLEEIADEIIESLSAWSHPQFEK

NL15
MSGMLEVLLIGRDLEALEEVKKYLEEILKKRGVDNVDIFIAHIDLSEGKGWCLKLALEKYEEFPD
TDKTIREANEEHKERVKEIKKIPDPEERDKALAEDAKELIATLKGKFVIVVYRPEDREWAEELAEK
LRKIGAKVYLLNVEEPDWKEKLEEIADEIIASLSAWSHPQFEK

NL16
MSGMLEVILIGRDRKALEEVKEYLEEILKERGVDNVTIFIALIDPERGEGWCLLLALEKYDSVEPT
DKTIEEAIKEHEENVKEIRKIPDPEKREEALAEDALKVLATFKGKVVIVVYTPEDREWAEELAEKL
RKLGARVYLLDVKEPGWRERLEEIAEEIIELLSAWSHPQFEK

NL17
MSGMVEVLLIGWDREKLEEVKERLEKILKERGVDNVTIYIAHLDPSKGEGYCLELALERYDEIPP
TDKTLEEENARHKENVKEIKKIEDPEKRDEALAEDAVPVIAALRGKVVIVVYTPERREEAERLAE
ELRRFGADVFLLNVEDPDYEEKLEEIADKIISKLSAWSHPQFEK

NL18
MSGMIEVVLIGRDLEALEEVKKELEKILKERGVDNVTIFIAHLDPSKGEGYCLKLALERYDEVPD
TDKTIEEANEEHKKNVEEILKIPDPEERERALAEDALKVIATFKGKVVIVVYRPEEREWAEELAEK
LRKMGAKVFLLDVEDPNYKEKLEEIADEIISLLSAWSHPQFEK

NL19

MSGMVEVVLIGRDLEALEKVKKELEEILKKRGVEDVDIFIAHLDPEKGEGWCLYLALEKYEEVP
DTDKTIAEANAEHKENVKEIKKIEDPEKRREALAEDAVKVIATLKGKVVIVVYRPEDREEAEKLA
EKLRKYGARVYLLDVEDPNWREKLEEIADEIISLLSAWSHPQFEK

NL20
MSGMLEIILIGRDKEKLEEVKEELRKLLEKRGITDVDIYIALLDPEKGEGYCLELALEKYDEVPDT
DKTIEEATAEHKENVKEIEKIPDPKERDKALAEDALKVISTFKGKFIIIVYTPEDREYAEWLKEKLE
KIGAKVFLLNYEEPDWKEKLEEIVDEIIASLSAWSHPQFEK

NL21
MSGMLEIILIGWDEKDLEEVKKKLEEILKEKGINDIKIYKALLDPEEGKGYCLRLALERYDEVPET
DKTLEEADEEHRENVKEIIKIEDPEERDKALAEDALELIATFKGKYVIIVYRPEHKKEAEELKKKL
EKLGAKVFLLDVSDPNWKEKLEEIVDEIIADLSAWSHPQFEK

NL22
MSGMPPLYLLLVGPGITEEEIEMAKEAAKEVGLDDLKVYKIRWDPEKGEGYCIELAVEKKAKRIP
TDLTLEEAAERHKEVAKKAKEIEDPEKRDELLVEDMGVVLGAIERVIIIAKYKEHYELYKKLFEDA
GIEAMFFLHPDPSKPFTLEWLIEQLKKAKEFFESAWSHPQFEK

NL23
MSGKPPLYFLLVGPGVTDEEIELAKEALKEVGLEDVEVYAVARNPEKGDGYCLRLAAEKDAEIIPT
DMTLEEAAERHKKVSEEAKKIEDPEEREKLLIEDAGRVIGALKRVVVLAEYPEYKEWFEKLLTKA
GNEYLLFLHPDPSKPLTKEWLIELFKKVKEYFASAWSHPQFEK

NL24
MSGMPPLVILLVGPGITEEEMERVREAAREVGLEDAVVYRVPIDESKGEGYCLKLAAEEDADVSP
TDITLEEAAERHKKVREEAKKIKDPEKRRKLLIEDAGRVIGALRRVLVIAFEEEWYEWYKKLLER
SGVEYMLYLHPDPSKPISDEWLKEILEKAKEYFYSAWSHPQFEK

NL25
MSGMPPLHLLLVGEGITEEELELAREAAKEVGLDDLEVYAVETDPSKGEGYCIELAAEEDRPIEPT
DRSVAELAEEHKKVAEEAKKIEDEEKRKEMLIDDTFRVIGAFRRVIIIADRQEWLDKFKKLFELAG
VEYMLYLMPDPSRPLTREELIKWLKEAKEKFFSAWSHPQFEK

NL26
MSGMPPLYLLLVGPGVTDEEVELARRAAERVGLEDLQVYRIAPEPEKGDGYCLRLWLERKAPRIP
TDLTVDEAAEKHKEVIKEAEKIEDPEERERLLVEDTGRVLGTFDRVIVLAREEEVLEFYKRLFEAA
GIEAYFFLWPDPERPPTLEELEAWLREAQEFFASAWSHPQFEK

NL27

MSGMPPLYLLLVGPGVTEEEIELAREAARRVGLEDLEVYAIAPEPEKGDGYCLDLWLKADAPRIP
TDMTVDEAAERHKKVVEEAKKIKDPKEREKLLIEDTGRVLGTFKRVIVIARRPEDLEYYKELFEK
AGIEYKFFQWRDPERPPTLEELEEWLREAQEYFKSAWSHPQFEK

NL28
MSGMEHVVLIGWDRELLEEVKKELEEILKEKGVDNVEIYEALLDPATGEGACLLLALERYEEFPP
TDKSLEEVIKEHEENVKEIEKIEDPEKRAEALAEDAIPVISALKGRFVIVVYTPERREEAEELAEKL
RRLGMKVYLLDVTDPNWKEKLEEIADEIIESMSAWSHPQFEK

NL29
MSGMPKLTLLLVGPGITDEELELAREAAKEVGLEDLEVFALDRRPEEGDGYCLKLWLEAKAPRIP
TDKTVEEAAEEHKKVIEEAKKIEDEEERNKLLIEDTGRVLGTFERVIVIADTPEYYEFYKELFEYAG
IEAYFFLRPDPSRPLTLEELKEWLKEAQKYFESAWSHPQFEK

NL30
MSGMKTVVLIGWDREDLEYVRKKLEEILKEKGVTDVEIYEALLDPATGEGACLLLALERYDEFP
DTDKTLEEVIEEHKKNVEKIREIKDPEEREKALAEDAIPVISALKGKFVIVVYRPERREEAEELAEK
LEKIGMKVYLLDVEDPDYEEKLREIADEIIASMSAWSHPQFEK

NL31
MSGKPPLYLLLVGPGITDEEIELAREAAEKVGLKDLEVYALEREPEKGDGYCIDLYIKRKAPRVPT
DLTLEEAAERHKKVIEEAKKIKDEEEREELLVRDTGVVFGALKRVIVYAREQEWYDKYKELFDRL
GIEHYFFLHPDPSRPPTLEELIEWLKEAKKYFESAWSHPQFEK

NL32
MSGPPPLVLLLIGPNITEEEIELAREAAREVGLEGLPVYHIDWDPEKGIGYCFQLAAEKPTPVIPTD
ETLEELAERHKETRKEAEKIEDPEKREELLLEDVGRLLGALRRVLLLAFREELLELFRKLLERAGI
EFLAFLHPDPSKPLTKEWLKELLREAQAHFASAWSHPQFEK

NL33
MSGMKYVILIGWRQEDLDLVKEKLEEILKKKGVDNVVIYKSLLDPETGEGACLLLALERYDEVPP
TDKTLEEVIEEHRENRKEIAKIPDPAERARALAEDALPVIAALKGKHVIVVYRPEEEEEARELAEK
LRRFGMKVHLLNVEDPNFEEKLEEIVDKIIEDMSAWSHPQFEK

NL34
MSGMMYVILIGWREEDLDKVKEEMERILKEKGVDNVTIYKSLLDPETGEGACLLLALEKYDEVP
PTDKTLEEVIAEHKENRKEIAKIEDPEERAKALAEDALPVIAALRGKHVIVVYRPEEKEEAEELAE
KLRRFGMKVHLLNVEDPDYLEKLKEIVDEIIASMSAWSHPQFEK

NL35

MSGSPEVILIGFREEDLEEVRRRLEEILAEKGVTDVTIYRALIDPATGEGACLLLALERYDTFPDTD
KTLEEVIAEHRENVKEIRKIPDPEEREKALAEDALPVISALKGKFVIVVYRPERREEAEELAEKLR
KIGAKVYLLDVEDPDYEERLEEIADEIIASKSAWSHPQFEK

NL36
MSGAPEIILIGFREEDLDLVEKKLREILAEKGVTDVTIYKALIDPETGEGACLLLALERYDEFPDTD
KTLEEVIAEHRENVKEIRKIEDPAERERALAEDALPVIAALKGKFVIVVYRPERREEAEELAERLA
RFGARVYLLDVEDPDFEKKLEEIADEIIASLSAWSHPQFEK

NL37
MSGMEEVVLIGRDREDLERVERELREILKERGVEDVTIYRALLDPETGEGACLLLALERYDEFPD
TDKTLEEVIAEHRENVKEIRKIPDPEEREKALAEDALPVISALKGRFVIVVYRPEDREFAEELAEKL
RKIGMRVFLLDVESPDYRKRLEEIADEIIASRSAWSHPQFEK

NL38
MSGMTEVVLIGWREEDLEKVKEELERILKEKGVEDVEIYRALINPETGEGACLLLALEKYDEFPP
TDETLEEVIARHRENVEEIKKIEDPKEREKALAEDALPVISALKGKFVIVVYRPEDRERAEALAEK
LRKIGMKVYLLNVEDPDYRKKLEEIADEIIKSMSAWSHPQFEK

NL39
MSGMEKVVLIGWRREDLEYVKKKLEEILKERGVDNVEIYEALIDPETGEGACLLLALERYEEFPD
TDKTLEEVIEEHRKNVEEIRKIPDPAERERALAEDALPVISALKGKFVIVVYRPEEREEAEELAEKL
RKIGMKVYLLDIEDPDFEEKLREIADEIIASMSAWSHPQFEK

NL40
MSGMMEVVLIGWDREDLERVQKELERILKEKGVDNVTIYRALLDPSTGEGACLLLALERYKEFP
DTDKTLEEVIKEHEENVKKIREIPDPEERAAALAEDALPVISALKGKFVIVVYRPEREEEARELAK
KLEELGMKVFLLNVEDPDYKKKLEEIADEIIASMSAWSHPQFEK

NL41
MSGKTTVILIGWDEEDLKEVEEKLREILKERGVTDVDIYRALINPETGEGACMLLALEKYDEFPD
TDKTLEEVIKEHRENVKEIRKIEDPEKRAEALAEDAIPVISALKGKFVIIVYRPEREEEARELAEKL
RKIGAKVYLLNIEDPDWKEKLEEIADEIIESMSAWSHPQFEK

NL42
MSGMTTVILIGWNREHLDEVKEELERILKERGVTDVDIYKALIDPETGEGACMLLALERYDEFPD
TDKTLEEVIEEHRERVKEIRKIPDPAERAAALAEDAIPVISALKGKFVIIVYTPENEEEARRLAEKLR
RIGAKVYLLNIEDENWKEKLEEIADKIIESMSAWSHPQFEK

NL43

MSGMEEVILIGWEQEDLDEVKEKLEEILKEKGVEDVTIYKALLDPETGEGACLLLALERYDEFPD
TDKSLEEVIKEHKENVKEIKKIPDPEERDKALAEDALPVIAALKGKFVIVVYRPEHREEAEELAEK
LRKVGAKVYLLDITEPGWKEKLEEIADEIIKSMSAWSHPQFEK

NL44
MSGKTEVILIGWREEDLDYVKKKLEEILKEKGVTDVTIYKALIDPETGEGACLLLALERYDEFPPT
DKSLEEVIKEHKENVKEIEKIEDPEKRDEALARDALPVIAALRGKFVIVVYRPERREEAEELAERL
RKVGAKVYLLNIEEEGWEEKLEEIADEIIASRSAWSHPQFEK

NL45
MSGKTEVILIGWDREDLDEVKEYLEEILKEKGVEDVDIYQALIDPETGEGACLLLALERYDEFPDT
EKTLKEVIEEHKKNVEEIKKIEDPEERDRALAEDALPVIAALKGKFVIVVYRPEHREEAEELAERL
RRVGAKVYLLDITEEGWKEKLEEIADEIIESMSAWSHPQFEK

NL46
MSGMLDVILIGRDEKKLEEVEEELRKILKERGIDNVTIYRELIDPETGEGYCMLLALEKYDEVPET
DKTLEEAIEEHKKNVEEIKKIEDPEEREKALAEDALKVIATFKGKFVIIVYSPEDREFAEWLKEKLE
RFGAKVYLLDYEDPNWKEKLEEIADEIIASLSAWSHPQFEK

NL47
MSGSMDVILIGWDEEKLKEVEEKLREILKKRGVDNVTIYRELIDPKTGEGYCMLLALEKYDEVPE
TDKTLEEAIEEHKERVKEIEKIEDPEEREKALAEDALKVIATFKGKFVIIVYTPEEREKAEELAEKL
RKFGAKVYLLDYEDPNWKEKLEEIADEIIASLSAWSHPQFEK

NL48
MSGSLDVILIGKDEEKLEEVKEELERILKERGVDNVTIYRELIDEKTGEGYCMLLALEKYDEVPET
DKTLEEAIEEHRERVKEIEKIEDPAERERALAEDALEVLATFKGKFVIIVYTPEEREFAEWLAEKLR
RFGAKVYLLDIEDPNWREKLEEIADEIIASLSAWSHPQFEK

NL49
MSGMMDVILIGRDREKLEEVKEELERILKERGVDNVTIYIELIDPETGEGYCMLLALEKYDEVPET
DKTLEEAIAEHKENVKEIEKIEDPEEREEALAKDALKVIATFKGKFVIIVYTPEDREFAEWLKEKLE
KFGAKVYLLDIEDPNWKEKLEEIADEIIASLSAWSHPQFEK

NL50
MSGMLDVILIGKDKEKLEEVEKELREILKERGVDNVTIYRELIDPETGEGYCMLLALERYDEVPE
TDKTLEEAIEEHKENVKEIKKIEDPEEREKALAEDALKVLATFKGKFVIIVYTPEDEEFARWLKEK
LERFGAKVYLLNIEDPNWKEKLREIADEIIASLSAWSHPQFEK

NL51

MSGMRPLVLLLVGPGITDEEIELAREAAKEVGLEDLPVYRINIDPEKGEGYCIRLAAERDAPVIPT
DKTLEEIAAEHKENREKARKIEDPEKREELLIEDMGRLIGALDRVILLARDEESLEWWRKLLERA
GVEYVDRLHPDPSRPWTKEELIEFLKEAKEFFFSAWSHPQFEK

NL52
MSGEELRYTLEWRRPDGVEVRLHLTFRRVEEPREVYFGPCAARVLARTAGVDEEEIRERLERILAE
VREELGDRFTIDAIVVLRAPTPEQAEAAAEKFLAALREYDAEHPEIDPQPHKYAVVRRDGIVVIAI
ARGDVPQERLEELAERKLEELLAESAWSHPQFEK

NL53
MSGEKKTYVMEWEKPDGVRVTLLLTFTKVDEPTKVLLGPCAAEVLAKTAGVDKEEIERWLKEIL
KEVEEELGPNFTVDVVVIFDAPEPEQARAAAERFLAALEEYDARRKELDPLPHSYAVIERDGKVFI
AIKKGEVPEEELRRIAEEAYELYKKLSAWSHPQFEK

NL54
MSGETLTKVLRWERPDGVRVTLLLKFTKVDEPTKVLLGPCAARVLAETAGVDEEEIRRWLEEILK
EVEKELGPNFTVDVVVILEAPEEEQARRAAERFLEALRRYDDEHKDWDPRPHSYAVIERDGAVFI
AIKKGDVPEEELRRIAEEAYEYYRTLSAWSHPQFEK

NL55
MSGEELRYRLEWERPDGVRVTLLLKFTKVEEPTEVYLGPCAARVLAETAGVPEEEIREKLEEILEE
VRKRLGPNFTVDVVVIISAPKEEQAREAAERFLKLLEEYDKEHKDILPEPHRYAVIERDGLVFIAIK
RGPVPQEELEKIAEEAYEWYKKLSAWSHPQFEK

NL56
MSGETVEKELRWRRPDGVEVTLKLKFTKVDEPSKIYLGPCAAEVLARLAGVDPEEIREELERILK
EVEKELGPNFTVDVVVVISAPEPEQAERAAEEFLRLLEEYDERHKEELPEPHRYAVVRKDGRVYI
AIKRGDVPEEKLREIAEKVLEEDLKKSAWSHPQFEK

NL57
MSGETVERRLRWERPDGVRVELRLRFTRVEEPTEIYLGPCAARVLAELAGVDPEEIRRELERILEE
VRRELGPNFTVDVVVVIEAPEPEQARAAAERFLELLREYDAENAEELPEPHRYAVVERDGRVFIAI
KRGDTPQERLEEIARREMERLEKESAWSHPQFEK

NL58
MSGETVERELEWERPDGVKVTLKLKFTKVEEPSKIYLGPCAAEVLAELAGVDPKEIEEELKRILE
EVEKELGPNFTVDVVVVIEAPEREQAEEAARRFLRLLEEYDERHKETLPEPHRYAVVEEDGKVYI
AIKRGDVPQEELERIAREVVEEDKKKSAWSHPQFEK

NL59

MSGMEVRKELRWERPDGIRLTLLLRFRKVDEPTKIYLGPCAARVLAALAGVDEEEILRRIEEILKE
VEKELGPNFTVEVVVIIEAPEPEQAREAARRFLELLREYDEEHKEELPEPHRYAVREEDGAVFIAIA
RGDVPLEKLEEWAQKAMEERKKESAWSHPQFEK

NL60
MSGMEVRYELRWRRPDGVEVTLLLTFTKVDEPTKIYLGPCAARVLAKLAGVDEEEIRERIEEILRE
VEKELGPNFTVKVVVVIEAPEEEQARKAAEEFLAALKEYDEEHAEEFPEPHRYAVREEDGRVFIAI
ARGDVPLEELERWAEEVMEEYRKLSAWSHPQFEK

NL61
MSGEERRYRLRWERPDGVRVELVLKFTKVEEPTEIYLGPCAAEVLAKLAGVDPEEIREELERILEE
VREELGPNFTVDVVVVISAPTPEQARAAAREFLELLEEYDKKHKEEFPEPHRYAVVEEDGKVYIAI
KRGDVPEEELRRIAEEVVEEDKKKSAWSHPQFEK

NL62
MSGEEREYRLEWERPDGVRVRLILRFTKVEEPTEIYLGPCAAEVLAALAGVDPEEIRERLEEILEE
VKKELGPNFTVDVVVVIEAPEPEQARAAAERFLALLEEYDERHKEEFPEPHRYAVVEEDGKVYIA
IRRGDVPQEKLEEIARRVVEKDKKESAWSHPQFEK

NL63
MSGEEVKYRLRWERPDGVVVELHLTFTKVEEPTKIYLGPCAAEVLAKLAGVDPEEIRERLEEILK
EVEKRLGPNFTVDVVVVISAPTEEQARKAAEEFLALLEEYDEKHKDEFPEPHRYAVVEENGKVYI
AIKRGDVPEEKLREIAEKVVEEDKKKSAWSHPQFEK

NL64
MSGETIERRLTWERPDGVRVELHLRFTKVEEPTKIYLGPCAAEVLAKLAGVDPREIERRLEEILKE
VEKELGPNFTVEVVVVISAPTEEQARAAAERFLELLEEYDKKHKEEFPEPHRYAIVEEDGKVYIAI
KRGDVPQEELERIARRLVEKDKKESAWSHPQFEK

NL65
MSGETREYVLEWERPDGVRVRLVLRFTKVEEPTEIYLGPCAAEVLAKLAGVDPKEIERRLEEILEE
VREELGPNFTVEVVVVIEAPTEEQARAAAEEFLAALEEYDKEHAEEFPEPHRYAVVEEDGRVYIAI
KRGDVPQEELERIARRVVERDRKESAWSHPQFEK

NL66
MSGKELRYELRWERPDGVVVTLRLTFREVEEPTKIYLGPCAAEVLARLAGVDPKEIEAWLKKILE
EVEKELGPNFTVDVVVVISAPTPEQARAAAEEFLELLREYDERHKERFPEPHRYAVREEDGKVYI
AIRRGDVPQEELERIALEEIERDKKESAWSHPQFEK

NL67

MSGMEKRYELEWERPDGVRLRLLLKFTKVEEPTEIYLGPCAAEVLAELAGVDEEEIREKLEEILK
EVREELGPNFTVDVVVVISAPEREKAEEAAERFLRALEEYDRRHAEELPEPHRYAIVEEDGKVYIA
IRRDDVPEEELERMAREEIERLKKESAWSHPQFEK

NL68
MSGKELYYELRWERPDGVRVTLYLKFTKVEEPTEIYLGPCAARVLAELAGVDEEEIRKKLEEILEE
VREKLGPNFTVEVVVVISAPEPEQARAAAERFLAALEAYDRENAERLPQPHRYAIVEEDGKVYIAI
RRDDVPEEELERFARETIERLKKKSAWSHPQFEK

NL69
MSGETLEYVLEYERPDGVRVRLHLKFTKVDEPSEVILGPCAAEVIAKTAGVPEEEVLKELEKILEE
VRKKLGPNFTVDVVVIIEAPEPEQARAAAEEFLALLRELDARDPERFPEPHKYAVVEEDGKVYIAI
KRGDVTEEELREIAEETLRRDKEESAWSHPQFEK

NL70
MSGEEREYELRWERPDGVRVTLWLRFTKVDEPSEVILGPCAAEVLAALAGVPEEEVLRELERILE
EVRKELGPNFTVDVVVIIEAPEPEQARAAAEEFLERLRALDAADPARFPEPHKYAVVEEDGKVYI
AIKRGDVTEEELKEVAEAALKRDKERSAWSHPQFEK

NL71
MSGMERRYRLRWERPDGVRVELLLKFTKVDEPSEVILGPCAARVLARLAGVPEEEVLAELERILK
EVREELGPNFTVDVVVIISAPEPEQARKAAEEFLALLRELDEEDPERFPEPHKYAVVEEDGKVYIAI
KRGDVTEEELKEVAEEALREYKEESAWSHPQFEK

NL72
MSGKELYYELRWERPDGVVVRLRLKFTKVEEPTEVELGPCAARVLAALAGVPEERILRELEEILE
EVRKKLGPNFTVDVVVVIEAPEPEQARAAAERFLAELRRLDASDPELFPEPHKYAVVEEDGKVYI
AIKRGDVTEEELEKVAREVIERDKKKSAWSHPQFEK

NL73
MSGKEVYYRLSWERPDGVRVELLLKFTKVEEPTEVLLGPCAARVLARLAGVPEEEILAELERILK
EVREKLGPNFTVDVVVRISAPEPEQARAAAERFLAALRELDASDPELFPEPHKYAVVEEDGVVYI
AIKRGDITEEELEKIAREALERYKKESAWSHPQFEK

NL74
MSGMEVYYELEWERPDGVRVSLLLKFRKVDEPTEVELGPCAARVLARTAGVPEEEILKKLEEILK
KVREELGPNFTVDVVVKIEAPEPEQARAAAERLLAALRELDASDPARFPEPHKYAVVEEDGVVYI
AIKRGDVTEEELRKIAEEALERYKKESAWSHPQFEK

NL75

MSGKTKYYELRWRRPDGVEVRLLLTFTKVDEPTEVLLGPCAARVLAELAGVPEEEILRRLEEILK
EVREELGPNFTVDVVVKISAPEPEQARAAAERFLAALRELDASNPELFPEPHKYAVVEEDGKVYI
AIKRGDVTEEELRKVAEKALERDKKRSAWSHPQFEK

NL76
MSGETVRRELRYERPDGVRVSLLLTFTKVDEPSEVLLGPCAARVIAETAGVPEEEVLAELERILEE
VRKELGPNFTVDVVVRIEAPEPEQARAAAERLLAALRELDAENPELFPEPHKYAVIEEDGKVFIAI
KRGDVTEEELREIAEKALEEYKKESAWSHPQFEK

NL77
MSGEKVERELRWERPDGVRVSLYLTFTKVEEPSEVLLGPCAARVIAKTAGVPEEEVLAELERILKE
VEEELGPNFTVDVVVIIEAPEPEQARKAAEKFLAALRELDASNPELFPEPHKYAVIEEDGKVFIAIK
RGDITEEELRKIAEEALKKYKKRSAWSHPQFEK

NL78
MSGKTVRRELRWRRPDGVEVSLHLTFTEVDEPSEVLLGPCAARVIAKTAGVPEEEVLKELERILK
EVEEELGPNFTVDVVVVIRAPEPEQARAAAERLLAALRELDAADPELFPEPHKYAVVEEDGVVYI
AIKRGDVTEEELREIAERELERDKKESAWSHPQFEK

NL79
MSGMEKKYVLRYERPDGVRIELVLKFTKVEEPTEVLLGPCAARVIARLAGVPEEEVLKEIEEILKE
VKEELGPNFTVDVVVVIEAPEPEQARRAAEELLAELRRLDAADPSLFPEPHKYAVVEEDGKVYIAI
KRGDVTEEELLEIAKEVMEEDKKESAWSHPQFEK

NL80
MSGKSVEYELTWERPDGVRVTLYLKFTKVEEPTKVYLGPCAAEVIGRTAGVPPEEIEKEIKKILEE
VEKELGPNFTVEVVVRIEAPEPEQARAAAERFLAALREYDARDPSRDPEPHKYAVIERDGVVYIAI
KRGDITEEELREIAERYMEKYEKKSAWSHPQFEK

NL81
MSGETVEYELTWERPDGVRVTLHLRFTKVDEPTKVYLGPCAARVIAKTAGVPEEEIREEIERILKE
VEEKLGPNFTVDVVVRIEAPEPEQARRAAEEFLKALEEYDKEHPDIYPEPHKYAVIEEDGVVYIAI
KRGDITEEELREIAREYLRKVKAASAWSHPQFEK

NL82
MSGETVKYELTWERPDGVRVTLHLTFTKVDEPTKVYLGPCAARVLARTAGVPEEEIEKKIQEILK
KVEEELGPNFTVDVVVRIEAPEPEQARRAAEELLKALEEYDAEHPEIYPEKHKYAVIERDGVVYIA
IKRGDVTEEELREIAERFLDEYLAESAWSHPQFEK

NL83

MSGMEVEYELTWERPDGVKVSLKLRFRKVEEPTEVYLGPCAARVLAETAGVPEEEIREELERILK
EVREELGPNFTVDVVVVIEAPEEEKAREAAERFLKALEAFDAEDPERDPEPHKYAVVEKEGKVFI
AIKRGDVTEEELKELAEKFAEEYLKRSAWSHPQFEK

NL84
MSGERVEYELTWERPDGVRVTLHLTFTKVDEPTEVYLGPCAARVLARTAGVPEEEIEEELRRILEE
VKKKLGPNFTVDVVVRISAPEPEQARAAAERFLEALREFDAEDPEKDPEPHKYAVVEKEGEVFIAI
KRGDVTEEELEKLAREYAEKYLKRSAWSHPQFEK

NL85
MSGETVKYTLTWERPDGVRVSLHLTFTKVDEPTEVYLGPCAAEVLARTAGVPEEEIREELERILKE
VEEKLGPNFTVDVVVVIEAPEPEQARAAAEEFLEALREFDARDPARDPEPHKYAVVEKDGKVFIA
IKRGDITEEELRELAERFAEEYLKRSAWSHPQFEK

NL86
MSGETVEYELRWRRPDGVEVTLHLKFTKVEEPTEVYLGPCAAEVLAATAGVPREEIEERLKEILE
EVKKELGPNFTVDVVVRIDAPEPEQARAAAERFLAELERFDAEDPEKDPEPHKYAVVEKEGKVFI
AIKRGDVTEEELEKLAREFAEEYLKRSAWSHPQFEK

NL87
MSGAAELLQRALAWLILLELRRRGVTLEEIIAALRKADEIVAAKDPELAEKLRKKHEETIALLRAN
PELETLPLEELERRIEALLDEAVAALKEIRTDEELRAALREAFKQGVCLAIAGYLDEVLAAWDRLV
ERLVEALPEWRELIEEIWREKREELRRLASAWSHPQFEK

NL88
MSGAAEKLKEALAWLILLELRKRGVTLEEIIEALRKADEIVAAKDPELAEKLRKKHEETIALLKAN
PELETMPLEELEKIIDRKLDEAIEALKEIRTDEELREALEKAFEQGVCLAIAGYLDEVLAAWERLV
ERLKEALPEWRELIEELYEEKKEELRKLASAWSHPQFEK

NL89
MSGAAKKLREALSWLIYLELRKRGVTLEELEAALEAFDERVAAEDPELAARLREKHDATLALLR
ENPELAEMPLEKLEELIDKKLDEAIEELKKIKSDEELRKALEEALKEGWCLATAGYLEEVLRAHR
RLVERLKEALPERRELIEEIFARYEKELRERASAWSHPQFEK

NL90
MSGMLKLELVDKETAKKNFEKIIFEYLPKALEIFDKMLEEVYDEMDEEEKKRFEELKEKHKKALE
IIDKIEEILEVEYVENPEYDEIRIKVKIDGKEEELTLLKLYHDPEKREESKEQGGCLLIGLSQLDVTIK
LDNEKMLEFIKRFAEELGIPIEVKKSAWSHPQFEK

NL91

MSGMLKLALVDKEEAKKNFEKIIFEYLPKALKIFDEMLEKVYDEMDEEEKKKFEELKEKHKKAL
EIIDKIEEILDINYIKNPEYDEISIEVKIDGKKEKLTLFKLYHDPSKQEESKEQGGCLLIGLSKLDVTI
KLDNEEMLDFIKRFAEVLGIPIEVERSAWSHPQFEK

NL92
MSGEPPIYLLLVGPGITDEEVELAREAARRVGLEDMEIFKLAREPEKGDGYCLRLAAEKDAEIVPT
DKSLEELAEEHKKVAEEALKIEDPEEREKLLVKDAGRVFGALKRVVVIANYPEYYEWFKKILERA
GVEYLLYLHPDPSRPWTLEELEEILREVKERFLSAWSHPQFEK

NL93
MSGMEPLYLLLVGEGITEEEIELAREAAREVGLDDLEVYRIDTDPSKGHGYCIKLAAEKDAEIKPT
DRSIEELAEEHRRVSEEARKIEDPEKREKMLIEDTGRVLGAFRRVIVIADREEYLEWFRKLFERAGI
EYKLYLRRDPNRPLTREELLEILREAKEYFESAWSHPQFEK

NL94
MSGMPPLYLLLVGEGITEEELELAREAAKEVGLDDLQVYRVETDPSKGEGYCLELAAEEDAEIEP
TDKSIDELAEEHKRVAEEARKIEDPEERRRLLIEDTGRVIGAFRRVIVIADRQEWLDMFRKLFERA
GVEYKLYLRPDPDRPLTREELIEILKEAKEYFFSAWSHPQFEK

NL95
MSGMPPLYILLVGPGVTPEELELVREAAREVGIDDAEVFAVATNPEKGEGYCLRLAAEKDAEIIPT
DETLEEAAERHKKVREEAKKIEDPEKREELLVRDAGRVFGALKRVLVIARYPEWYEELKKLLERA
GVKYMLLLHPDPSRPLTREELIEILRKAKEFFYSAWSHPQFEK

NL96
MSGMTITLTDPFTGRKVTVEIEGEVTLWNLPLFLAQLDRALAEAWGRPELEELARHHEAIYEEIQ
ALKPGEEVYIEIGFRGQLLRFRITPDEITFEEILGPATPERGGCTDPVLAQLPELAAKIEKLIEEWEK
KGKVTIHPEPAEVKVRILERSAWSHPQFEK

NL6 catalytic knockouts

NL6_C49A
KIEIILIGFREEDLEKVKEELEKILKEKGVEDVTIYRSLIDPEKGEGYALLLALERYDEVPETDKTIE
EAVEEHKENVKEIRKIPDPAERDKALAEDAIKVLATWKGKYVIIVYRPEEEEKARWLKEKLERLG
MKVFLLNIEEPNWKEKLEEIADEIIKEH

NL6_H74A
KIEIILIGFREEDLEKVKEELEKILKEKGVEDVTIYRSLIDPEKGEGYCLLLALERYDEVPETDKTIE
EAVEEAKENVKEIRKIPDPAERDKALAEDAIKVLATWKGKYVIIVYRPEEEEKARWLKEKLERLG
MKVFLLNIEEPNWKEKLEEIADEIIKEH

NL6_D97A
KIEIILIGFREEDLEKVKEELEKILKEKGVEDVTIYRSLIDPEKGEGYCLLLALERYDEVPETDKTIE
EAVEEHKENVKEIRKIPDPAERDKALAEAAIKVLATWKGKYVIIVYRPEEEEKARWLKEKLERLG
MKVFLLNIEEPNWKEKLEEIADEIIKEH

NL6_G47P
KIEIILIGFREEDLEKVKEELEKILKEKGVEDVTIYRSLIDPEKGEPYCLLLALERYDEVPETDKTIEE
AVEEHKENVKEIRKIPDPAERDKALAEDAIKVLATWKGKYVIIVYRPEEEEKARWLKEKLERLGM
KVFLLNIEEPNWKEKLEEIADEIIKEH

##### 3.3 Lysine ligases

The 79 lysine-ligase sequences are numbered KL0–KL78, with each enzyme followed by its cognate peptide sequence (KL0AP (acceptor peptide), KL1AP (acceptor peptide), and so on). Peptide entries exclude maltose-binding protein (MBP) and the shared linker but retain any additional N-terminal glycine–serine spacers. Each MBP–peptide fusion has the N-to-C-terminal order MBP–GGSGGSENLYFQSGGSGGS–peptide; the shared linker and MBP sequences are listed after the enzyme–peptide pairs, with MBP last.

KL0
GLEIRPATPEEAAKAVEIGKKAIEANGGHPAPGSCGYIAARVVEAVPGARLALLVGDGEVIHVLADTGLTDA
NGEPLLADPTLVQLVAPKLAALHGVPPITVDQAVAILVAGINAYRDGDMDAAVDAIVAGLEAALGITLTPEER
AFVRRLARELALAPRSTVASRFPEVHREIAAFR

KL0AP (acceptor peptide)
SAAAEARAAKAAAEAASPL

KL1
GISLRPLTDEEARAAYEIGKKAVDANGGRPAPGSCGYLAYRLAERTPGARVVAVVGDGEIIHVITDTGLVDA
DGRPIYADPILVQTVLPYLAKKHGKPIPTADQAVDVLLAGVNALKSGSVEAAAQALVDATEEVLGVTLDEA
ERQFIRDYGVELVFTTEEEYIKNAVSALERLARWL

KL1AP (acceptor peptide)
SDADRLRAAKAGALAATPR

KL2
ALEIRPATPEEAALAVEIGKAAVDAAGGRPAPGSCGYLAAGVADRVPGGRVALVVGDGRIHHVVADTGLTDE
NGNPIVADPALVQYLVPKLAKKHGVPVPTADQAVDILLAGVEAYRDGDRDKAADAVVEAIEKTLGVKLSPE
DREFVRNLTKKVIITTKENYAKNAVAGFDAIAAYS

KL2AP (acceptor peptide)
SAGDALAEAKARAEAAAPS

KL3
SLEVRPLTPEEEAEAVAIGEAAIAANGGRPAPGSCGLIAAKIRERIPGSRLVAVVGDGRILHVLTDVGVTDENG
RPILADSSLGQTALPVLAKKHGQPIPTVDDVVDFVLAAVEAYINGNMELAQQAAVDKAEEILGVKLNEEER
EFLREATTKLVVITAEEAATTKVDDFDMLAAFG

KL3AP (acceptor peptide)
SDADRYAAAKARALAAAPR

KL4

LDLGIRVVLVPREEIPAVVAAGRALAEAVGFPVPTCAEQAALAVERLGDVASLAVILDEDGRPLHVLAWVRG
KKRVLIDAQGGFIVANTQDDPEARRAFASSVLLAAQQLGADVLIIVGPREGVEFFRELAKELGIPVEILGAET
LEEAIDLARELA

KL4AP (acceptor peptide)

STAAAARRAKINGVLRMPP

KL5

NFLIALIATFILAKELAAVADLSDPGAAILVITTSKAYGTLMALGLTEAAEKVRELGLKAVRSGSYPPVRVLTP
EETAAVGALGLALAAAGTGWSCGAIAYRATQIDPDYRAVLVTDEDGNPFHVIAVYRNDFKAVADPIVGQIAL
AAGATPEEVLAIALGTSLGVALVVGVPLELAVRAAQVVAQASLEVEREIARLRA

KL5AP (acceptor peptide)

SGGQLLREAKRVAIEAAGG

KL6

MADYRGKYVATGLGVLVELARRAGVPAEAVEAAVAAFEEAFAAIYELLLRGDVRYISLEEARAYSERWRAF
GEEVAARHSCGYIAALLAGKAGGRVVVIGEDGVPVHVIAEVVDPKTGETIFLDPLAGQGIVPGILKALGRPV
TLEGVAEYTVPAILQEWKEGKLFRLP

KL6AP (acceptor peptide)

SLAARARAAKQRLLDAAVW

KL7

SPFEFGKALLAKKGYAYEVVTDAVPEANALGTLIAGIAIYLGNYKEYSLEEYETWYAEHASEETGGCARRA
YEIAKEYAKKGYTPKVIVLLDKDGKPFHVVAAYLDPKTGITYLVDPLIWQLIGGDPATATPEQIARARRGAEL
GLKIVYDESYPVLKKDLYA

KL7AP (acceptor peptide)

DPLQAAVDAKRVAIAGGFA

KL8

GAADERAEALLAEYKATVARVGETVVANPVAQIAPDVRLGRADPALAFRAAELYLELVDLLGYKAVSRRTG
HGGCLALAALIARHVAEKAGAETAVLGVVGDGEVVHMLAYARLPATGARIVVDPAGVQGSLGLGENPGSA
DVVAGAERLADEAVRALRGAPPGATVALAGIPGDGPGRLSGVIVIPAEEATVEALLEAADIVIEKSKTNGAA
ALGVLGLGIAMAAALALAGRPEEAGPVLEVGARRALEIARS

KL8AP (acceptor peptide)

SLGRLLREAKGAALAALGL

KL9

MFLIVFLTSYLVGKKLARLVDLSDPAARFVVIFTSTLVGALYALGHPEAAEKALELGLKAAASGELPKLRVL
DAEEGRAVGVLGIALGVLGTGMTCGAIAWLIGQLDPRYVVAVVLNSKGEVFHVIAVRDGDLARPVDAIAGQ
IALWAGATPEEVRAIVLGGYLGLAFVLGWPLEEAVHRAELAAEATLRTKELIEAAKK

KL9AP (acceptor peptide)

SGGDLLEEAKRVALEAAGV

KL10

MAAFTAAALAAMAAIGGKGRILSLEEGPEVRAAGVKQGGGCGNIAVDAARWLQETKDLSLLRIIVVEDSN
GNPIHALVAYVEKGSDTLHLLDDIPAVIALGEVPAELRADVRAALHTIQLVALALALEAARDLPITRVRLDAA
VPDRPQSDVNAKLKRIIPGEVLPIADFDDPAALAAALVDSALAAAEAAIR

KL10AP (acceptor peptide)

SAIDAAAAAKRAALERMPR

KL11

SPAEAAQVINRIARERFGLPGGCAALALAVIRGLGVPLVRTAAQDPALLALMERMIADVPQEHIDTIAAWAL
KALGAQPTAAAPVFLYNSAGEPVHVILVAALADGTWGALDFLLLQGGRMPGADDYVSGVADIDTLAAQAG
ELVARHLRALAPPEEADKIGAAERDYVAGRVRAVARDLARVA

KL11AP (acceptor peptide)

GAGDVLARAKAAALRSAPE

KL12

RDALTPEQLALADSVTGRIVPNDLALTRPYAEAGRRVGGHCAPSAVAAQEAFPEVELGLLLDADGRVIHAM
AVWRPAPDRVVIADVFAGVALEVPIPAGQTAADVVQLYGAALLVSALTGDPLPSDATETPAGREVLDYMLST
PEGQAQHRAGVLGRELGAQARAAGATDLRGVAAYVTSTYA

KL12AP (acceptor peptide)

DYAQAARDAKFGAIRAAGA

KL13

LKPLTPEELRKLTEEEYLALAEEIGKRVAEELGAGEAARLLAIAAALMTPAVIVAAYRAHVEGLPLAVAVLSR
GDTPIHALALRTDGTVTEVIDPGGVQVGSELASEAIALSLVEMANYLGVPATLGTPVPLPNGFRVATARIALP
TPERYEAIAALVERHGFDVRLVPAEEAPALARRVADDPSVGGCGALAAAIVLQELGETELAREAIERAGF

KL13AP (acceptor peptide)

SAAALAIELKDEAIRLARA

KL14

MADYRGLLVATGLETLVRLAREAGVPEAAIEAAVAAFEKAFAAIYELVLAGDVREISLEEARAYSEKWLQFG
EEVAARHSCGYIAAELAGRAGGRVLVIGEDGEPVHVIALVTDPRTGEPVLLDPLAGQGIVPGILKAEGRPITL
EGVAEYTVPAIEEELAAGRLFRLP

KL14AP (acceptor peptide)

SRAARARAAKQELLDAAVW

KL15

MNAAARAFLDELAALSPEELLERMVAESLPLAQAVAELAGVKPLDRAEALAAARIGFYAVVDPSVTPEQAA
ALVRELTPTEAAFAVIGANAALIHLRLYRVISVDELVERAAALGFPISDELADRAREAEAAGAALVLLGTPEN
PVHVLAVAYDPALGRYILLDPAAIQVLVPRAARALGLTPPPGGGCAVLALALLVAHHEGIRDPRAFVERVLRL
ASRH

KL15AP (acceptor peptide)

SPVERARAIKEAALEVGPR

KL16

LEVRSDNFVPVADYAAVIADALDELGYSYDPAALAAASALLVKADEVARQMADAGLWRVVSVEEIRKLIAE
LLAAKDPNYHGCANVALAIVEALAGVFDAQPAVLREGDTYYHAVAVVALPGTDDPPVVVDPSLVQLAAAF
AGVDLTDPAQDAEIRRLAAAVLRAILRHNPDLPERYRRVLEAAADNLVAAVTA

KL16AP (acceptor peptide)

SLAAEYLELKNGAIALAGG

KL17

SAVDAAVVINRIAREKFGLPGGCGAIAYIVARRLGVPVVRGPNLDDELWALMEELYNSVPEEVRDTIAQAVL
DAIGVKPAAATPVVYYDSAGRPAHVDLVFKAEDGTWGAVDFLTLQAGLPLGADDVITGLATAEELAKAFAE
RFIAVLRSLASPEAAALVGDDVRTYLAEVTLAALEALEAAV

KL17AP (acceptor peptide)

AAGDALAQAKADALASAPL

KL18

GIRFSGMKLVTGADVSGGRARALGVPDALVDAVREAAARALAHVLQLIRDGHVRPISLEVARLFPLLLSLL
GPFFSVLFSCSYIAAEIAEKYGGEVYEVGEDGEPTHVVAVVKDPKTGEWVVFDPVYTQGLAPLVLKAEGRP
VTLEGVAELSVADIIAEYERGIYSTVP

KL18AP (acceptor peptide)

SLAAAARAAKLAAFLTGFL

KL19

MADFRGLLVVTGLDRLVELAREAGVPAAAIDAAVAAFERAFDAIYALVLRGDVTEISIEEAEAYSTRWLALG
EEEAARHSCGYIAAKLAGQAGGRVLVIGEDGEPVHVIAYVTDPRTGEPVLLDPLAGQGIVPGILKALGRPITL
EGVAEYTVRAILEELAAGRTFRLP

KL19AP (acceptor peptide)

SLAAEARAAKQALLDAAVW

KL20

VSPTVQRALKVFYRLAEIVAAAVGAPAEAVAGSTLLVFDRNGNLLEVLRPDLDAALKAAGLTPEQIAAYEEA
LDAIIGFLQAMRNGESDGILRLARNVQPTDDPFDLLKRAELITNVEQFKNAFKNVKVGDLLFLLDSNGNPIH
VLAVVAADDELVVLLDTLGFQVKIGTDAVVEAQKLAGVPPITYGGGCAGLSRIASKLAKELG

KL20AP (acceptor peptide)

GYSRRAAEAKARALAAGPA

KL21

SIAEAGREALATLLDNGSATLVPVADADLAALRAGALALGADPAAVERQFAARRLVRAVLDAAGVRTIADL
APLFDVEGPALYAGAFTAISALAGLLILQKGAQPGEVWAIGINGEPIHVLAVVAREGDTLITIFDSLVGYLLEV
TRTKPVETDIDIYKISHEEAVAGSNATGGGCGAIAVAAATLFQ

KL21AP (acceptor peptide)

SIAEAFREAKEALLALSQQ

KL22

MAEDVYNAARETVRTVLSGLDPADPGARVGATHFGLAAAGAVAIGGYEAYERILEAARAVAEAGEPVPIRE
LNEEEFGKLAEAALAELRQGVVGNCVALAALMKEVRPDSRVIALLDGNGNIFHIVVSLDDRYLAIADPIIVQ
TVVMAGKSLEHLEVAVEAGVRGVADVLGVPEDLTQQVIDTLVTAGRRYVATLERARA

KL22AP (acceptor peptide)

SAGDLARRAKEAALGATGR

KL23

AREKTLALIEELAKTPLEELVKIFVDNSYPLFVAFARLAGLEPLPKEEVEKIVLEVARMLLDPAYTPEEVAAYV
GSLSPMEAAAANLAVGVALLHAGEYRLLDVDEAIQATVDAGSPASPELVAGYREQYKKGAVAIALGRPDNV
VHVIATMIDPATGEAIVLDPALVQGLVPLAARGLGLTPPPGQICMLLTLAMVVAIGSGETDPEAVARRVLELVA
RP

KL23AP (acceptor peptide)

DVVARAAAAKAAALARAPA

KL24

MAEIVEREARRLGHAIADKLDLSDPGARALVVTMSVMAGVLRALGKEAAAEEVRRLALEAADEGALPTV
RPLNEAEAEAAKEVGIANGVKGTGQICGALALEAHRLDPRLEVWALLDENGNIVHVTGYYDGDPAVPWDP
IAVQVAAAAGASREELVELAAAAFRALSDIVGIPPELADHFARVLAEAAVIVGAEIEAAKK

KL24AP (acceptor peptide)

HGSRYIGPAKEAALSAALA

KL25

MAPLTAEELAAADPADVIGGLAEAADAAADALGGDERDRLAGRVSAIAVLAAIAALPELEARGLTRALVVL
WAGDQPVHAVAVLVHPDGITFIDPIGLQVGSERALQTTVAAAIAGANFAGAPAEPGEDITLPTGEVLPTVRVA
RPPEARIRALRAVLDRAGIRVEVVPLEEAPRLAVLAIQLGFPGSCPGLAAAALYAHLGDDERAQALIDASPR

KL25AP (acceptor peptide)

SPAQLLLEAKRRALALAPR

KL26

LLGDLRVVPVTPEEEPQVVAESERLRRELGLTKPLCGELAVRIARAIGPAARVIRVVDSDGRVIHVLVWVEGK
EKVLVDPQGAVIGAAIEADPEKGGALAAAIARIAEHVGADVLLGLGEHPANPLAVQGIQGVRPDLRVEFFAD
EAALVQAVKDRL

KL26AP (acceptor peptide)

SLSDAFREGKEIGIRAVGY

KL27

MTVTSDQLVEVPDPRAVIRAVLDALGASWPEARLDEAVEAYRKYVEVLRRALDAGLARVLTVDELRALLA
VLRAARHPLYHGCGELALHIVDALAGRYDARPVLVSEGDRLGHVLVAVTIPGTDAPPVVIDPLLAQIALHAA
GVDLWDPAQRPLVRTGLAAFLRAAVEVNPDLPAAQREVLLEAADRLLAQVAA

KL27AP (acceptor peptide)

SLLDELNAHKLAALGAAGA

KL28

SIAELVREAVRAVAERDPSYSEADVELAADATIRALDQILAANPELAKLPAEVLIGMAASIAIIELTAGKPISVT
VLIDGRVVARIGDDGLPDERAEAIARRSPLRYTLVIKIKRSALNPETAIEVDVDGARLLVQGTAENPVHVVAI
VEEEGDTLVLLDNVQAANRLPAGSCTAYAIAALTGRPVGDVAEANALVLEQVKAKA

KL28AP (acceptor peptide)

NGPRAARMAKEAALALAHG

KL29

SSVTSKQYLRFPDPLAVISRALDELGISYDPARLEAAVAAYREFEANLLKALDAGNTRELTLEEFDALLDEYL
AAKPEGFGGCGVLGRAAAHALADDFDVTPVLVEADGVISHVLLRVTPKDLDIPPILLDNSLAQIAGGLAGID
LRDPANAPQGLRILAAILRAILEIDKDLPEWVRQLLEAVVAALERWLAA

KL29AP (acceptor peptide)

SLLDALIAAKDRMLAAAGL

KL30

SLEELIREAVRAVAEKDPRYSEEDVELALEAMMTNWRALVAANPALRDLPVYAQIGLAAAAAIISLLAGEPA
FLAVAVDGKLLAVTGEPILTPEEIEAIARRSPNRYVLVAAVPRSSLSPSRTSVVDVDGARLLVLGTRENPVHVI
AIVAEKGEYVVILDPVQAANGAPAGSCAAEAIAALLDRPIPDVAETNAKVRAALAAQA

KL30AP (acceptor peptide)

DNQNAAALAKEAALRVAGT

KL31

MRPLTPEEIRALDPETYIALAFEIADAVAGAAGADEATRLMARVAALLAPAFLVGIAEAHRQGIELFVSVVW
DGDTPIHAVAVVRDGDRATFYDPGGLQVGDLEASRVVAEAFREMALWLGVPAELGEPFTLPTGARMASVV
VTIPDAERAEALRVLVEKLGLEVRIVPVEEAVALAERDLATPGAGGCGALAAAVVLQAAGDTERARRYVEA
AGF

KL31AP (acceptor peptide)

SAGALALELKAEALRLARE

KL32

WEITAEHLIEVPNPREVIAAALDRLGLAYDEARLDAAVAALREFRVHLREMLRAGHTRPLTLEEFKALLAEY
RAANPEGLGGCGELALWVAEALSGDFDVQPVLVEADGKIAHVLLAVQPKDEDVTPILLDPALAQVAAGLA
GLDLRDPANRPAALALFAAILEALLELEKDLPEEVREILRRVVAALRAALAA

KL32AP (acceptor peptide)

SLIDRAIAAKERMIAAASG

KL33

GEKQERADKLLEEYLKALAVAKQQALDNPIAEIAPGTFIGTIDPDLAARLAGYLLELMDLIGYRAFVRRTGV
GGCLELSALIAEALRELTGAETGVIAIAGGGDVAHVVAWAYLPETGGAVIADPLGVQASLARHPAPRSEDIVA
IARELAEYAAEALRATPAGEITGVQAIPGATPGTLQGAIAIPVSDATLETLDRAIAEAVARLTDPGAKALTALGI
GIAMGVAAALAGRPDLAAPVIERATEIALKLAKE

KL33AP (acceptor peptide)

NLADAIREAKQAGLDALGL

KL34

MDLGIRVVEIPRERIPEVVAAGRRIAEEVGFPVPTCAELAAQAVDELGDVASLVVILDENGQPIHVLAYVRGR
KRVLIDSQGGFIVANTQDDPEARRAFASAVLLAAQQLGADVLIIVGPQEGVDFFRELAKELGIPVEILGAETL
EEAVALARELA

KL34AP (acceptor peptide)

STAEAARRAKIRGVLRMPP

KL35

MTPAEAAALANAELHALGAPVEVTADGNTVTINMPDGEAGCAAGAAAALVAAAGGEVEVNFGAGADAN
TRAQVENGVIGGRAAAGLGPTEAGAVVPVGAPGDIVHVVALVVVNGKKILIDGGPLSGALIAGIAEKAKEIG
ADAILKVTRHVLDPAASAAAINAANAAGLPVRVVGLLEDLTPEQRAALVARLAEVDRLSVSPVVVAEDRDE
AVRLGTAALAEAVA

KL35AP (acceptor peptide)

MDMAALRAEKLAAIAEFNA

KL36

LPPVVQRGLKVFHRANEIVAKAVGKPGTYVATSRLLVFDKNGKLVKTTSPDYDTAWRLAGLSDAQRAAYE
AALDAIVDYLESFTTGASDGTVRIARNVAPDEDPFDALARAATVTDAAGFVNEIANLKVGDLVFLLDEDGE
PVHVLAVVASDDEYTVLFDGLGFQVTIGGEAVVEAQKIAGIPPITYGGGCAGLARIAAKLARELG

KL36AP (acceptor peptide)

SYARRLREAKAAALAAGPA

KL37

SIAEAGRRGLALLLDSGSATLEPVPEADLAALREGALALGADPAAIDRQLAAREVVRAVLEAAGVKTIADL
APLFDVDGPALYAGAYTAISALAGLLILQKGAKPGEVWMIGINGKPIHVLAVVAREGDRLITIFDSLVGYLMH
VERRRPVETDIDIYKISHEEAVAGSNATGGGCGAIAVAAATLFQ

KL37AP (acceptor peptide)

SIAEAFREAKEYLLAAAQQ

KL38

PKPAPGEAIVMTGEEALAYSATGEDPGGCLSASGIAAKRTGGKVLVTAAHAMAEIDVDGKTVVADPTGAQA
AIVPWAPGKPIPQEQIEAIADAAYDAAVEGAWTNPAVGGVIFAVGGLNDVGAAGVPGVIIASASNAAVVLRR
VYGLPAEEVVPAAIALARGFYATLSPEQQAQVDMDAVLERVEEGGRAAAAPG

KL38AP (acceptor peptide)

AAHEIVREAKRAALEAGTA

KL39

MVISGEQLAAALTAVNAALAELVAAARRGAFASRTALFEAIAAAYAKAYADEAGAKTVIVDVDGFPLVVIVG
VDDLTGVRVLTPEELEALLKELGIGLDDPDIGRLCAPISLEVAKAVGNGAKVVIVYDPATGEPVHAVVVAKA
KDGTLVVVDPSQTQYGLPAALYLETGRLPDTETVARRSVELLTKAALAAINGR

KL39AP (acceptor peptide)

SVYARARAAKAAAAAAAAA

KL40

LRPLTAEELAAADPDVVLPGLFAAADAHADALGGDERDRLAARVSAVAVLAAIAALPTLRAQGLRYALVGL
WAGDEPVHAVGVILHPDGITFIDPIGLQIGSELALQATVGAAIDGANLAGAPATPGAPIALPDGRELPTVDVA
LPPEERIAALRALLERAGVSVEVIPLERAPEIARRALELGFPGACPGLAAAALYAHLGDDARARALIDGSPR

KL40AP (acceptor peptide)

SVAERLREAKRRALALAPR

KL41

MMEKAREVMDELAALSEEELLTRLVTENLPLAQAMAKLAGVKPVNLEQALAAARLGYDLVADPRYTPEE
VAEQIKKLSIVEASFAIIGAVAALIRARQYRVITPEELKEIANKLGYPIGDEIVELAKKSVVLGGAVVVIGPEDD
PIHVIAVAFHPVTGKFLVLDPAIFQYLVRYAAPKLGLPVPPNSNCAITALAMMVAVAEGITDPEEFVRRVLELA
SIF

KL41AP (acceptor peptide)

SLAERARAVKEAAREVPPD

KL42

AAASAPVPVSTLPPELQVKVTNAVDTAIALAEKALTKAFGQPVKVSYSDILVADLGDGLYAVFADITIEGAPAI
EDPVEAGLAAMKAEIELAKGGSGGCAGIALAINAYLQQRGYHPFFVLLIDENGNVIHVIVAATKDGNFGVL
DPLPLQMGDVDTFRELFERFLAALDAPNREESLRLSLENAEIIYRRLLELGMPEELARVIRDALQRVVEAAQ
HLPDSL

KL42AP (acceptor peptide)

GFAERFREAKEAALELGSA

KL43

MVLSAEQLAAALTAVHTALAELVAAANAGAYASRRELFTAAAEASAKAYEDEAGAKTVIVDVDGIPLVVIIG
IDDLTGVTVYTPEELRALLDRLGIGLDTPDIGLLCAPISLEVAEALGNGAKVVLITDPATGAPVHSVTVAKAA
DGTLVVIDPGITQYALPAALALETGALPDTLTVARESVRLLTAAGRKAINSL

KL43AP (acceptor peptide)

SVFAAARAAKAAAAAAAAA

KL44

KLKYVALPDAERRAAVAAGAAAAGVAADADAWWAAIRGVDPFVREAAKVSLARGADPVLPGGDERARIM
GELGVAFAQSVVPPELEITPVFGNVHGFLLVRDKRSGGVLVVDPTNAGTQVAPFSCVGATMAIAWGLGVVA
RDGRLTPADIVAMAALAVVFAELAARLFGRSLNERERALIFGASVAGLPKIVSLAYEKLKELR

KL44AP (acceptor peptide)

MDESAVRAAKEAGLAAILG

KL45

DFIAAVREVQALERELVPWSGGCAGVSAIMARATGAPVHFDPDARELIGRIFDHVLAKLSPEAEAALAAAA
AVVVGDREVRDALVVAIGEDGVFAHVLLVLFLTDGTYVGFDVLPFQFGRDVPIESQVFEADSLEALGREMG
RRAARVAAEAAGRTLTEEEIEAVGELAARNLVRAAREAGIIE

KL45AP (acceptor peptide)

SALQALIDAKAAALARAPR

KL46

SPAEAAQVINRIAREVFGLPGGCAALALAVIEGLGVPLVETDNQDPADLDLMRRMISLVPQELIDTIAAWALE
ALGAQPTAAAPVFLYDSRGEPVHVILVAALADGTWGALDFLLLQAGREPGPDDYVAGVADIDTLAVLAGEL
VIRHLRALAPPEAADLLGDETVTYVAERVRAALESLARHA

KL46AP (acceptor peptide)

GAGDALRRAKAAALRTAPS

KL47

MMEKAREVMDELAALSEEELLTRLVTQNLPLAQAMAELAGVKPVNLAEALAAARLGYRLVADPAYTPEQ
VAAEIKKLSIVEASFAIIGANAALIKAKQYRVISPEELKEIANKLGYPIGDEIVKLAEKAVKQGGAVVVLGPAD
DPIHVIAVAFHPVTGEFLVLDPAIFQNLVRLAAPKLGLPVPPNSNCAILALAMMVAVAEGITDPTEFVKRVLEL
AAIF

KL47AP (acceptor peptide)

SLAARARAVKEAAAEVPPD

KL48

SLEELVREAVRAVADRDPSYTEADVELAAEATIEALRQILAANPALRALPPWVLIGMAASIAIIELTAGKPISV
TVLIDGKVVARIGDDGLPDEEAERIARRSPNKYTLVIVIKRSALDPATAIEVDVDGARLLVQGTAEEPVHVVAI
VEERGDTLVLLDNVQAANRLPPGSCTAHAIAALTGRPVGDVAEANAKVLAQVKAQA

KL48AP (acceptor peptide)

SGPRAARMAKEAALALAHG

KL49

RLPKPGEAIRISGEEALELAAKGVDPGGCLRASGLAAKEMGGEVLVTDAHAVAKLDVDGKEVIADPLGAQ
AAYIPYRPGEPIPEEKINKVAEKTIEAAEKAAKEVPEIGGVIYTVGGLEEVGPAGVEGVIIASAANYAHILRRI
YGLPPEQLVEAALELARYLYEQLTPEQKAAVDMERVLERVREGGEAAARPE

KL49AP (acceptor peptide)

GSGSAAHEIVREAKRAALEAGTA

KL50

MLEIVPLTPEEEARAIAIGREAIEAAGGRPAPGSCGYLAALIRDKVPGSRLVALKGKGKIIHIVTDTGLRDAEG
NPILADPLLVQFLVPKLAEKYGKPVPTAEEAADIVLAGVRAYEEGDFDAAAQAVVDAIEKTLGIELDEEERE
FVYKIAREVTITTAKKLPETFPKAAKEIAAYK

KL50AP (acceptor peptide)

SLVEEAAEAKAAAEAAHPL

KL51

MKKFYERAKEKVKEFGGSAEVIPLERAPEVIAAGREYGGGCAQVIYDFLKELEEKEDLVVNKMIVLKDAE
GRPFHVVGVYVKKGSDVLTIVDTMVYTYASGEVPEEEKEKFYKGMEVALVAAIVAAIEAAKDQPIKKIRLN
VAIPDKPQSDPDARFKRLIENEAFPIDEYEDPLELARELIEKALEKAKEGLE

KL51AP (acceptor peptide)

SAMDEARAAKEAALAQMPR

KL52

ALEVRPLTPEEAAEAVAIGRAAVEAMGGKPGPGSCGYLAQLVIDRIPGGRLMAIVGNGGIIHVVADVNLKDA
EGRRILADPLLVQTVVPYLAKKHGKPIPTAEEAADIIMAGVEAYRKGDLDAAAEAVVEAVEKTLGVKLTPEE
REFIRKITKKIVIATEEQLAENFIPVAEEIAKFV

KL52AP (acceptor peptide)

SAAAEARAAKEAALARTPL

KL53

VPDLRGRPLVTGRRRLVEEARELGVPEEEIEAAVRRADRARAEIWAIYRAGRSREISVEEARAIYESYKEKGP
EVAAYHACPAIAARLAEEFGGRVVVIGGDGKPVHVVALAPHPRTGEETLLDSIYAQALAPLVLRGRGEEVTG
EAVVELAVETIREEFESGDTETFP

KL53AP (acceptor peptide)

SLEDEALAAKAEAILHTLL

KL54

PAPLTPEELRAADPEEVYRALGEAADAAADRLGGDERDRAAARTSAIMVLAAIAAIPEMEKQGLERALVIA
WRGDEPVHAIGVVVTPEGLRFIDPLALQIDSEKAAEAQVEAAIAGANYAGAPARRGETIRLPDGRTLPTVEV
ERPPEEYIEAIRAVLDRAGIRVEVRPLEEAPRLAEMARELGYPGACPALAAAALYAWEGDEERAQALLDMV
PR

KL54AP (acceptor peptide)

SVAAQLREDKLRALALAPR

KL55

AVEVEIRPMSDEKVRAALRRIEELGGVKIPEVELTPEQRAAAEAAIRAFIEEGMPLVRRAVDEALAAGTLAP
GRALIARDENGEPVHAFLVEEVDREAGRVVLEDFLPWQALRTPVAGLCLLLAAAAGGAGLTPEQAAEYER
VLPKYLEGYKRPEIDVEKLLEGLSRAARALA

KL55AP (acceptor peptide)

SLGARARALKEEALRIGGR

KL56

MTPAEKLAIKVLEALQELLREAAGRPGNYAGLNRLYVFDKDGRLVRVLEPNREQAKKDSGLSEEEYEAVEK
AQEETRRFLESFLDGTAKGKIRLRRGVKPDEPPEEALARAEEVTSPEQARRAIAEAEVGDLIFLLDENGEPLH
VLGVVARDDEVVVLFDGLGAQAVLGREGVRRLRELAGIPLLPEDGHCYDLARIAAHLAQKLG

KL56AP (acceptor peptide)

GSGSSYARRLREAKAAALAAGPA

KL57

PAPLTPEELRAADPAEVYAALKEAAEKAAEKLGGDERDKKAGRTSAIMVLAAIAAIPELEEQGLERALVVA
WEGDRPIHAIGVILRPDGIIFIDPLGLQVDSEKAKEVIVEAAIAGAEYAGAPAKRGKTIKLPNGEELPTVEVK
MPPREHIEAIRAVLRRAGIEVRVVPLERAPELAAQARELGFPGACPALAAAAIYALEGDQERAQALLDLVPR

KL57AP (acceptor peptide)

GSGSSVAAQLREDKLRALALAPR

KL58

MAERAYRVAREVAERLAAEADLSRPGARWTVWYTATAAGVLAALGYDAAAAAVVAAGEAAVRDGDYPEI
RPLTPEEAAAARELGRRLWLAGTGFGCAAIAGEVAELDPRYTLYGVLDREGRIWHVIAVPDGDLSRPVDPIA
IQIAARAGLSDEEIREVVRAAIRGVGEVLGEDPAEVEERAELLVEAGLRVAAERRAAGL

KL58AP (acceptor peptide)

AGADLLREAKRVALGAAGA

KL59

SMERVYEAARETVEAYLAGLDPEDPQARIGATFYGLAAGAAVAIGGEEAYRKILENAKAVAAEGKPIPTRVL
NEEEFKKMGEEALERLKKGEKGDCVAIAWIIKELYPDAEVVALLDGDGKIFHVVVALDPEYRAIADPILIQTI
LAAGKSLEHLEAAIDAAVRGVADVLGVPEEQARLAAEALKEAARRYAEILAAAEA

KL59AP (acceptor peptide)

SAGDLAERAKEAALAAAGI

KL60

PVPPPGEVVRLTPEQARELGKTPLDPGGCLAISARVAEALGGWVIADAVHAVAAYEHNGRTIIVDPFPLQVEYGLRPDTGEGREEKIRRAAERAIEAARAQADKPIPAGRVGFVVPPLDDIPVEQIPKILRFLGMHLAHVLRHLFGASPEEAVEAARAVVRGAFEQLPEEKKGKIDLERLVREVEEEVRAWAERP

KL60AP (acceptor peptide)

DFVERLAQLKREALAAAGG

KL61

AEAEKRAEELVEEIKEERAALGKEAVDNPLFPIAPGARVGRIDLERALEVAEKFLELLELIGYREAVRRIGAGGCLELSALIARELEELTGAEVGVIAVVGNGEVAHVFAWARLPATGGTVIVDPLGLQAALARHPAPDAETVRELARQAAELAAEALRRTPPGEVVGVAPIPGPEPGRLQGAIAVPVSEASVEKVKELIEEAVARLDDPGAKALTALAVGLAIGVAAAWKGRPDIAEPAIRYAVKKAEEYAKE

KL61AP (acceptor peptide)

GSGSNLADAIREAKQAGLDALGL

KL62

DAAEAEAEELFARYEELAAKLGKEAVENPVAEIAPNVRLGRIDLERALEAAELFLRMVDLLGYREVARRLGHGGCLALSALIARRVAERTGAETGVLGVVLDGEVYHTLAWARLPETGARVVVDPLGVQAAMALDPGATAAQFVAGAEQLADAAAEALRAARPGETVAVAGVPGKRAGELADVITVPASEATMERFRREAERIRALAESPAEAAAAVLGLGLAMGAALALEGRPEEAAPIVAEAAREAERLARE

KL62AP (acceptor peptide)

GSGSSLGRLLREAKGAALAALGL

KL63

MLHVRPLTPEEEAEARAYGEAIAAANGGRVTGESCALIAIRARERYPGAEVLALYDGDEILHVLLGLPIKDEEGRPLAVDPAFAQFVLPALARVVGRPAPTIEEAVDLVLAIARAYEEGGREAAAEALVRKMEEILGAPIPAEPREKVREAAREYAIVPVEEAPRATLKIAELANEIA

KL63AP (acceptor peptide)

SLADRIREAKAAAGRAMAR

KL64

SPLEEARAELAKYGIRYRSTVPDFPEAVELGARMLAQAAREGKLKILSIEEAEKVYEKYKDKVKGNCFEKAREIAEKYAAEGKEAGVAILFDAKGVPFHALGAYRDPADGQWELVDPLGEQLLGGPYEEATPEQIAAAKEAAEKGLKAFEAGLKEAEAEKAAA

KL64AP (acceptor peptide)

AAADAAARAKEAALGLGPL

KL65

SLLERLVELAKKGYTCLTASMYYIQTGKPIKQSEITPEDVEKIYAQWSPELKEAAQAVFDFAEKHGLEGVHVLAVKEYDPETNVITVYDPLTGNTLTLDLNDPAAALATIAEDWKKATDDPEALAALDKAIEEAPKLYAKSGVVEVNPNVKAEVHALAREPVEDPLDPRAEDFLVLFSVNENGKWKPLFLVILRRPK

KL65AP (acceptor peptide)

SAAEAIRRAKDIIIREQLP

KL66

MGEFRGMRLVTGEEYIVEMAREAGVPEERIEAVLEAARKARADVYKMLNAGKYREISVEEARAYAEKYRKYGEEEAARYACALIAAKLAERLGGRVLLVGENGKPVHVLAVAPDPRTGEEVVLDGMFGQYLAPLILAARGEPVTLEKVTEITVEAILKALEEGKYEEVP

KL66AP (acceptor peptide)

SLLERARAAKEAAFELSRP

KL67

VTVTAEQLARAIAAAARRMAEIAQRAARGEYKSIEEAMREAAEAVLEAYREHAGLEGEIIDVDGIPVAILRG
ASGFEGVERLSPEEAEAAVREWGIPPDHPDFRTLCLPIVARLAKELGRGAEVLVLRDPATGEPVHALLVFPLA
NGELALFDPLPFQAVYPAAYKAETGRWPTPEEVVDTAVEAAKKAIEEFLKEH

KL67AP (acceptor peptide)

SPFAEARREKARAVAAAFE

KL68

GFIKVGKVKIENKKTGTSGEADVMLAPAAEAEKVMKEIVEYARKRIEKGDRSVTCGLLTEGAFEKAKEKGA
KKVAPYLLFDSNGKPFHVGVIFEMPDGKLYLYDPLPLQELAVKEGKFSVEELRELGKKYFDELAKEIKEGEK
IEFEGEYLKATYELDPEKIREALA

KL68AP (acceptor peptide)

ASSDTAAQAKADALRIMGI

KL69

KLTVRPLTPEEEAKAIAIGKEAVEKKGGLPAPGACGYVASLMVEKLPGGRLMAIVANGKIVHVIAETGLTDE
NGRPLVLDPVAVQVLIPMLAKKYGKPAITAEQGADILVNAAKTYATESREAAAERFVEDVEKLLGIELPEEER
ELLKRLATEWFLTTAQGAADRFIELYRFLSKFK

KL69AP (acceptor peptide)

SAAEEARRAKAAALANTPA

KL70

PAPLTPEELRNADPAQVYRDLAAAADAAADRLGGDERDRNAGRTSAIMVLAAIAAIPQMEAQGLERALVIA
WDGDKPIHAIGVELTPEGIRFIDPLGLQIDSEKAKEVQVEAAIAGARYAGAPARRGETIRLPDGRTLPTVEVE
RPPEEYIEAIRAVLRRAGIEVRVHPLEEAPELARKARELGYPGACPALAAAALYALEGDEERAQALLDLVPE

KL70AP (acceptor peptide)

GSGSGSSVAAQLREDKLRALALAPR

KL71

MEITAEQLAEALRKAMAEIARLAADAYRGRYKSARELFEEGARAVAEAYREATGADVRIVDVDGIPLVVITG
VTALRGVRVLTPEEARELLEKLGITLDDPDFGVLCAPISVEVAERLGGGARVVLLEDPATGEYVHAFAVARAP
DGTLVIVDPVQVQFLLPAAIYIETGEKPDAATVGRRAVELLVEAGRKALAEA

KL71AP (acceptor peptide)

GSGSSVFAAARAAKAAAAAAAAA

KL72

MRPLTPEEIKALDPEEYRRLADEIATRIAREAGADEATERMARVGALITPAILYGIAEAKERGIERFVVVVWD
GDRPVHAVAVEIHGDRARIYDPGGLQVGGLEESEVIARAIGEMAKFLGVEAEPGEPYRLPTGAEIATVEVEIP
DKELAEAIKALVEKLGMEVRIVPAEEAPALADEDLARPGSGGCGALAAAVVLDRAGDVEAARAYVERAGY

KL72AP (acceptor peptide)

GSGSSAGALALELKAEALRLARE

KL73

PLPAPGEAIRISGREALELAKRGVDPGGCLRASGLVAERTGGRVIVTDAHAYAEVDVGGKEVIADPLGAQAA
YIPWEPGKPIPEEKIEKVAEATLKAAEEAAETIPPIGGVVFTIGGLEEVGPAGVEGVIEASAVDFALILRRVYGV
PPEEAVPAAVEFARRQIERLSPEQRAAVDEERLLERVRRGAEEAAAPR

KL73AP (acceptor peptide)

GSGSGSAAHEIVREAKRAALEAGTA

KL74
EDIERVRELGERVAEAAEIELEPIPDEEMDKIKEKAIELSGNPEAIDRQVETGKKNREVIESRGVKNIKDALKL
LDGDEKERNTGAQLILSAAAAVKAAEIGAKPGDVLLIEGGGGPIHVVAVSHRPAPGAMRIFDVFPGAYVTAR
VTKPLESDITVRRISKEEAIAGDREYGGGCGALAAAAAELGF

KL74AP (acceptor peptide)
GAAVALRAFKDRALAAAAP

KL75
MTLEEALALAGTPLGCLPAALVSALTGKRLPQFGDPAEIEAAAATLDPEEAQRYLDALDYVLEKYNLEPVH
VLALIDYDPATGEIRLYDPLTGEEFTVGADDGKAVALEFAERLKEKAPSEKAKEAAEAAVEKIKGLKNMKPI
EFEYKGLKITVIPLVPKDAPDDTPVEEAELVVIRITPKDDPGRGLVVIARPRPK

KL75AP (acceptor peptide)
MAGPALREAKAAFLRDHPR

KL76
GINFAGMRLVTGEREARAEALAAGIPEARVDEILAAARRAEEEIIALMRAGYVREISYEEAREAARAFEALG
PEVAALHACGAIAAELAEKYGGKVYLIGKPGAPVHVVAVAPDPKTGEPVVLDALFTQALAPEILRARGEPV
TLEAVTAMAVEWIERQWAAGDYTTIP

KL76AP (acceptor peptide)
SLAEAARAAKQRALDLARS

KL77
MREEAEKVMRELQAMSPEELLRRLVEQNLPYAQAAAELAGVKPVNREEALAAARLGYEIVTDPSYTDEEI
REIIEGMSIVEAGFATVGALAAMIKAGRYRVISAEELQEKAAAFGYPIGDALVERAKEAEKKGAAVVVLGDP
DDPIHVIAVAKHPKTGEWLVLDPAVFQQLLPYIAPKLGVPVPDPNNCAAIALAMMVAVAEGITDPREFVERV
VRLAAEA

KL77AP (acceptor peptide)
SLAERARAVKDAALEVPPS

KL78
MPDFRGLRVVTGLRRLVERAREAGVPQEAIDAMVAAFEKAFAAIYALVLAGHVREISIEEARAASEKWLKY
GEEVAAMHACGYIAAELAREYGGRVLVIGRDGGPVHVVALVEDPKTGEPVILDPLAGQGLAPAILKAEGRPI
TLEGVAELTIEAIREEWEAGEYRRLP

KL78AP (acceptor peptide)
SLAAEARAAKQALLDAAIW

MBP_linker
GGSGGSENLYFQSGGSGGS

MBP
MKIEEGKLVIWINGDKGYNGLAEVGKKFEKDTGIKVTVEHPDKLEEKFPQVAATGDGPDIIFWAHDRFGGY
AQSGLLAEITPDKAFQDKLYPFTWDAVRYNGKLIAYPIAVEALSLIYNKDLLPNPPKTWEEIPALDKELKAKG
KSALMFNLQEPYFTWPLIAADGGYAFKYENGKYDIKDVGVDNAGAKAGLTFLVDLIKNKHMNADTDYSIA
EAAFNKGETAMTINGPWAWSNIDTSKVNYGVTVLPTFKGQPSKPFVGVLSAGINAASPNKELAKEFLENYL
LTDEGLEAVNKDKPLGAVALKSYEEELAKDPRIAATMENAQKGEIMPNIPQMSAFWYAVRTAVINAASGRQT
VDEALKDAQTNS

#### 3.4 Lenacapavir ligases

LL1
MSGLGRVTDEQAETLGRRLGRAIADVLAPHLRPGDHVTIFDPVSEEALADPGSAGARFARAFREELRRELE
RRTGATVTERLVPQRLPPDYRDHPFIGHAIAVGIGEPPALDIPPMRGGCEEVARAIADELARAGVPAYVVEGY
VDQEAYRRGTPVEELPVQARIVERRGSHHHHHH

LL2
MSGLRPTSVEDGKIIGRRVGRAIAKELAKHVKPGTNLTIFDPVSEEALRDRNSPEGAFVEGFREGLVEELKKE
TGANVNIVDIPQILPPDAEEWPDIGHALVVAIGEPPPLDIPPVVGGCEQLARQVLEELRRRGIDSWLIEAGVD
RRDYEEGKPVEELRTFAEIKEHSGSHHHHHH

LL3
MSGGGGLPGRTREDAERARAHVRRAAEEIERAARRLGLRAEARGFVGLGEDPVGGTREAAAWLREAARD
GRRGIVVDPAPEVVLRGETETPLLRRAREALDEAARAAREAGGALVAVAQPLTRDEVQRAWDTAGPWAGH
VSVAVALEGADEAVETFVRRLVRAGLEAVQHVRETGRRGCHDVTFLVGRLAATLEPASDEEIAAAEAAGEV
RIYRGSHHHHHH

LL4
MSGNLGYNIPTDLVLEAIKRAFEEGIKKLKKRPLAVIDPVLDLNELEKTLDPETFAELKELDEKHRELLRDIA
KEYDAEIIPVPQIITKQDVDNLDPNDPGGHAWVITDDLDDARKAAEETEPKIGNCLPIAKEVADKIKRGETLL
CKGRAYPNGKTEYEVEILKSGSHHHHHH

LL5
MSGPTGRGVPIERLLEAHRRATEEGLRRLEGPRRAIIDPRIDLERVERTARPEDLEFIRELEERHNALIETIARR
HGVEVIYVPQVITEDQLLRTDPDAPGGHAWLLTDDVDAARRAAERVEPIEGGCEETALAVADEIEEGRVWL
VEGRAYPDGETRTRVELVKAGSHHHHHH

LL6
MSGPTGRGVPTERLLEAHRRATEEGLRRLEGPRRAIIDPRIDLEQVERTARPEDLEFVRELQERHDALIETIAR
RHGVEVIYVPQVITEDQLLRTDPDAPGGHAWLITDDVDAARRAAERVEPVEGGCEEIALAVADEIEEGRVW
LVEGRAYPDGELRTRVELVKAGSHHHHHH

LL7
MSGPGGLGVEPELVLEANERALREGLRLLSGEPRAIVDPVLDLDRVEREAPPDVLARLRELQERHHRLLREI
AEREGLEIIPVPQYLTREQIESTPPGQDGGHAWVITDDPDAARRAAETTPPLDGDCETLSLQVARQISRGRVV
LCQGIARPDGRLEARAEVLRPGSHHHHHH

LL8
MSGNAALNVPTDLLLEAHRRALEEGLKLIKRRPLALIDPRLDLPRIEREADPETLARVKEIQEQHDQLIERIA
REHDIELINVPQTLTKEQIENTPPDQPGGHVWLITDDVEDARRAAEKTPTITGGCEQICEEVARELRRGTVLR
CRGRAYPDGEIRYEVKVLKEGSHHHHHH

LL9
MSGHAALNVPTDLLLEAHRRALEEGLKLIKRRPLALIDPRLDLPRIEREADPETLARVKQLEEEHNRLIERIA
KEHDIELILVPQTLTKEQLENTPPDQPGGHVWLITDDVEDARRAAEKTPTITGGCERTCEEVAKELRRGTVL
RCRGRAYPDGEIRYEVVVLKEGSHHHHHH

LL10
MSGTLGRNVPEELVLEAQRRALEAGLRRLKRRPGAIIDPRLDLDEVRRTLPPEDFRRVRRLDERVRELHERL
AEEHGVELIPVPQRVTREAWERTPPDRPGGHAVVVTDDVEGAREAARRVEPREGECERIARQVAEEVREGE
VLLARGEARPDGELRAEVTELKRGSHHHHHH

LL11
MSGRNGYNIPIEIVLEAIRRAFEEGIKKIKKKPLTVIDPVLDLEELEKVLNEEDFNIIKELDEKVRELLTNIAQE
HNAEIIPVPQIITKEDIENLNPNDPGGHAWVITDDLEDARKAAEETNPKISNCLNIARQVAEKIKKGTVVLCE
GVAFPNGETGYNCEILKSGSHHHHHH

LL12
MSGNAGLHVPLDHLLEAHRRALEEGLRLLHRRPLALIDPRLDLPYLERTCDPETLARIKEIQTEHDRLVERIA
REHGVELIRVPQTLTREQIENTPPGQPGGHSWLLTDDAEDARRAAEQVEPVNGGCERICEQVARLLRRGTVL
LCRGEAHPDGELRHEVRVLKQGSHHHHHH

LL13
MSGPLGVGVPTELLLEAVRRAIEEALRRIPERPLAIVDPLLDRDRLRQELPPELAEEILERDDAHRRLLEEICR
EHDVELIEATQVITRDMVETTPRGMRGGHALLVGDDVDALRRAARRVEPVEGGCLEMARRIAEEIRSGRV
WLAEGAARPDGELETRVELLRPGSHHHHHH

LL14
MSGRAGLNVPTDLLLAALRRATLEGLERLRRRPLAIVDPQIDLEEVERTATPEQLRRVREINTQHTELLRQLA
REHDLEIIPVPQTLTREDLEHLGPDDPGGHALVITDDVEAARRAAEETPPLNGNCLRLAREVAERIRRGEVLL
CEGRAYPDGRLEARVELLRRGSHHHHHH

LL15
MSGPVPRGVPEELVLEALRRALEEGVRRLRRRPLAIVDPVIDLDRVEREASPEDLAWLRRLERAYRELLERF
CREHDMELIPVPQIITREQIDAVPEDAPGGHALVVTDDVDGARRAAEEVPPIEGGCRRISLAIAERIERGEVLL
VEGVAEPDGTLSVRVTLLRRGSHHHHHH

LL16
MSGHTPLNVPIDRLLEAHKRALEKGLKKIKKRPLTLIDPRLDLDKLKKELPPDIYKLVERYNNEHEKLLKDIA
KKNNIKIVYVPQILTKERIERTPPDDPGGHSWLITDDPDEAEKAAKKIPTMEGGCDRICEKVAEEIKKGTVFR
CEGRAYPDGTLESKVTLLKKGSHHHHHH

LL17
MSGRAGLGVPEEHLLEAHRRALEAGLARLPRRPLAIVDPRLDLDEVARVLPPEELARLRRLNDRHRALVRE
HARAADAEVVEVPQVWTRHMLAALPPDEPGGHAWLVTDDVDAARRAAERVPPVEGGCERVARAVADELR
DAPVLLCRGRAYPDGSTRAEVETVRPGSHHHHHH

LL18
MSGPARGGAPRETLLEAVRRALEEGLERLPRRPLAIVDPELDLDRVRRELPPDLAAEIIRFSEEHRGLVEEIAR
EHDAEVIPVPQVVTEEEWLSLPPDRPGGHAVVITDDVEGARRAARRTEPVLGGCEEAALRVADEIRSGTVLL
AEGEAHPDGRLRARVTELRGGSHHHHHH

LL19
MSGRIGIGVPTDLVIEANLRALEEAVARLPHRPLTMVDPVLDEDRVRRELPPEEADEIIERARTFHDRLREHA
RRHDVEIVDVPQVLTREQYERTPPGAPGGHAWVVTDDVEGARRAARVTPTVNGGCETLARRVAAEIPRGT
VLLAEGAATPDGELRTRVTELRQGSHHHHHH

LL20
MSGPARGGAPRELLLEAVRRALEEGLERLPRRPLAIVDPELDLDRVRRELPPDLAAEIIRFSEEHRQLVEQIAR
EHDAEVIPVPQVVTREEWLSLPPDRPGGHAVVITDDVEGARRAARRTEPVLGGCEEAALRVADEIRSGRVLL
AEGEAHPDGRLRARVTELRGGSHHHHHH

LL21
MSGRIPLHVDPEYVLEAQRRAIEEGIRRLNAPRLTIVDPQIDWDEVERTLEGEELRYVREIDERHRQLLREIA
REHNVEIIHVPQRLTRKDVENTPPEYPGGHALVITDDIEGARRAAEEVPPIEGGCEECARQVARRIERGRVLL
CSGRADRFGRTEVEVELLRQGSHHHHHH

LL22
MSGKLGLHVPIEHVLEALRRAFEEGLERLHGPRLALVDPRLDLESVERELRGEALERIRELNRRHRELLEHL
AREHGVEIVDVPQRVNREKIETIRPEEPGGHAWVLTDDVEAARRAAERVRPQEGGCEELALQVAEEIRRGN
VWLCEGRAWPDGQTESRVTLLRRGSHHHHHH

LL23
MSGAGIYGPLTEHVRKQIHALYQAIVESEGIPTPEIIKKKLKEHGIDISSAYTHVKEVKTGEEVIEEAKETARE
NYKRGIVIGHSVLVVLEDGSGILASSPLGSLEALRKALEKAKEQGKKIKAIVDPIAIPEGVTDPEKIEKLKEQK
KEQLEGIAKEFGLSDVKIIPVIQETRSIEEILREIEKNGFYLGDCAETIAALLELLEGGDISRVKNEKIREEIRRA
ARELRGSHHHHHH

LL24
MSGAGIVGPLTPKQKATLEAVHDALLENEGLATPEQIKEKLEEKGIDISEAKPYVKEVNTAEEVIEFSNEVAN
ELAKMGIPIGHNTLVKLEDGSYIIGASPFGGLEATDKAIEEAVEQGKKPKAIVDPVAVPKGETDEEKIEEMKE
KKKEELKEIAKKHGLEDTEIIPVLQELPSLEEILAEIEKNGKYLGTCSLVLAGLVELLENGKITLDLPPKVEKD
VKEAAKKLSGSHHHHHH

LL25
MSGGPIPEPTARALGGEETLERTERALDRVAEALEARHPGLRVLRVEIEPGYGHVYATVDPALFPRARFDVVL
VGQTPGPGWDEYIATPEGRRCLRLVEEEMLAAARAGWIVIDPVPRPLLEERARALARRGYRVLGVDGPEHP
ALCEVLTIGTAAGLLGLEPEEVLALAPDLPPEDREALAEAVRIGLLVRELLLADALPTPAQIEELARRHGLPPG
SVRVYLPAGSHHHHHH

LL26
MSGGAETLTPEQARRMAEALVREALRRGVRRVLLGRQRVELRSPEAQQAAQAHFDTLLAAREALERAEED
GPFYLLDFTPIDHPELLQPEVEYTGEVEVPAGATGHSVLITDVPPIPVEAGRGGCTAAALESVRRTEERLGRP
VPAVLVNAAPTREHAELGRQLAERLMREYQLPNVEVITLVVGSHHHHHH

LL27

MSGGGEHLTPEEARRMYEAIVEEALRRGVGRIELGRQEVGLTSDEARAGAQAGLDVLLRAEEELRRARTRS
PLTLLDFTPVDHPRLLPVEHTYRGRLDVPPGVRGHAVLLTDADVPVVTGGRGGCIGLCLEAARLARERLGR
PVEATLLVGAPTREEADLGVQLARELARERGLDHVGVEPVVVGSHHHHHH

LL28
MSGGPGILDNPELREKIAEDTIKLIEKLVEKYKPKYLVLVPQYEGDKEENIKLLKELNIPPGHAIGILTNKKKG
IIKPIYPGKEEEILEEIENYLKEINDEIIAIIDPVIVTGEDRKYGELITNEMEKLAKKYNVKLIILKGPEVKPKTG
GCVGTVIGTLEGLDYEIIADENLLKARKYADEDTIEAGKRAGKELKKELEGSHHHHHH

LL29
MSGGGGAYVPTREIAERLGAEPRHPEALAIARRVAELTGADVLGAYRIPGPERPAFVILVQQVHPEHGPTSDR
AVLLEFARRNPRPGHAWVIGRARIDGRDVPLLIDPTLEDLGEPLVRGRRPLPGNCVDVSREVAERLERAGLL
AGPGLVVEVGVDEPGEAAVRVNERALAAVLRGLRDGDIREAIRRVERAVREAGGWVRIRPAEEDLSRGSHH
HHHH

LL30
MSGGGGAYVPTREIAERLGAERDHPEALAVAEAIAEVTGATVLGAYRIPGPERPGFVVLVQQRHPEFGPTSD
RERHLELARRNPRPGHAWVIGRARVGGRAVPLLIDPTLEDLGPPLVRGRREFPGDCREVSREVAERAHRAGI
LVEPPLVVEVGADEPGEAAVEVNERALAAVLRGLQHGDVEEAIRRAERAVREAGGWMEVRPAEEDLSVGS
HHHHHH

LL31
MSGSEAEERALEILREFFARMNERTGLDIIPVPLYPGRETMEEITDKILALIREHEANIYIDALVPLSKYRAVGE
DQLADEIIAMKRRILAEIEKRGGYVILIRQDNDLDTLRAGLARIRAGKGVHVVVLIRADESLVERLKAVGAA
IAALPDAFTEPLDLPIDSPRLIVLPGIKGVGNCIILIPLYLALALDLADLFREYAARLLEVLPELREAVDIALAVV
DRLGSHHHHHH

LL32
MSGDPIEEALKYTPEEYEAAVRAAILAVPGVTGILDLDLSDEELFTPGLREVTAPDGQRIGLLRVPQANTRLL
TPEQIAELRKQVVTADVPAGLLVRLDTFDPSLKDRDTLLRRLQLVRLRTVPVGDLAPGEFRIIVVPGHIIALFR
PAADVTLRLPTAVGRGCPLVIILALTQLGHDPVEVAQAVIDALRAAGYSDAEIRALFEQALRDNADIPELAAQ
ARLVLALLERGSHHHHHH

LL33
MSGDPIAEALRYTPEEYREATRAAILAVPGVTGILDLDLSDEELFTPGLREVTAPDGQRIGLLRVPQANTRLL
TPEQIAELRKQTVTADVPAGLLVVLDTFDPSLRDRDTLLRELQLVRLRTVPVGDLAPGEYRIIAVPGHILALF
RPAADATLTLPTAVGRGCPLVIILALTQLGHDPVAVAQAVIDALRAAGYSDEEIRALFEQALRDNADIPELAAQ
AELVLALLEEGSHHHHHH

LL34
MSGDPIAEALKYTPEEYRKATIAAILAVPGITGIYELDLSEEELFTPGLREVTAPDGQRIGLLRVPQANTRLLS
PEEIAELRKQRVVAKVPAGLLVRLDTFDPSLRDRDTLLRQLQLVRLRTVPIGDLAPGEFRIIPVPGHIIAVFRPA
KDTTLDLPTAVGHGCPLVIALALTQAGHDPVAVARAIIDALRAAGYSDEEIRAMFEQALRDNADIPELAAQA
ELILALLEEGSHHHHHH

LL35
MSGDPIAEALKYTPEEYREATIAAILSVPGITGIVELDLSEEELLTPGLRVVTAPDGTRIGLLRVPQANTRLLTP
EQIAELRKQRVRADIPAGLLVRLDTFDPTLRDRDTLLRQLTLERLRTVPIGDLAPGEFRIIPVPGHIIALFRPAK
DTTLDLPTAVGRGCPLVIALALVQMGKDPEEVAKAIIDAWRRAGYSDEEIRALFEQALADNADIPELAAQAE
LILALLKKGSHHHHHH

LL36
MSGSPIAEALRYTPEEYRAATVAAILAVPGITGIAELDLSDEELFRPGLTVVTLPDGTRIGLLRVPQANTRLMT
PEEIAALRKQRVRADVPAGLLVYLDTFDPSLLDRDTLLRRLQLVRLRTVPLGDLAPGEYRIIPVPGHIIAVFRP
AKDTTLDLPTAVGRGCPLVIALALTQMGHDPEEVARAIIDALRAAGYSDEEIRALFEQALADNKDIPELAAQ
ARLILELLKEGSHHHHHH

LL37
MSGDPVEEALKYTPEEYRKATIEAILAVPGITGIYISDLSDEELFTPGLRVITLPDGSKIGLLRVPQANTRLMTP
EEIAELRKQTVTADVPAGLLVRLDTFDPSLRDTDTLLKQLQLERLRTIPIGDLKPGEYRIIPVPGHIIAVFRPAK
DTTLRLPTAVGRGCPLVIALALTQLGHDPVEVARAIIDALKAAGYSDEQIRAAFEQALEDNKDIPELAAVAEL
ILELLKEGSHHHHHH

LL38
MSGDPIEEALRYTPEEYRAAVRAAILAQPGVTGVLDLDLSEEELFRPGLTVVTAPDGQRVGLLRVPQADTRL
LTPEQIAELRKQVVTADVPAGLLVRLDTFNPSLLGRDTLLRQLQLERLRTVPVGDLAPGEYRIIAVPGHIIALF
RPAADTTLTLPTAVGRGCPLVIALALVQLGHDPTEVARAIIDALRAAGYSDEQIRALFEQALRDNADIPELAA
QAELILALLEEGSHHHHHH

LL39
MSGDPIAEALKYTPEEYRAATEAAILAVPGITGIHRLDLSEDELFTPGLREITLPDGRRVGLLRVPQANTRLLT
PEEIAELRKQTVTAAVPAGLLVRLDTFDPTLKTRDAHLHALQLERLATVPVGDLAPGAFRIIPVPGHIIAVFRP
AADTTLTLPTAVGRGCPLVIALALTQLGHDPVEVARAIIDALRAAGYSDEEIRAAFEQALADNADIPELAAQA
ELILALLAEGSHHHHHH

LL40
MSGDPIAEALKYTPEEYRAATIAAILAVPGITGIHILDLSEEELFTPGLRIITLPDGRRVGLLRVPQANTRLLTPE
EIAELRKQTVVADVPAGLLVRLDTFDPTLATRDAHLKALQLERLRTVPIGDLAPGEFRIIPVPGHIIAVFRPAA
DTTLTLPTAVGRGCPLVIALALTQLGHDPVAVARAIIDAMRAAGYSDEQIRAAFEKALEDNKDIPELAAQAEL
ILALLDEGSHHHHHH

LL41
MSGDPIAEALKYTPEEYRAATTEAILAVPGVTGVVILDLSEEELFQPGLRVITAPDGQRIGLLRVPQANTRRLT
PEEIAELRKQRVRADIPAGLLVRLDTFDPSLRDEDTLLRELQLERLRTVPVGDLAPGEFRIIPVPGHIIALFRPA
ADTTLDLPTAVGRGCPLVILLALTQLGHDPVAVARAVIDALRAAGYSDAEIRAAFEQARADNADIPELAAQA
DLVLELLEEGSHHHHHH

LL42
MSGDPIEEALKYTPEEYRAAVEAAVLAVPGITGILEVDLSEEELFTPGLRVFTAPNGQRVGLLTVPQANTRLL
TPEEIAELRKQTVTADIPAGLLVRLDTFDPTLLTKDPKLRALQLVRLRTIPIGDLAPGEYRIIALPGHIIAVFRPA

ADTTLRLPTAVGLGCPLTLALALTQMGHDPVEVARAIIEAMRAAGYSDEEIRAAFEQALADNADIPELAAVA
RLILALLEEGSHHHHHH

LL43
MSGMRPRVDPETVAAAVDRIVAETGLDPLIALTGPDAELLAATEAALRARGHRVLADTDPLALAERVAAEL
GPAARPRLVLVPQVMTPERLARLRALVDAGEIGGHVFLAAAGDRVAIGVVLGLDTDVLTTLVVSLAKLGLE
VADPVPPSLIPAFVEAAKAAGVPVEVVPAPPAPLDATCVNYVIAIGRLMTEEGLPLAEAARRANEVLPTIPPIG
SHHHHHH

LL44
MSGARPAVDPETVADAVDLIVEQTGLDPLIALTGPDAALLAATAAALRARGHRVLADTDPLALADRVAAAL
GPAERPRLVLVPQVMTPERLAEMKARVAAGEIGGHVFLAAAGDRVAIGVVLGLDDDVLITLVVALARRGLV
VADPVPPSLIPRFLAAAAAAGVPVEVVPAPPAPLDATCVNYVIAIGRLMTEEGLPLDEAARRANEVLPTIPPIG
SHHHHHH

LL45
MSGPRPAVDPERVAAAVDLIVERTGLRPLVALTGPDAALLAATEAALRARGHEVLADTDPVALADRVAARLG
KAEAPRLVLVPQVMTPERLAAMRAAVAAGEIGGHVYLAAAGDRVAIGVVLGLDVEVLTTLVVELARRGLV
VADPVPPSLIPAFLAAAAAAGVPVEVVPAPPAPQDATCVNWVIAIGRLMTEEGLPLAEAARRAPEVLPEIPPI
GSHHHHHH

LL46
MSGPRPRVDPARVADAVDLIVRETGLRPLVALTGPDAELLAATEAALRARGHEVLADTDPLALADRVAARL
GPAADPRLVLVPQVMTPEKLARMQAAVAAGEIGGHVFLAAAGDRVAIGVVLGLDNDVLITLVVALARRGLV
VADPIPPSLIPAFLAAAAAAGVPVEVVPAPPEPLDATCVNYVIAIGRLMTEEGLPLAEAARRANEVLPEIPEIG
SHHHHHH

LL47
MSGARPAVDPQRVADAVDLIVERTGLDPLIALTGPDAALLAATEAALRARGHEVLADTDPLALARRVAARL
GPAAAPRLVLVPQIMTPERLAELQARVAAGEIGGHVFLAAAGDRVAIGVVLGLDVDVLTTLVVELARLGLVV
ADPVPPSLIPAFVAAAAAAGVPVEVVPAPPAPMDATCVNYVIAIGRLMTEEGLPLDEAARRANEVLPEIPPIG
SHHHHHH

LL48
MSGPRPAVDPATVAAAVDLIVERTGLRPLIALTGPDAALLAATEAALRARGHEVLADTDPLALAERVAARLG
KAENPRLVLVPQVMTPERLARMRALVAAGEIGGHVYLAAAGDRVAIGVVLGLDVDVLITLVVALAKLGLV
VADPVPPSLIPRFLAAAAAAGVPVEVVPAPPAPMDATCVNYVIAIGRLMTEEGLPLAEAARRANEVLPTIPPI
GSHHHHHH

LL49
MSGLRRLGEEGAVELGRRMGRAAARVLAPRLPRGSRLTIFDPVSEEALEDPESIEARVNRAIRGELRRELEER
TGATVTVVDVPQRLPEDRYDDPDIGHALVVGVGEPPPLDIEPVVGGCEELARRVAEELARAGIPAWLVQVG
VDREAFERGVPVEELPVWAEVRLEAGSHHHHHH

LL50

MSGGPFFNPERQRELLRLLKVRHIDEEEVERLREEYRLSRLQAEVLAAIDRIIEEHREELIEDIVEILRRLREEY
GWEELYIVDTSPDVERALRGEPAHPVARIVLEAARRFERRYGVPVRIIPVPQGMPPEEYAETILEHPIEVRAIII
LPGHAVGIGRPIREEASPERRERVREILERVYPRLEQRARETGLGLCVEDVLLLAAASALLEAGDDPDEIRRS
VRERLERLPIIIPDATRRALGEEYVEFVRRRVEEAREEVERLVLEELEGSHHHHHH

LL51
MSGGAGDPRNTREDVERARREVTAGAERIAAAARALGLEARGHPVVGVGRDLAAATREAARLAREAAAR
GERAIVIDPAPRVVLEGDTSTPLTREARAALDEAAEAVREAGGALVLAAQPLTRADLERAEADAGPLAGHA
SLVLALRGPAEAVEALVRRLVRAADRAVEHVERTGQPGCDLIVHEIGRLAATEEPASDEEVARLEAEGRVRV
VRGSHHHHHH

LL52
MSGEEEIEKALEILEEFCQEINEKTGLKIIPIPLHFGKKTIDEITQEILNKIKEEEANIVIDAIIPLKKLREVGYEE
YAKQIEEMYRKLNEYVKEMGGKLILVEQDNNLEEIKIGLENIKQGNGGHVIFLVKGPEDLVQKLLEVGEKL
KKLPYASEEPLDLPIKDENVIILPGIKGIGNCIVTIPLILSLALNLEDLFREFCEKLYEVIPQLKETIENALKILEE
LGSHHHHHH

LL53
MSGGARGPTDALLRALARIAAAARAAGLTAVLVYLVAPGETAPPAPPGTDLVLRRPVPLPPEERFSPAFAAEI
GRVLAEVAARLPGAEAVVDPVPPPLVPAVRAAFDASGAPARLVPVPQPEDGLTPAGIDEAIRAGAAAAAALP
DLAPRALVSLALVRGGRIVGGHFSLVLGPGPPPGPDAALAVVDALIDASIDVGRLGGCAETCAAVLLALALG
RDADRAAGELERRLRAARRAAGLPAGSHHHHHH

LL54
MSGRVPLGVPLDLVLEAHRRALEEGLRRIPRRPLAIVDPRIDLDEVERTAPPEILAQVRELQERHEELLRTIAE
EHGVEIIPVPQRLTEEQLRNTDPEEPGGHAWLVTDDVEGARRAAERVPEEEGGCERIARQVAEEIRRGEVLL
CEGRAYPDGELETRVELLRRGSHHHHHH

LL55
MSGAGIRAPLTEKQRQILQAVHEALEKNEGCATGEDIREELRRRGIDISDAVTRVRRVKTAQEVIDLLEETAD
ELRQKGLPVGHAALVVLPDGSGILGASSFGALEALEKAIEEAKKEGMKPVAIVDPVAVPKGLTDEERIEQIRQ
KKTERLQQIAKKHGLEDVKIIPVIQETPSLEEVLEEIERNGRYDGYCGRIIAALLELLEGKPITYDLPERVRQDI
RRAAEELRGSHHHHHH

LL56
MSGGASSAHPLIGRPAPEAGREVGEELRRLLGPARGPTDDPLAREGARLGAELGLAVARALGGGAAAAVAA
RLPHAVLVTALALDADGRLVGGHSWLLSRRHLLDPGASPLNLEAARRLGLRPVLVPQHFPPEAGEDLARLR
AAGVGPREDPRVERRPELARTGPGTCLGVIERLLREGHPELGLRPGTPVLGVATRVGALPGAPPAAVEAARA
AAEAVRAALLAPGSHHHHHH

LL57
MSGYERAEARIEETAERLRALGFRVILLPRLPDAPRRIREALAEGRRLGPDGWVVGHVALALLPERRTAILLV
GLGVVRDPRDPEQVRAALRRAHENLLEAMEHAVRRYRPEVLIDPTPLAFRERIEIAREHGVLLVLVQQRLPP
ELRELLREVFGDRVDPELSRRLLARPGAVLVVGPDGRVVAERDAGPLLEPVPPDAPLCDLLCVAVARGLHEH
GFPTDPEGLAAAVVRGLEELPGLLREAAGSHHHHHH

LL58

MSGPPAEQATADHLGGRETLERASRELDRVAEALQRRHPGLRIIRVRVGPRLGHVWATIDPALLPRARFDVV
LVEQEPGPGWEEAAQTPEFRRLRERILEEMLAEARAGRIVIDPVPREDLDARVDALRRAGFRVLAVRGPRHP
ANCEVTTIGVAAGLLGIPPEEVLALAPDLPPELRAALAEAVRIGRLVRDLLLADAPPSPAEVERLAAEHGLPP
GSVRVFEPAGSHHHHHH

LL59

MSGPADVAEAVRRGARLWAAGRRRAVLDLPSGLTARVTIAPARHGAGVTVELLDADGRFLGGHAAAVARR
PLRLAQVLDPADLEATREAASPAEAWRRTLARLPARLAAFDAGDRAALRGPLVERLPNGLWLIDSGAGPLG
PEPPDVVGPPGSCAHLTLGVLAALDRHGPGATPEQVAAAGRDLLRETLAAPDPAAAFLDLAERHGLRLPHP
RTRANVRVMLELLRADPGAARLYIDGIRAVPPGSHHHHHH

LL60

MSGLALDPETVGRAVARLALERGVDTIVASALANPFLLEGIREELRRRGSPIPLLVRDLTDPENRERVLEELR
GRRRVLVIGSQPIPGLNSPEERERALRTTRNTVETGLLALAPGGTLIIDCAAVPVPRERLEEAARDLARRLGIE
DLRGHAFAEVRGLRPSPEAAERTARLTVLATRLALEQDPPLGACEFVALGVAIGAVVHAGGEAEIVSIVRAD
ADIERVVELIRRAFGLLRDHPELRELARRVRLGEPEVERIEAGEPLTLVRIPVRRRPPGSHHHHHH

LL61

MSGGGERLTPEQARRMGEAIVREALRRGVGRIRLGRQRITLPSPEARAGAEAHTNTLLEAQERLERERHDG
PYTLIDFTPVEHPRLLPPEHRWEGRLEVPPGVRGHAVLLSDAPVPVVEAGPGGCLAACLEAARLAQERLGR
PVRATLICAAPTREDAERARRLAERLARERGLPNVDVVALVVGSHHHHHH

LL62

MSGGTPAGERGLEVAVRHYPVLRRLYEASPEERAELVTELLREAGAEVEEIVEVHGRPVIVARLDGRRVVDP
LQHPEDTPEENLAKGRRAAELAEPGDVIDPNIGTPLGLAIVRELLRSDAPIAVGGIDAEGEITVRTARTAEEAA
ALIRELLDRGWAGHIALRTDAREVAHAGGGGCGEICIGAAAALGARVYIVRTGDPVVDERAERVAALAREA
GAEVVFLDGSHHHHHH

LL63

MSGAHTQRAMQFTPEEYEAETRKAILETKGITGIETLDLSDEQLLQPGLYDYTSPDGTKIGLLKVPQADTDL
LSPEQIEELRKQRIKTHVPKGTLVFIDAYNPLLRGKETKLRQLTIANLRTVPVGELKPGEYRIIPVPGHLIGVLR
PREDVELDLPTAIGAGCHLIIALALQQLGFNPEEIAREIIRAEKRAGLSDEEIKRDLQRAYDDNKDIPELKRIA
ETMLRLLEEGSHHHHHH

LL64

MSGGGVATRELALAVAARRGETLGEYLRREAERARRAVERAEALGKRGVLVPQPLVPIPEENPLDALGHTF
AVVADDPLAVARRAVDTPDTRPAAIMDALAGDPAVRALVDPVPEDLEAIDDDLAPGGCEELAVERGAELVR
AGEAQGLVIVTVLYEARRLTVEEAVERVARRAEELGLRVVDRLVNEERRIGAVALVPAGSHHHHHH

LL65

MSGGAGVIENKEEREKITEDIIKLIEKLVEKYKPKYLVLVPQYEGNIEKTKELLKKINAEPGHSIAILTNKKKA
LIFPVTPGKTEEILEQIEKELKELNAEIEAIVDPIPVTGELKEEGERIKKRMEELAKKYNVKLIELKGPEEPAPV
GSCAPLVIGTAEGAGYEIIAPDVIIKAYKLAPKEIIEEGIKAGKELKELLEGSHHHHHH

LL66
MSGGPVRPAPEEVAAAVARLVEEAGLDPVVALTGRDRAQLERLAAALRRRGLEVLADTDPEALAARVAAH
LGPAAEPRLVLVPQRLDPEALARARARVARGEVGGHLGLAAAGDRVAVLVTVGLDPGTHLRLAVALAARGL
TVVDPVPPELVPALRAAAARAGVPLEEVPAPPAPHPGSCVQVCLAVARLHRREGLPVGEAARRAPEVLPEVP
PVGSHHHHHH

LL67
MSGGGGAYVPHRRVLEELGAEPRHPEALRVARAVAEETGLDVLGAYRIPGPRRPALVVLVEQEHPVFGPTSD
REVHEELARRSPRPGHAWVVGRIRIDGRDYPLLIDPTLERLGPPLVRGERREVGGCRETARRVAEALERAGL
LAEEPLLVEVGVDEPGPRAVEVHERALRALLEGLARGDYREAVRRAVEAVREAGGWARVRPALEPPEAGSH
HHHHH
